# The architecture of protein synthesis in the developing neocortex at near-atomic resolution reveals Ebp1-mediated neuronal proteostasis at the 60S tunnel exit

**DOI:** 10.1101/2020.02.08.939488

**Authors:** Matthew L. Kraushar, Ferdinand Krupp, Paul Turko, Mateusz C. Ambrozkiewicz, Thiemo Sprink, Koshi Imami, Carlos H. Vieira-Vieira, Theres Schaub, Dermot Harnett, Agnieszka Münster-Wandowski, Jörg Bürger, Ulrike Zinnall, Ekaterina Borisova, Hiroshi Yamamoto, Mladen-Roko Rasin, Dieter Beule, Markus Landthaler, Thorsten Mielke, Victor Tarabykin, Imre Vida, Matthias Selbach, Christian M.T. Spahn

## Abstract

Protein synthesis must be finely tuned in the nervous system, as it represents an essential feature of neurodevelopmental gene expression, and dominant pathology in neurological disease. However, the architecture of ribosomal complexes in the developing mammalian brain has not been analyzed at high resolution. This study investigates the architecture of ribosomes *ex vivo* from the embryonic and perinatal mouse neocortex, revealing Ebp1 as a 60S peptide tunnel exit binding factor at near-atomic resolution by multiparticle cryo-electron microscopy. The impact of Ebp1 on the neuronal proteome was analyzed by pSILAC and BONCAT coupled mass spectrometry, implicating Ebp1 in neurite outgrowth proteostasis, with *in vivo* embryonic Ebp1 knockdown resulting in dysregulation of neurite outgrowth. Our findings reveal Ebp1 as a central component of neocortical protein synthesis, and the 60S peptide tunnel exit as a focal point of gene expression control in the molecular specification of neuronal morphology.

**Graphical abstract:** 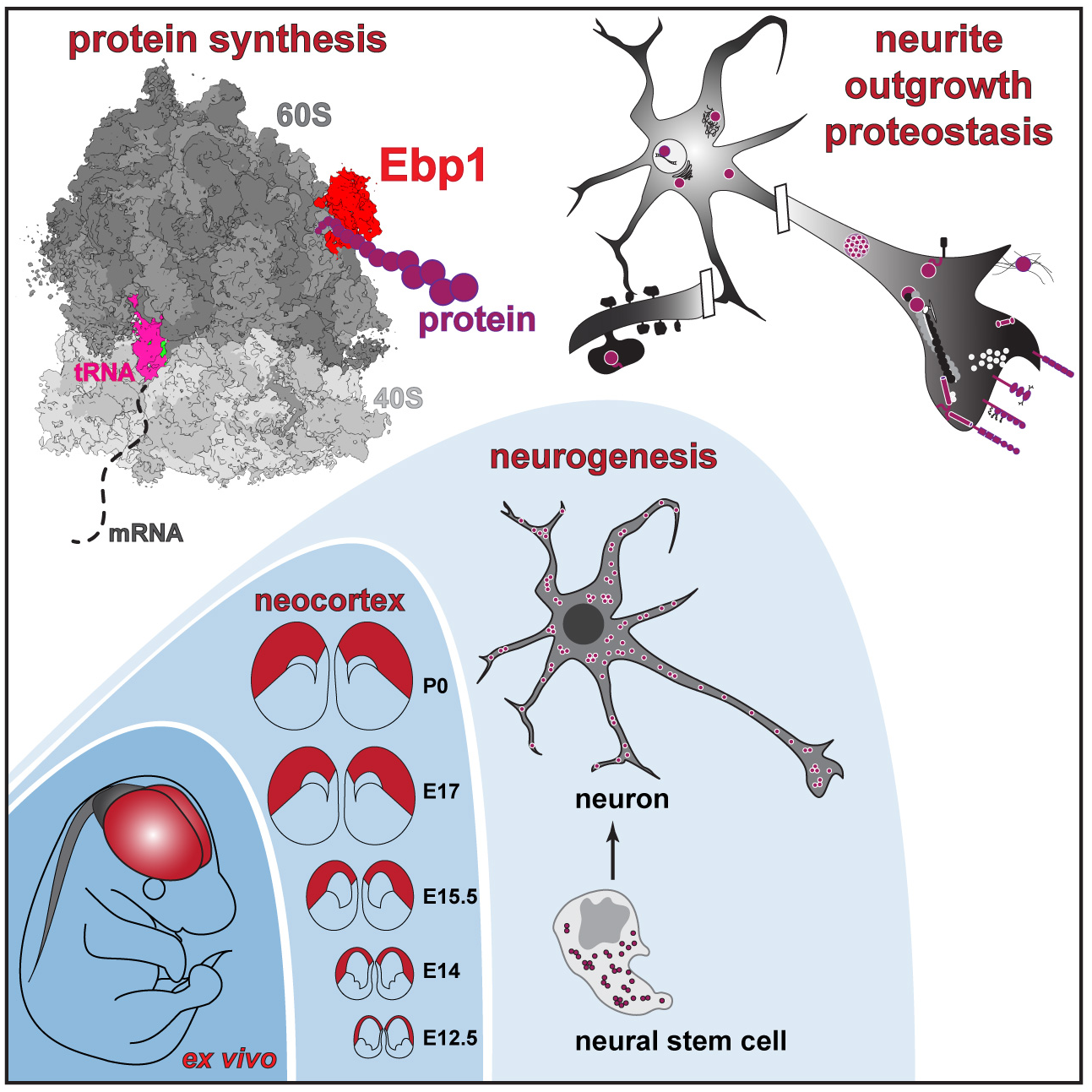

## INTRODUCTION

Proteostasis, the fine-tuned balance of protein homeostasis, is fundamental in establishing the molecular landscape of the nervous system. The demand for spatially targeted and precisely timed protein synthesis is exceptionally high in mammalian nervous system development, where neurogenesis relies on spatiotemporal gene expression, driving amorphous neural stem cells to generate intricately branched neuronal architecture (Hanus and Schuman, 2013; Hipp et al., 2019; Holt and Schuman, 2013; Jayaraj et al., 2019; Jung et al., 2014; Sossin and Costa-Mattioli, 2018). This is particularly true in the evolutionarily advanced mammalian neocortex, the central neuronal circuit of complex cognition in the brain (Rakic, 2009). Concordantly, the nervous system is uniquely susceptible to abnormal proteostasis, a major driver of neurodevelopmental and neurodegenerative disease (Bosco et al., 2011; Kapur and Ackerman, 2018; Kapur et al., 2017; Sossin and Costa-Mattioli, 2018). How proteostasis is achieved, therefore, stands as a crucial question towards understanding neurogenesis in the neocortex.

The neurogenic phase of stem cell maturation in neocortical development follows a general trajectory conserved across mammalian species (DeBoer et al., 2013; Kwan et al., 2012; Molyneaux et al., 2007) (**Figure 1A**). Neural stem cells (NSCs) lining the lateral cortical ventricular zone initially undergo symmetric divisions to expand a pool of cells forming the basis of the cortical plate. NSC divisions then transition to asymmetric with newly born neurons migrating superficially, ultimately forming a layered cortical plate composed of structurally and functionally distinct neurons. Predominantly subcortically projecting lower layer neurons are born first, followed by intracortically projecting upper layer neurons born second. In mice, lower layer neocortical neurons are born at approximately embryonic day 12.5 (E12.5), with the switch to upper layer neurogenesis at E15.5. By postnatal day 0 (P0) neurogenesis is largely complete, with ongoing stem cell divisions yielding cells of the glial lineage. The distinct functional connectivity of neurons in different neocortical layers emerges from the architecture of their projections, where the refinement and targeting of dendritic inputs and axonal outputs pattern neocortical circuits (Harris and Shepherd, 2015). The elaboration of intricate neuronal projections requires proteostasis of the neurite outgrowth and synaptic proteome (Hanus and Schuman, 2013; Jung et al., 2012), a fine-tuned balance of proteins like cell adhesion molecules that establish neuronal connectivity (de Wit and Ghosh, 2016).

**Figure 1.**
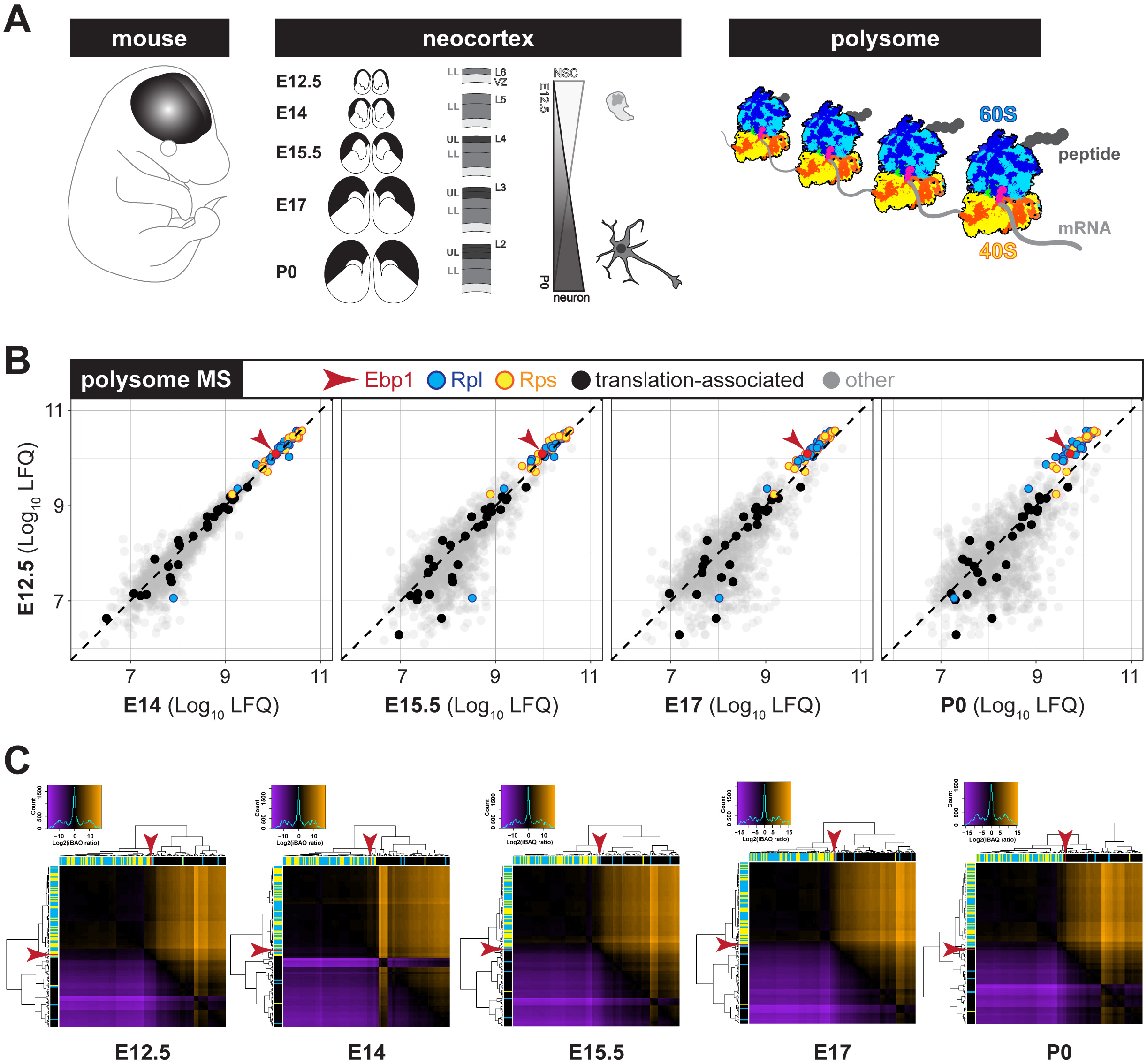
Ebp1 is nearly stoichiometric to ribosomal proteins in translating ribosomes during neocortical development. **(A)** Schematic of the experimental system to measure the architecture of active protein synthesis (polysomal ribosomes) from the *ex vivo* neocortex across embryonic (E12.5, E14, E15.5, E17) and early postnatal (P0) neurogenesis. See text for details. **(B)** MS analysis of neocortical polysomal complexes across development. Scatter plots compare early neurogenesis E12.5 vs. each subsequent stage, demonstrating the enrichment of Ebp1 (red arrow) among ribosomal proteins (RPs) of the large (Rpl, blue) and small (Rps, yellow) subunits, in contrast to other translation-associated proteins (black). **(C)** Stoichiometry cluster heat maps quantifying the differential enrichment of each RP (Rpl, blue; Rps, yellow), translation-associated protein (black), and Ebp1 (red arrow) per developmental stage. Expression of adjacent proteins on the x-axis is shown as higher (orange), lower (purple), or similar (black) relative to each protein on the y-axis. Legend and histogram at top left for each stage. See also **Figures S1-3**.

Analysis of the molecular landscape in the developing neocortex has largely focused on transcriptional regulation (Lein et al., 2017; Silbereis et al., 2016), with the neocortical transcriptome coming into focus recently at the single-cell level (scRNAseq) (Nowakowski et al., 2017; Telley et al., 2019; Yuzwa et al., 2017). However, the functional output of gene expression is protein, and bridging the neocortical transcriptome to proteome is the current challenge. The ribosome is the gatekeeper of proteostasis, poised at the final essential step of gene expression as the macromolecular hub of protein synthesis. Mounting evidence positions the ribosome in a dynamic and executive role at the crossroads of cellular proliferation, differentiation, and disease (Kraushar et al., 2016; Mills and Green, 2017; Shi and Barna, 2015; Xue and Barna, 2012); however, the architecture of ribosomal complexes and proteostasis control in neocortical development remain unknown.

In this study, we analyze the molecular architecture of native ribosome complexes from the *ex vivo* mammalian neocortex during developmental neurogenesis at near-atomic resolution. With a combination of mass spectrometry, biochemistry, and multiparticle cryo-electron microscopy, we reveal that that ErbB3-binding protein 1 (Ebp1) participates in high occupancy binding to the 60S subunit of both non-translating and translating ribosomes through high affinity electrostatic interactions with the peptide tunnel exit site in the embryonic and perinatal neocortex. Ebp1’s role in nervous system development and specific function in protein synthesis is unknown. Ebp1 enrichment scales directly with ribosome levels and is cell type-specific: dominantly expressed in early-born NSCs, compared to later-born NSCs and post-mitotic neurons – in contrast to other exit tunnel cofactors. Ebp1•ribosome interaction occurs in the cytoplasm of NSCs in the neocortical ventricular zone at early embryonic stages when ribosomal complex levels are highest, and persists in post-mitotic neurons of the expanding cortical plate as steady state ribosome levels decline. With a combination of pulsed stable isotope labeling by amino acids in cell culture (pSILAC) and bioorthogonal noncanonical amino acid tagging (BONCAT) coupled mass spectrometry, we show that Ebp1 maintains neuronal proteostasis, particularly impacting the synthesis of cell adhesion, synaptogenic, and neurite outgrowth associated proteins. Concordantly, *in vivo* embryonic Ebp1 knockdown selectively in early-born neocortical NSCs results in increased branching of neurites projected by maturing neurons in the cortical plate. This study is the first near-atomic resolution analysis of protein synthesis in the nervous system, positioning Ebp1 and the 60S peptide tunnel exit as a focal point of gene expression control during neocortical neurogenesis.

## RESULTS

### Neocortical ribosome mass spectrometry identifies Ebp1 as an abundant, high occupancy translation cofactor during development

To analyze the architecture of neocortical ribosome complexes across development, we first optimized a protocol to purify actively translating ribosomes *ex vivo* rapidly and stably without the use of chemical inhibitors, which bias the conformational state of the ribosome. The goal was to capture the full repertoire of integral translation cofactors binding ribosomes engaged in various stages of the translation cycle throughout neocortical neurogenesis, spanning the early neural stem cell (NSC) predominant embryonic stage (E12.5) to the post-mitotic neuronal post-natal stage (P0). We focused our analysis on complexes of isolated 80S ribosomes (monosomes), and chains of multiple 80S actively translating mRNA (polysomes) (**Figure 1A**). Neocortex lysates were fractionated by sucrose density gradient ultracentrifugation to purify 80S monosomes, actively translating polysomes, with corresponding input total lysates for mass spectrometry (MS) analysis (**Figure S1**). Sample reproducibility was observed in hierarchical clustering of the MS data, with the data clustering by biological triplicate, 80S vs. polysomes, and early vs. late developmental stages (**Figure S2**).

Results from the neocortical polysome MS are shown in **Figure 1B**, comparing protein levels at E12.5 with each subsequent developmental stage. As expected, core ribosomal proteins (RPs) were among the most enriched proteins in the polysomes, including RPs of the large 60S subunit (Rpl) and small 40S subunit (Rps). Known translation-associated proteins were enriched to varying degrees in polysomes throughout development. Unexpectedly, we observed Ebp1 co-purifying at levels comparable to the RPs themselves in polysomes, higher than any other translation-associated protein. Ebp1 is metazoan-specific and was only observed to play a niche role in protein synthesis, promoting non-canonical internal ribosome entry site (IRES) dependent translation of a specific viral mRNA (Pilipenko et al., 2000), and suppressing eIF2a phosphorylation in conditions of cellular stress (Squatrito et al., 2006), by unknown mechanisms. Largely studied in the context of cancer, Ebp1 influences cell proliferation and differentiation (Nguyen et al., 2018), in pathways including the epidermal growth factor receptor ErbB3 (Lessor et al., 2000; Yoo et al., 2000), and other mitogenic signaling cascades. Its role in the developing nervous system, general function in protein synthesis, and whether translational regulation is connected to its role in cancer are unknown. Thus, we were intrigued by Ebp1’s exceptionally high enrichment in polysomes of the developing neocortex, and observed a similarly robust association with 80S complexes measured by MS (**Figure S3A**). Furthermore, Ebp1 was among the most abundant proteins measured in total neocortical lysates across development (**Figure S3B**), similar to the RPs.

The core of the eukaryotic 80S ribosome is a macromolecular machine consisting of ∼79 RPs on a scaffold of 4 rRNAs, with translation-associated proteins transiently binding to catalyze and modulate ribosomal functions. To quantify the balance between translation-associated cofactors and core RPs in neocortical ribosomes across development, we generated stoichiometry matrices at the five developmental stages measured by MS (**Figures 1C and S3C-D**). Hierarchical clustering of these data visualizes the distribution of super-stoichiometric, stoichiometric, and sub-stoichiometric proteins associated with ribosomal complexes. The majority of core RPs approximate stoichiometric levels throughout development in polysome (**Figure 1C**) and 80S (**Figure S3C**) complexes, and are likewise maintained at similar levels in total steady state (**Figure S3D**). While the majority of translation-associated proteins are sub-stoichiometric to core RPs in ribosomal complexes, Ebp1 is nearly stoichiometric in polysome and 80S complexes. Across development in polysomes, the ratio of Ebp1 to the Rpl or Rps median is 0.63-0.86 to 1, indicating that Ebp1 may play an integral role in neocortical translation, rather than niche for a small subset of transcripts or during transient conditions as previously reported (Pilipenko et al., 2000; Squatrito et al., 2006).

### Ebp1 enrichment is cell type and temporally specific, scaling with the dynamic level of ribosomal complexes

The unusual abundance of Ebp1 as a translation-associated protein may correspond to a particular expression pattern in neocortical development. We next assessed the cell type and temporal specificity of *Ebp1* mRNA expression in scRNAseq data (Telley et al., 2019) measuring the transcriptome of early and late born NSCs maturing into lower and upper layer neurons, respectively (**Figure 2A**). Strikingly, *Ebp1* mRNA is particularly enriched in early born NSCs in the ventricular zone, with levels decreasing abruptly during both neuronal differentiation, and in the later born NSC pool. The particular enrichment of *Ebp1* mRNA is in contrast to *Rpl* and *Rps* mRNA expression patterns, which are maintained at stable levels in NSCs regardless of birthdate, but likewise decline during differentiation. This observation was reflected in analysis of total neocortical lysates across development by RNAseq (**Figure 2B**), with *Ebp1* mRNA steadily decreasing from E12.5, while *Rpl* and *Rps* mRNA decreases lag behind at E17. However, corresponding MS measurement revealed Ebp1 protein levels decline abruptly at E15.5 along with Rpl and Rps proteins in the neocortex, suggesting their levels are regulated in concert, with protein changes anticipating mRNA changes for the RPs. Notably, total Ebp1 protein is consistently maintained 2-5 fold higher than the median level of RPs, in contrast to the corresponding mRNA. Concordant with the above findings, immunohistochemistry analysis (**Figures 2C and S4**) demonstrated particularly high Ebp1 levels in the ventricular zone (VZ) and nascent cortical plate (CP) at E12.5-E14, including the ventricular and pial surfaces. Ebp1 is persistent in maturing neurons laminating the CP at later stages, albeit at lower levels. Interestingly, Ebp1 was observed in the P0 VZ that contains early gliogenic progenitor cells (Kriegstein and Alvarez-Buylla, 2009) at substantially lower levels than in the neurogenic E12.5 VZ. This observation may relate to a prior immunohistochemical study in the postnatal hippocampus, where Ebp1 was reported to be particularly enriched in neurons compared to astroglia (Ko et al., 2017). Thus, *Ebp1* enrichment is specific to both differentiation status and NSC birthdate in the neocortex.

**Figure 2.**
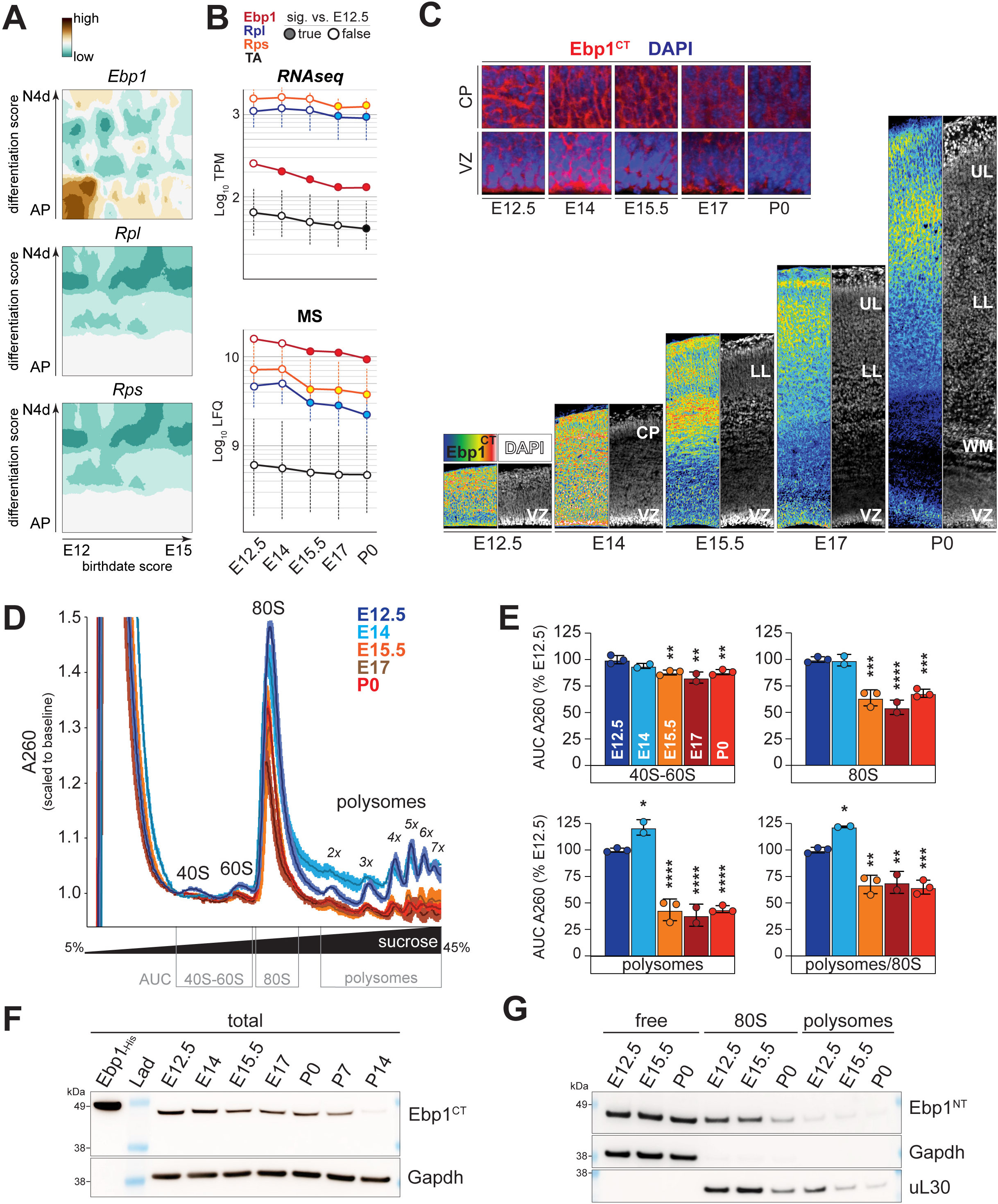
Ebp1 expression is cell type and temporally specific in early-born NSCs, in concert with transiently elevated ribosomal complex levels. **(A)** Expression heat maps of *Ebp1* compared to averaged *Rpl* and *Rps* family mRNA enrichment in scRNAseq analysis of the developing mouse neocortex derived from (Telley et al., 2019). Relative expression shown for apical progenitor (AP) NSCs during their differentiation into mature neurons (N4d) on the y-axis, corresponding to NSC birthdates E12, E13, E14, E15 on the x-axis. **(B)** Neocortical expression of *Ebp1* (red), *Rpl* (blue), *Rps* (yellow), and translation-associated (black) genes measured in total steady state levels by RNAseq (top) and MS (bottom) across developmental stages. The median expression is plotted ± s.d. Significant changes assessed by one-way ANOVA and Bonferroni corrected *post hoc* test vs. E12.5, *p* < 0.05 considered significant. **(C)** Immunohistochemistry analysis of Ebp1 expression in coronal sections of the developing neocortex ventricular zone (VZ) populated by NSCs, and cortical plate (CP) populated by maturing neurons. Early-born NSCs generate lower layer (LL) neurons, while later-born NSCs generate upper layer (UL) neurons. White matter (WM) axons, nuclear DAPI staining in grey. Zoomed images (inset) correspond to the VZ and leading-edge of the CP at each stage, with Ebp1 (red) and DAPI (blue) labeling. Similar results were obtained with the Ebp1^NT^ antibody (**Figure S4**). **(D)** Analytic quantitative sucrose density gradient ultracentrifugation of A260 normalized neocortical lysates measuring the relative abundance of ribosomal subunits, 80S ribosomes, and polysomes. A260 curves plotted as mean ± s.d. across replicates for each stage. **(E)** Curves from (D) subdivided for area-under-the-curve (AUC) analysis as shown (grey boxes) calculated by a Reimann sum, and graphed as mean ± s.d. with significance testing by one-way ANOVA and Dunnett’s *post hoc* test vs. E12.5. \**p* < 0.05; \*\**p* < 0.01; \*\*\**p* < 0.001; \*\*\*\**p* < 0.0001. **(F)** Western blot analysis of Ebp1 enrichment in total neocortical lysates, in comparison to full-length histidine-tagged Ebp1 (Ebp1-His). Full blots are shown in **Figures S5A-B**. **(G)** Western blot analysis of Ebp1 enrichment in pooled extra-ribosomal (free), 80S, and polysome fractions (**Figures S5C-D**).

We observed that, along with Ebp1, total steady state RP levels decrease at E15.5-P0 in the developing neocortex by MS (**Figure 2B**). We next asked whether this reflects a timed global shift in the balance of actively translating ribosomal complexes during prenatal neurogenesis. Neocortical lysates were subjected to analytic quantitative sucrose density gradient fractionation corresponding the stages analyzed by MS, loading equivalent A260 optical density units to directly compare the distribution of 40S-60S, 80S, and polysome levels at each stage (**Figure 2D**). Gradient curves demonstrated a timed decrease from high levels of 80S and polysomes at E12.5-E14, to a lower steady state from E15.5-P0. A decrease of 35% 80S and 64% polysome levels during the transition from E14 to E15.5 was calculated by area-under-the-curve with a Riemann sum (**Figure 2E**). This decrease in ribosomes is not wholly accounted for by the availability of individual subunits in the cytoplasm, as 40S-60S levels decrease 4% from E14 to E15.5. Taken together, these findings suggest mature, active ribosomal complexes exist at elevated levels during early neocortical neurogenesis, and transition to a lower steady state level at later stages, concordant with MS findings. These data are in line with previous observations of RP downregulation in the mouse forebrain between E8.5-E10.5 during neural tube closure (Chau et al., 2018), and dynamic levels of ribosomal complex proteins in the E13-P0 neocortex (Kraushar et al., 2015). Global shifts in steady state ribosomal complex levels may reflect a dynamic equilibrium of cellular homeostasis (Delarue et al., 2018; Mills and Green, 2017; Sinturel et al., 2017) in neocortex development.

Ebp1 has been previously reported to exist as a full-length 48kDa protein (“p48”), and a 42kDa isoform (“p42”) with a 54 amino acid N-terminal truncation generated by *Ebp1* mRNA splicing (Liu et al., 2006). Total neocortical lysates across developmental stages were analyzed by Western blot, probing for Ebp1 with a C-terminal targeting antibody (Ebp1^CT^) that recognizes both long and short isoforms (**Figures 2F and S5A**), and with a N-terminal specific antibody (Ebp1^NT^) that recognizes only full-length Ebp1 (**Figure S5B**), compared to signal for recombinant full-length Ebp1 with a N-terminal histidine tag (Ebp1-His). Results showed that the dominant protein isoform of Ebp1 in neocortical development is full-length. Furthermore, the Western blot measurements mirror the MS measurements (**Figure 2B**), demonstrating higher total Ebp1 levels at E12.5-E14, with a decrease at E15.5 to a lower steady state. Notably, total neocortical Ebp1 levels are higher overall in the embryonic and perinatal period, while measurements at P7, and particularly P14, show decreased enrichment – in agreement with previous measurements in whole brain lysates (Ko et al., 2017). Taken together, full-length Ebp1 is the dominant isoform expressed in the neocortex with a particular enrichment in embryonic development, and its levels decrease at E15.5 in concert with a global decrease in 80S and polysome levels.

We next sought to measure the balance of ribosome-associated Ebp1 compared to “free” extra-ribosomal Ebp1 in neocortical development. Input normalized neocortical lysates at E12.5, E15.5, and P0 were fractionated to separate extra-ribosomal free Ebp1 vs. 80S and polysome associated Ebp1 (**Figure S5C**). Western blot analysis of individual gradient fractions (**Figure S5D**), and pooled fractions constituting free, 80S, and polysome complexes (**Figure 2G**) showed that free extra-ribosomal Ebp1 is maintained at high levels throughout development, while only ribosome-associated Ebp1 decreases at E15.5. These findings suggest that total Ebp1 levels decline from E12.5-P0 secondary to a decrease in ribosomal complexes along with ribosome-associated Ebp1.

### Ebp1 binds the 60S with high affinity in the cytoplasm of neocortical neural stem cells and neurons

While previous studies suggested that Ebp1-ribosome interaction is rRNA-dependent and dissociates in high salt conditions (Squatrito et al., 2004, 2006), its specific binding mode is unknown. We next performed *in vitro* binding assays with recombinant Ebp1 and purified 40S, 60S, and reconstituted 80S derived from rabbit reticulocyte lysate (RRL). Samples were pelleted through a sucrose cushion, followed by Western blot analysis of the supernatant (unbound Ebp1) and pellet (ribosome-bound Ebp1) (**Figure 3A**). Ebp1 co-pelleted with the 60S and 80S exclusively, demonstrating concomitant decreases of Ebp1 in the supernatants. These findings were reinforced by a binding assay with mouse neocortex derived 40S and 60S (**Figure S6A**). Thus, Ebp1 specifically binds the 60S subunit and is persistent in 80S complexes, suggesting Ebp1 interactions do not interfere with the subunit interface, nor does its binding require mRNA. Furthermore, Ebp1•60S binding is conserved between mouse and rabbit species, and across reticulocyte and neocortical derived ribosomes.

**Figure 3.**
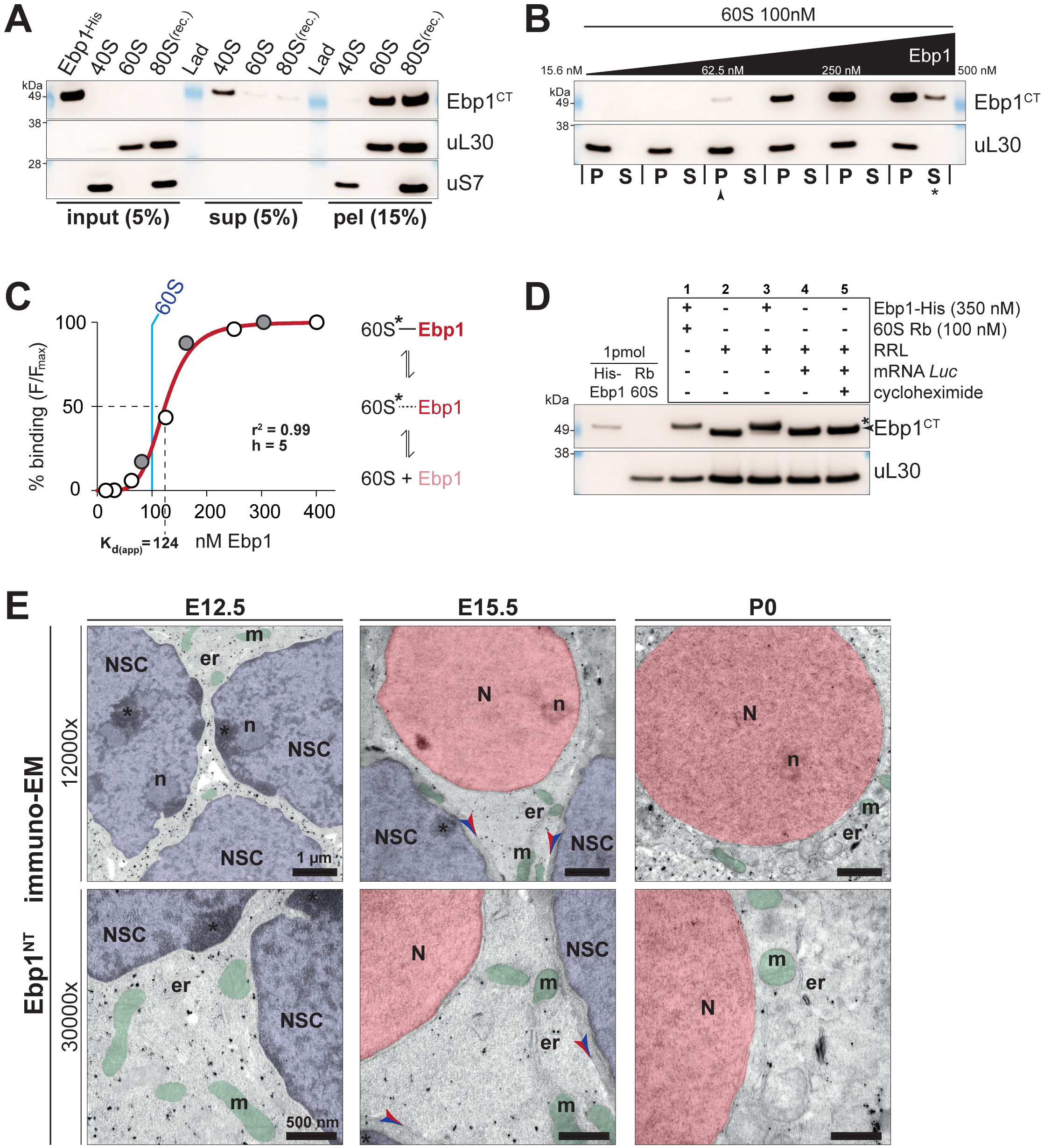
Ebp1 binds the 60S subunit with high affinity in the cytoplasm of neocortical NSCs and neurons. **(A)** Western blot analysis of recombinant Ebp1-His binding to purified rabbit reticulocyte lysate (RRL) 40S, 60S, and reconstituted 80S in the pellet (pel) compared to unbound in the sucrose cushion supernatant (sup). Ebp1 signal shown compared to 60S RP (uL30) and 40S RP (uS7) markers. Binding assay with mouse neocortical 40S and 60S shown in **Figure S6A**. **(B)** Relative binding affinity of escalating Ebp1-His concentrations (15.6-500nM) to 100nM 60S measured by pelleting assay and Western blot analysis. Binding first detected in the pellet (P) at 62.5nM (arrow), with excess Ebp1 in the supernatant (S) seen at 500nM (star) in comparison to saturating signal in the pellet. **(C)** Results from (B) (white circles) plotted with an independent replicate experiment (grey circles), and curve best fit to the data (red line, nonlinear least squares fit, with one-site binding and Hill slope accommodation). Interpretation of binding curves further described in **Figure S6B**. **(D)** Ebp1 binding dynamics assessed by pelleting assay and Western blot. Binding pellet signal for (1) super-saturating levels of Ebp1-His (350nM) in the presence of rabbit 60S (100nM) compared to (2) native Ebp1 in RRL (∼100nM ribosomes), (3) saturating Ebp1-His in RRL, and native Ebp1 in RRL translating *Luciferase* mRNA (*Luc*) with (4) and without (5) cycloheximide to stall elongation. Signal for native Ebp1 (arrow) in comparison to the slightly larger Ebp1-His (star). **(E)** Immuno-electron micrographs showing immunogold labeling for anti-Ebp1^NT^ (black dots) in coronal sections of the neocortex at E12.5, E15.5, and P0 at low (12000x) and high (30000x) magnification. Neural stem cells (NSC) and neurons (N) are identified by their distinctive nuclear morphology (blue, NSC; red, N) and their localization in the developing cortical layers (ventricular zone, NSC; expanding cortical plate, N). Nucleoli (n), mitochondria (m, green), endoplasmic reticulum (er), cell-cell junctions (arrows; blue on NSC side, red on N side). See also **Figure S7**.

We next determined the relative affinity range of Ebp1•60S binding, where experiments were similarly conducted with a constant rabbit 60S concentration, combined with doubling concentrations of Ebp1 measured in the pellet vs. supernatant (**Figure 3B**), and data plotted (**Figure 3C**). The curve best fit to data (r^2^=0.99) indicates Ebp1 reaches a K_d(app)_ at ∼124 nM, with saturated Ebp1•60S binding at ∼200 nM, relative to 100 nM 60S. These data indicate Ebp1 binds the 60S with a high relative affinity, reaching saturation at ∼2-fold excess Ebp1 over the 60S. Given the 2-5 fold excess of total steady state Ebp1 levels over the 60S RPs measured by MS (**Figure 2B**), superstoichiometric Ebp1 would be hypothetically sufficient to yield a high degree of Ebp1•60S association *in vivo*. Furthermore, a nonlinear least squares fit, with one-site binding and Hill slope accommodation, suggested Ebp1 binding includes 60S conformational activation (**Figures 3C and S6B**).

While isolated 60S was sufficient for Ebp1 binding, whether its binding mode undergoes dynamic Ebp1 turnover, is diminished/enhanced by active mRNA translation, or can be modulated by translation inhibitors is unknown. To answer these questions, the following binding conditions were constituted in parallel: (1) saturating levels of Ebp1-His in the presence of rabbit 60S; (2) RRL containing native Ebp1; (3) saturating Ebp1-His added to RRL; native Ebp1 in RRL translating *Luciferase* mRNA (*Luc*) with (4) and without (5) cycloheximide to stall elongation. Results are shown in **Figure 3D**, with Ebp1-His and 60S inputs as markers for the binding pellet of each condition (1-5). Native Ebp1 in RRL (2) co-pelleted with the ribosome as did Ebp1-His to the 60S (1), undergoing dynamic binding (3) demonstrated by the nearly complete turnover of native Ebp1 with saturating Ebp1-His. Active *in vitro* translation of a *Luciferase* mRNA (4) did not impact the stability of Ebp1•60S binding, nor did elongation stalling by cycloheximide (5). These findings indicate the Ebp1•60S binding mode occurs with dynamic turnover, in conditions irrespective of active mRNA translation.

Ebp1 has been reported to localize to both the cytoplasm and nucleus/nucleolus in cultured cells lines (Liu et al., 2006; Radomski and Jost, 1995). Whether Ebp1•60S complexes are formed during 60S assembly in the nucleolus as previously suggested (Squatrito et al., 2004), at the nuclear membrane for 60S export, or on mature 60S in the cytoplasm remained unclear in the neocortex. We next prepared coronal sections of the neocortex at E12.5, E15.5, and P0 for immuno-electron microscopy (immuno-EM), probing for Ebp1 with both Ebp1^NT^ (full-length isoform only; **Figure 3E**) and Ebp1^CT^ (full-length and truncated isoforms; **Figure S7A**) antibodies. At all developmental stages analyzed, immunogold labeling for Ebp1 demonstrated predominantly cytoplasmic signal, occurring in clusters throughout the cytoplasm of both NSCs in the ventricular zone, and neurons populating the cortical plate. Ebp1 was largely absent from the nuclei and nuclear membrane of NSCs and neurons, including the nucleolus. Furthermore, Ebp1 was not observed in mitochondria, or in strict proximity to the endoplasmic reticulum or plasma membrane. Notably, Ebp1 was also observed in dendrites of maturing neurons at P0 (**Figure S7B**), suggesting Ebp1 localizes throughout cytoplasmic compartments of both neocortical NSCs and neurons. Thus, Ebp1 may function to regulate protein synthesis with 60S binding in the cytoplasm, rather than as a nuclear assembly or export factor, during neocortical neurogenesis.

### The structure of neocortical Ebp1•ribosome complexes *ex vivo* at near-atomic resolution

To analyze the architecture of neocortical ribosome complexes and visualize the physiologic binding mode of Ebp1 at near-atomic resolution, 80S and polysomes were purified by sucrose density gradient fractionation from P0 neocortex lysates, pooled together, and frozen on grids for cryo-electron microscopy (cryo-EM). Micrographs confirmed the presence of both 80S and polysome complexes in the sample (**Figure 4A**). High-resolution cryo-EM data collection (**Figure S8**) and initial single-particle reconstruction yielded a map of the complete 80S, along with extra-ribosomal density (red) adjacent to the 60S peptide tunnel exit (**Figure 4B**). Fitting the crystal structure of mouse Ebp1 (PDB 2V6C) (Monie et al., 2007) to the extra-ribosomal density unequivocally identified Ebp1 in complex with the neocortical 60S. Furthermore, robust density was present for nearly the entire N-terminus, identifying the full-length isoform of Ebp1 is bound. The direct visualization of native full-length Ebp1 binding to 60S *ex vivo* strongly supports the physiologic nature of this Ebp1•60S binding mode in the neocortex.

**Figure 4.**
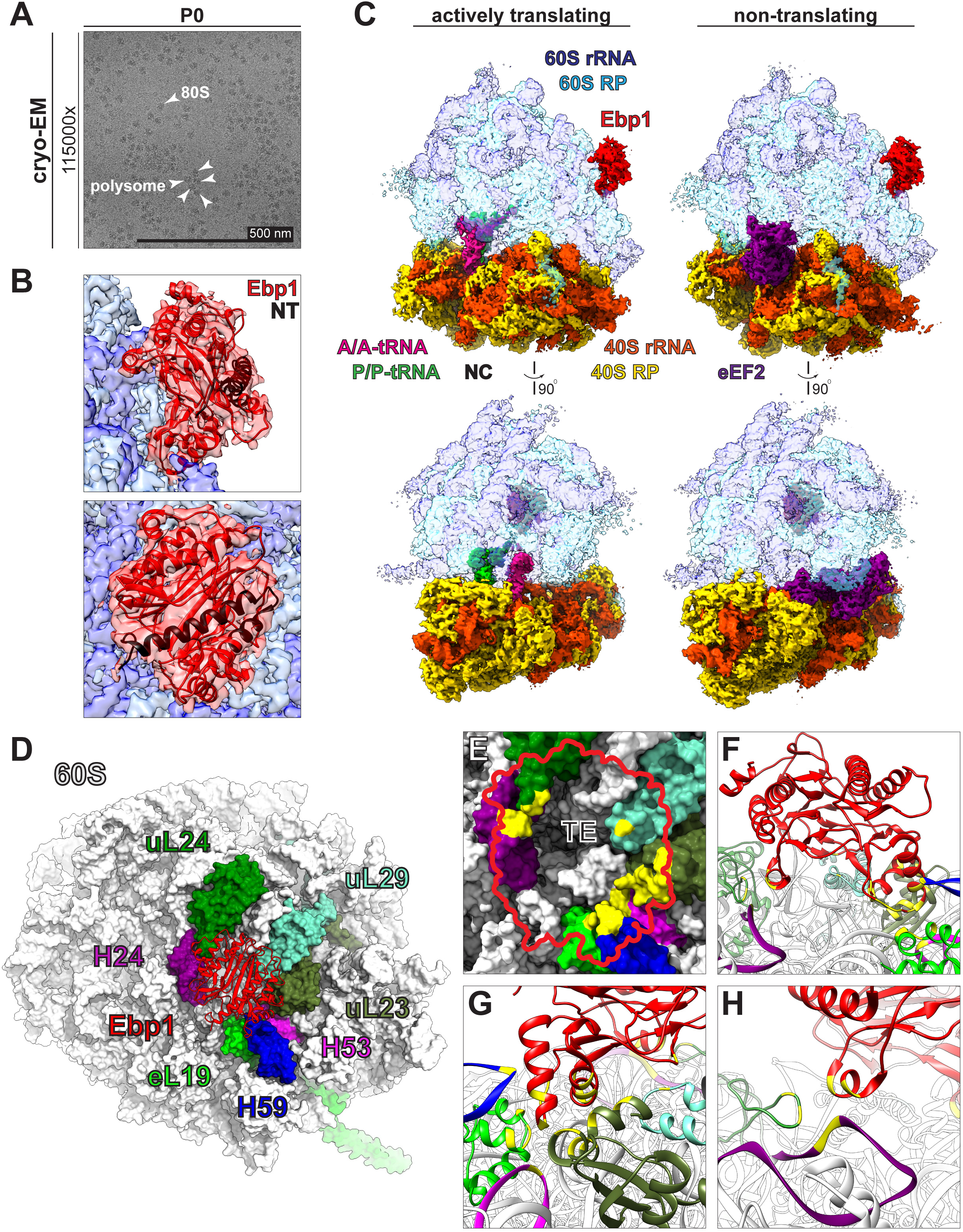
Structure of the neocortical ribosome with Ebp1 bound to the 60S subunit at the peptide tunnel exit. **(A)** Cryo-electron microscopy (cryo-EM) micrograph of pooled 80S and polysome complexes (arrows) from P0 mouse neocortical lysates *ex vivo*. **(B)** Cryo-EM map generated by 3D refinement of (A) with extra-ribosomal density (red) conforming to the crystal structure of mouse Ebp1 (PDB 2V6C (Monie et al., 2007), red ribbon) on the 60S surface (rRNA, dark blue; RPs, light blue). Ebp1 density includes the N-terminal residues (NT, black ribbon) corresponding to full-length Ebp1. **(C)** Multiparticle classification and high-resolution map refinement (**Figures S8-S10**) of both actively translating (left, classical state with A/A, pink, and P/P, green, tRNAs) and non-translating (right, rotated state with eEF2, purple) ribosomal complexes. Ebp1 (red), 60S rRNA (dark blue), 60S RPs (light blue), 40S rRNA (orange), 40S RPs (yellow), nascent chain fragments (black). **(D)** Model of the Ebp1 (red ribbon) binding surface at the 60S peptide tunnel exit, including 60S rRNA helices H24 (purple), H53 (magenta), H59 (blue), and 60S RPs eL19 (lime green), uL23 (olive), uL24 (forest green), uL29 (aquamarine). **(E)** Aerial view of the Ebp1 footprint (red outline) over the 60S peptide tunnel exit (TE), with rRNA helices and RP model surfaces colored as in (D), and residues/nucleosides making electrostatic interactions with Ebp1 highlighted in yellow. **(F-H)** Zoomed views of the Ebp1•60S model, colored as in (D-E), with binding residues/nucleosides likewise highlighted in yellow.

To disentangle the ribosome conformational states bound by Ebp1, we proceeded with hierarchical multiparticle sorting and 3D classification of both large and small scale heterogeneity intrinsic to the data (Behrmann et al., 2015; Loerke et al., 2010) (**Figure S9**). Ribosome complexes in both the rotated and classical conformations were first sorted, including populations with (1) eEF2 and (2) eEF2+P/E tRNA in the rotated state, and populations with (3) A/A+P/P tRNAs, (4) E/E tRNA, and (5) without tRNAs in the classical state. In each of these five states, a strategy of modified focused classification (see Methods) was utilized to separate sub-states with and without Ebp1, yielding ten total classes. Across all states, Ebp1 was bound to 48% of ribosomes. Likewise, Ebp1 was bound to ∼50% of the ribosomes within each of the five sub-states. We proceeded with high-resolution refinement of Ebp1-bound and unbound populations in the rotated state with eEF2 (3.1 Å global resolutions), and the classical state with A/A+P/P tRNAs (3.3 Å global resolutions). High-resolution cryo-EM maps are shown in **Figure 4C**, representing both actively translating (classical state with A/A+P/P tRNAs) and non-translating (rotated state with eEF2) ribosomes with Ebp1 bound to the 60S. Taken together, our findings suggest Ebp1 occupies its neocortical 60S binding site with high occupancy *in vivo* based on both MS estimates (**Figures 1B-C**), and conservative estimates with multiparticle sorting (**Figure S9**) (assuming some destabilization during sample freezing), approximating 50% occupancy. Furthermore, Ebp1 binds to both actively translating and non-translating neocortical ribosome states with approximately equal probability.

The near-atomic resolution of our data (**Figures S8 and S10**) permitted modeling of the entire neocortical Ebp1•60S complex. **Figure 4D** visualizes the peptide tunnel exit (TE) surface in electrostatic proximity to Ebp1, including four RPs (eL19, uL23, uL24, uL29) and three rRNA helices (H24, H53, H59). An aerial view of the Ebp1 footprint over the TE surface highlights the 60S RP residues and rRNA nucleosides making electrostatic interactions with Ebp1 (**Figure 4E**), demonstrating that Ebp1 contacts the immediate TE surface. A side view of the Ebp1 model at the TE (**Figure 4F**) shows Ebp1 forming a concavity above the TE vestibule, stabilized by electrostatic interactions (**Figures 4G-H**) at the concavity rim with gaps (∼26 Å at the widest point) that would permit peptide chain exit. Therefore, Ebp1 binds the 60S peptide TE, creating a pocket above the TE vestibule with a porous interface.

### Ebp1 binding requires a conserved 60S helix H59-H53 swinging latch mechanism

Multiparticle sorting of our data into Ebp1-bound and unbound states enabled identification of 60S structural changes facilitating Ebp1 interactions with an internal negative control (**Figures S11A-B**). We observed that in the Ebp1-bound state, the tip of H59 undergoes a backbone rearrangement enabled by a 235**°** flip of H59 G-2690, releasing contact with H53 G-2501, G-2502, and C-2513 as seen in the canonical unbound state (**Figure 5A**) – resulting in H59 G-2690 transitioning to intra-helical base stacking interactions. This “swinging latch” mechanism further includes a 73**°** flip of H59 U-2687, with the base reaching into the insert domain of Ebp1 (**Figure S11C**), locking Ebp1 into position. This particular movement of H59 U-2687 was previously observed for the binding of the yeast nuclear export (Bradatsch et al., 2007) and peptide tunnel quality control factor (Greber et al., 2016) Arx1 to the 60S – thus representing a conserved binding mechanism. However, unlike Arx1, Ebp1 binding does not require stabilization by rRNA expansion segment ES27 on the solvent side (Greber et al., 2016), suggesting that aspects of its binding mode are distinct. Furthermore, the concerted restructuring of H59 may represent a 60S “activation step” to facilitate Ebp1 binding, reflected in the Ebp1•60S binding curve with Hill slope accommodation (**Figures 3C and S6B**). Thus, in this model, increasing concentrations of Ebp1 increase the probability that H59 is stabilized in the activated structural state, permitting Ebp1 re-binding events to occur more frequently for higher aggregate 60S occupancy. Taken together, Ebp1-ribosome binding requires a 60S H59 swinging latch mechanism likewise utilized by yeast Arx1.

**Figure 5.**
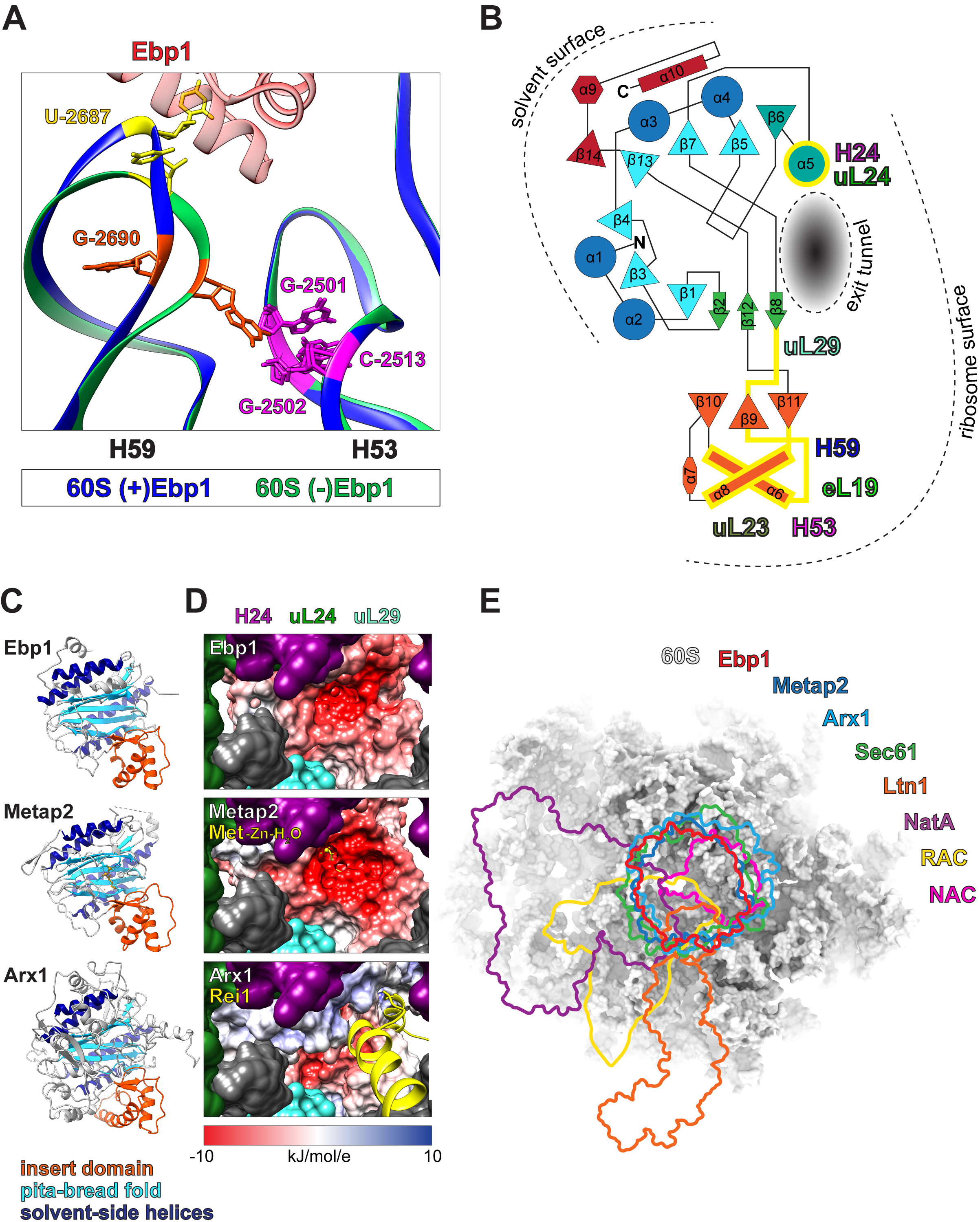
The Ebp1 insert domain utilizes a conserved H59 latch mechanism for 60S binding incompatible with simultaneous binding by other tunnel exit proteins. **(A)** 60S rRNA H59 and H53 models corresponding to states with Ebp1 (blue) vs. without Ebp1 (green), adjacent to the Ebp1 (red) insert domain. See also **Figures S11A-C**. **(B)** 2D structure diagram of Ebp1 3D domains adapted from (Kowalinski et al., 2007), orienting Ebp1 on the ribosome surface, and highlighting the domains involved in Ebp1•60S binding (yellow). **(C)** Global alignment of Ebp1, Metap2, and Arx1 demonstrate the conserved orientation of the β-α---α-β insert domain (orange) for 60S binding, pita-bread β6 fold motif (light blue) positioned over the TE, and solvent-side α4 motif (dark blue). See also **Figure S12**. **(D)** Electrostatic potential maps of Ebp1, Metap2, and Arx1 oriented as in (C), and viewed from within the TE from the perspective of emerging nascent chain, with 60S rRNA helices and RPs colored as in Figure 4D. Methionine•Zn•H_2_O and Rei1 models in yellow. See also **Figure S11D**. **(E)** Schematic with the mouse neocortical 60S model surface centered on the peptide tunnel exit, and the overlapping footprints of known eukaryotic tunnel exit surface binding factors. See also **Figure S13**. PDB IDs: Metap2 1KQ9 (Nonato et al., 2006), Arx1 5APN (Greber et al., 2016), Sec61 3J7R (Voorhees et al., 2014), Ltn1 3J92 (Shao et al., 2015), NatA 6HD7 (Knorr et al., 2019). EMDB IDs: RAC 6105 (Zhang et al., 2014), NAC 4938 (Gamerdinger et al., 2019).

### Ebp1**•**60S binding is incompatible with simultaneous binding of all other eukaryotic peptide tunnel exit cofactors

The neocortical Ebp1•60S complex establishes previously unassigned functions to Ebp1 structural domains (**Figure 5B**; adapted from (Kowalinski et al., 2007)). Ribosome binding by Ebp1’s insert domain and α5 helix positions β-sheets 1, 3, 4, 5, 7, and 13 directly over the TE. The alignment of mouse Ebp1 with orthologs across eukaryotic taxa, in addition to Metap2 and Arx1, demonstrates conservation of key binding resides (**Figure S12**). Previous studies have commented on the structural similarity between Ebp1 and the methionine aminopeptidase Metap2 (Kowalinski et al., 2007; Monie et al., 2007), and between Arx1 and Metap2 (Greber et al., 2016). Indeed, Ebp1, Metap2, and Arx1 share similar structural features, and putative binding motifs (**Figure 5C**). A common β-α---α-β insert domain facilitates 60S binding, the “pita-bread” β6 fold motif is positioned over the peptide TE, and a solvent-side α4 motif is available for potential molecular interactions. Their binding ultimately creates different electrochemical environments at the TE (**Figure 5D**). In the event of Ebp1 or Metap2 binding, emerging peptide chain would encounter a deep, strongly electronegative pocket; however, the key residues in the Metap2 β-sheet pita-bread fold catalyzing aminopeptidase activity (Nonato et al., 2006) are absent in Ebp1 (Kowalinski et al., 2007; Monie et al., 2007), rendering Ebp1 catalytically inactive. Furthermore, the Ebp1 α5 domain facilitating electrostatic contacts with H24 and uL24 is absent in Metap2 (**Figure S11D**); however, a Metap2•60S complex structure has not yet been solved, and thus Metap2 structural adjustments may exist. In contrast, the yeast Arx1 pita-bread fold threads Rei1 into the peptide tunnel to probe the 60S as a quality-control step preempting active translation (Greber et al., 2016). Thus, while sharing structural features and a conserved binding mode, the binding of Ebp1, Metap2, and Arx1 engage distinct functional states of 60S-nascent chain complexes, with an emerging nascent peptide chain encountering distinct environments.

The binding of Ebp1 would be sterically incompatible with the simultaneous docking of other 60S TE cofactors, competing for limited real estate surrounding an emerging nascent peptide chain (**Figure 5E**). The footprint of Ebp1 is shown superimposed on the footprints of Metap2 (Nonato et al., 2006) and Arx1 (Greber et al., 2016), in addition to: the ER translocation channel Sec61 (Voorhees et al., 2014); the Ltn1-NEMF ubiquitin ligase complex (Shao et al., 2015); the N-terminal acetyltransferase NatA (Knorr et al., 2019); the ribosome-associated complex (RAC) coupling nascent chain elongation and folding (Zhang et al., 2014); and the nascent polypeptide-associated complex (NAC) preventing ER mistargeting and suppressing aggregation of synthesized proteins (Gamerdinger et al., 2015; Shen et al., 2019). While the particular abundance of Ebp1 among these TE cofactors would potentially support its occupancy, the dynamic turnover of Ebp1 might allow for other TE cofactors to bind if recruited by their associated nascent chain moieties as needed. Furthermore, the neocortical cell-type and temporal specificity of Ebp1 enrichment is in contrast to some of the other TE cofactors, such as Ltn1, while similar to others, such as RAC (**Figure S13**). Dynamic enrichment of Ebp1 vs. other TE cofactors in neocortical development may represent the dynamic regulation of protein synthesis in response to the unique demands of particular stages of neurogenesis. Given Ebp1’s high occupancy of the neocortical ribosome tunnel exit, we hypothesized that depletion of Ebp1 would disrupt the balance of proteostasis in neuronal protein synthesis.

### Neuronal Ebp1 regulates acute protein synthesis and chronic proteostasis impacting axonal, dendritic, and synaptic proteomes

To interrogate the potential function of Ebp1 in maintaining neuronal proteostasis, we established a system to measure the impact of Ebp1 depletion on acute protein synthesis and chronic proteostasis in a mouse neuronal cell line. Neuro2a cells dominantly express the full-length isoform of Ebp1 (**Figure S14A**), which associates with 80S and polysomes (**Figures S14B-C**), similar to mouse neocortex. We next confirmed robust and specific knockdown of Ebp1 in Neuro2a by siRNA, with *siEbp1* targeting the mouse sequence effecting nearly complete knockdown, in contrast to oligos targeting the human sequence, and non-targeting control (**Figure 6A**).

**Figure 6.**
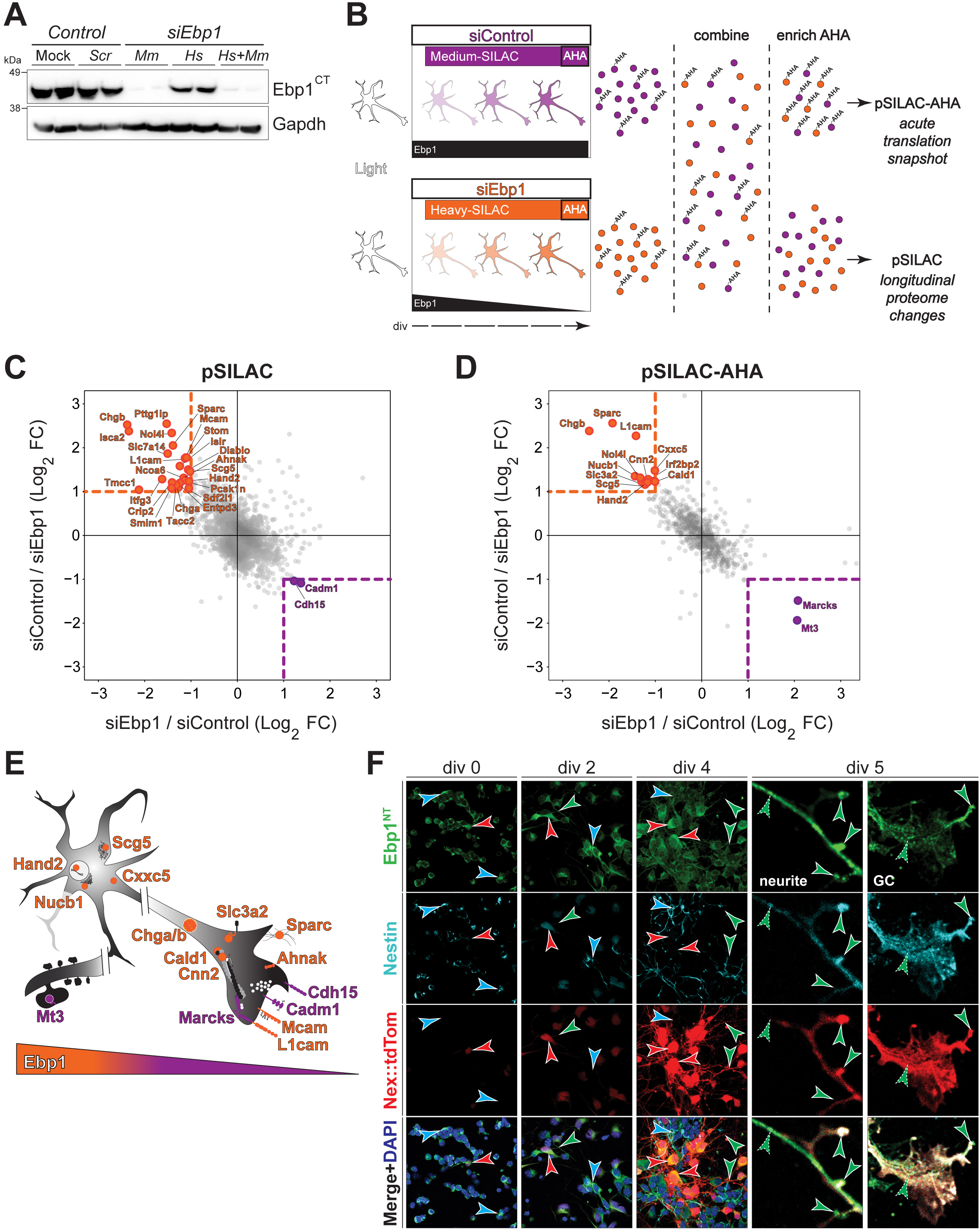
Ebp1 knockdown disrupts acute protein synthesis and chronic proteostasis in neuronal cells, impacting neurite and synaptic proteins. **(A)** Western blot confirmation of Ebp1 knockdown in mouse Neuro2a cells with siRNA oligos targeting mouse vs. human *Ebp1* mRNA sequences, in comparison to mock transfection and scrambled siRNA controls, in biological duplicate. **(B)** Schematic of the strategy to measure both the chronic proteostasis and acute protein synthesis responses to Ebp1 knockdown in Neuro2a cells with pSILAC and BONCAT coupled mass spectrometry. Cultures with transfection of *siControl* vs. *siEbp1* were concurrently incubated with medium (purple) vs. heavy (orange) SILAC amino acids, respectively, after the initiation of knockdown to label all proteins synthesized while Ebp1 levels are depleted (pSILAC) over 3 days *in vitro* (div). On div 4, AHA was pulsed before sample collection to label newly made proteins at the nadir of Ebp1 knockdown (pSILAC-AHA). **(C)** pSILAC and **(D)** pSILAC-AHA labeled protein levels with *siEbp1* conditions plotted relative to *siControl* conditions. Proteins with significantly lower (orange) or higher (purple) levels in *siEbp1* conditions are highlighted, while unchanged proteins are shown in grey. Confirming robust knockdown, Ebp1 levels were below the MS quantification threshold in *siEbp1* conditions, and thus not plotted. The threshold for significant change is set to >2-fold change from control in both replicates (dotted lines). **(E)** Schematic representation of the subcellular localization and function of known neuron-associated proteins significantly impacted by Ebp1 depletion from (C-D). Proteins with levels promoted by Ebp1 (*siEbp1* < control) are highlighted in orange, and proteins suppressed by Ebp1 (*siEbp1* > control) are highlighted in purple. **(F)** Primary neuronal cultures from the E12.5 neocortex of *Nex:Cre;Ai9* mice analyzed by immunocytochemistry at div 0, 2, 4, and 5 to monitor Ebp1 expression and localization in Nestin-positive (cyan arrows) neural stem cells (NSCs) throughout their maturation into Nex-postive (red arrows) postmitotic pyramidal neurons. Ebp1 localization in growing neurites is indicated (green arrows). Zoomed images at div 5 with clustered Ebp1 foci in neurites and growth cones (dotted arrows), including the leading edge of neurite protrusions (solid arrows). See also **Figure S14**.

The strategy to measure the response of the Neuro2a proteome with a combination of pulsed stable isotope labeling by amino acids in cell culture (pSILAC) (Schwanhäusser et al., 2009) and bioorthogonal noncanonical amino acid tagging (BONACT) (Dieterich et al., 2006) by MS (Eichelbaum et al., 2012; Howden et al., 2013) in this Ebp1 knockdown system is shown in **Figure 6B**. SILAC isotopes labeled all newly made proteins throughout the course of Ebp1 knockdown for longitudinal proteome changes, while pulse labeling with a methionine analog (AHA) captured a snapshot of newly synthesized proteins at the nadir of Ebp1 knockdown. pSILAC (**Figure 6C**) and pSILAC-AHA (**Figure 6D**) proteomes were measured by LC-MS/MS, and analyzed for differential protein enrichment in *siEbp1* vs. *siControl* conditions. Importantly, Ebp1 levels were below the quantification threshold in *siEbp1* conditions, confirming robust knockdown. Thus, this approach captured the impact of Ebp1 depletion on both longitudinal proteostasis (pSILAC) and acute protein synthesis (pSILAC-AHA) in neurons.

Results for both the pSILAC and pSILAC-AHA MS showed that in *siEbp1* conditions, proportionately more proteins decrease compared to those with increased protein levels relative to control, suggesting Ebp1 largely enhances protein expression. Cell adhesion molecules (CAMs) (de Wit and Ghosh, 2016), such as L1cam, Mcam, Cadm1, and Cdh15, were particularly impacted by Ebp1 depletion. Notably, Ebp1 maintains the balance of CAM levels, promoting L1cam (Maness and Schachner, 2007) and Mcam (Taira et al., 2005), while suppressing Cadm1 (Robbins et al., 2010) and Cdh15 (Bhalla et al., 2008), in addition to regulating CAM modulators such as Slc3a2 (Feral et al., 2005). CAMs play a critical role in neuronal migration, synaptogenesis, and neurite outgrowth/branching during development, plasticity, and disease (Biederer et al., 2017; Maness and Schachner, 2007; Missler et al., 2012; de Wit and Ghosh, 2016). Notably, six proteins were detected as significantly changing in common between the pSILAC and pSILAC-AHA datasets, such as L1cam, reinforcing that their regulation by Ebp1 is direct and protein synthesis specific, rather than a secondary effect. Collectively, gene ontology (GO) analysis of Ebp1 regulated proteins (**Figure S14D**) demonstrated cell adhesion (*P*<0.01; biological process) and secretory granule (*P*<0.01; cellular component) pathways as the most significantly impacted. Some of Ebp1-regulated proteins are predominantly locally translated in neurites (Zappulo et al., 2017), such as Cnn2, Mcam, and Sparc (**Figure S14E**). A schematic summarizing the impact of Ebp1 depletion on proteins with a known neuronal function is shown in **Figures 6E** and **S14F**, implicating Ebp1 in the regulation of cell-cell adhesion, synaptogenesis, and neurite outgrowth in neurons.

Given that Ebp1 influences the neurogenic proteome associated with neuronal processes in Neuro2a cells, and is enriched in early-born neocortical NSCs, we next sought to visualize native Ebp1 expression during the progressive differentiation of early-born neocortical NSCs into post-mitotic pyramidal neurons undergoing neurite outgrowth. Primary cultures were prepared from the E12.5 neocortex of *Nex:Cre;Ai9* mice (Turko et al., 2018), which label post-mitotic pyramidal neurons with tdTomato by activation of the *Nex* locus, followed by immunohistochemical analysis of Ebp1 expression at div 0, 2, 4, and 5 (**Figure 6F**). Ebp1 is enriched in cytoplasmic foci colocalizing with Nestin labeling in NSCs at div 0, in addition to the earliest differentiating Nex-positive cells. Cytoplasmic Ebp1 expression persists in differentiating Nex-positive neurons and extends into growing neurites, albeit at overall lower levels with differentiation. A decreasing enrichment pattern in dissociated cell culture reinforces *Ebp1* mRNA (**Figures 2A-B**) and protein (**Figures 2B-C and S4**) levels in neocortex tissue, suggesting that the trajectory of decreased Ebp1 enrichment during neuronal maturation may be cell autonomous, rather than in response to a signal in the tissue environment. At div 5, puncta of Ebp1 expression in neurites and growth cones is particularly apparent with further magnification, including the most distal aspects of extending processes. Ebp1 localization to neuronal processes and growth cones is reinforced by prior cell culture studies (Ko et al., 2017; Kwon and Ahn, 2011), which observed enhancement of axon regeneration after injury with Ebp1 overexpression in hippocampal slice culture. Taken together, these findings correlate with Ebp1’s maintenance of neurogenic, neurite, and synaptic proteostasis. Therefore, we hypothesized Ebp1 regulates neurite outgrowth in early-born neocortical NSCs.

### Ebp1 regulates neurite branching of early-born neocortical neurons *in vivo*

Since we observed particularly high Ebp1 enrichment in early-born NSCs of the developing neocortex (**Figures 2A-C and S4**), we next sought to study the effect of early Ebp1 depletion in NSCs during their maturation into neocortical neurons *in vivo*. *In utero* electroporation (IUE) of a *shEbp1* knockdown plasmid along with a CAG-GFP transfection reporter at E12 was compared to scrambled shRNA control, followed by analysis at E16 during initial neurite outgrowth (**Figure 7A**). Analysis of GFP signal in coronal sections of the E16 neocortex demonstrated increased branching of neuronal processes in *shEbp1* conditions compared to control, as normal pyramidal neuron projections include a single unbranched axon extending towards basal white matter (WM) tracts, along with an apical dendrite oriented towards the pial surface. Tracing the morphology of transfected neurons (**Figure 7B**) highlighted the impact of Ebp1 depletion on neurite outgrowth at various neurite lengths, with Sholl analysis (**Figures 7C-D**) demonstrating a significantly increased branch number in *shEbp1* conditions – an approximately two-fold increase for proximal segments (**Figure 7C**). Importantly, this increased branching phenotype was rescued by co-electroporation of an Ebp1 overexpression plasmid (*oeEbp1*) along with *shEbp1*, with neuronal morphology tracing and branching analysis quantified as indistinguishable from control conditions.

**Figure 7.**
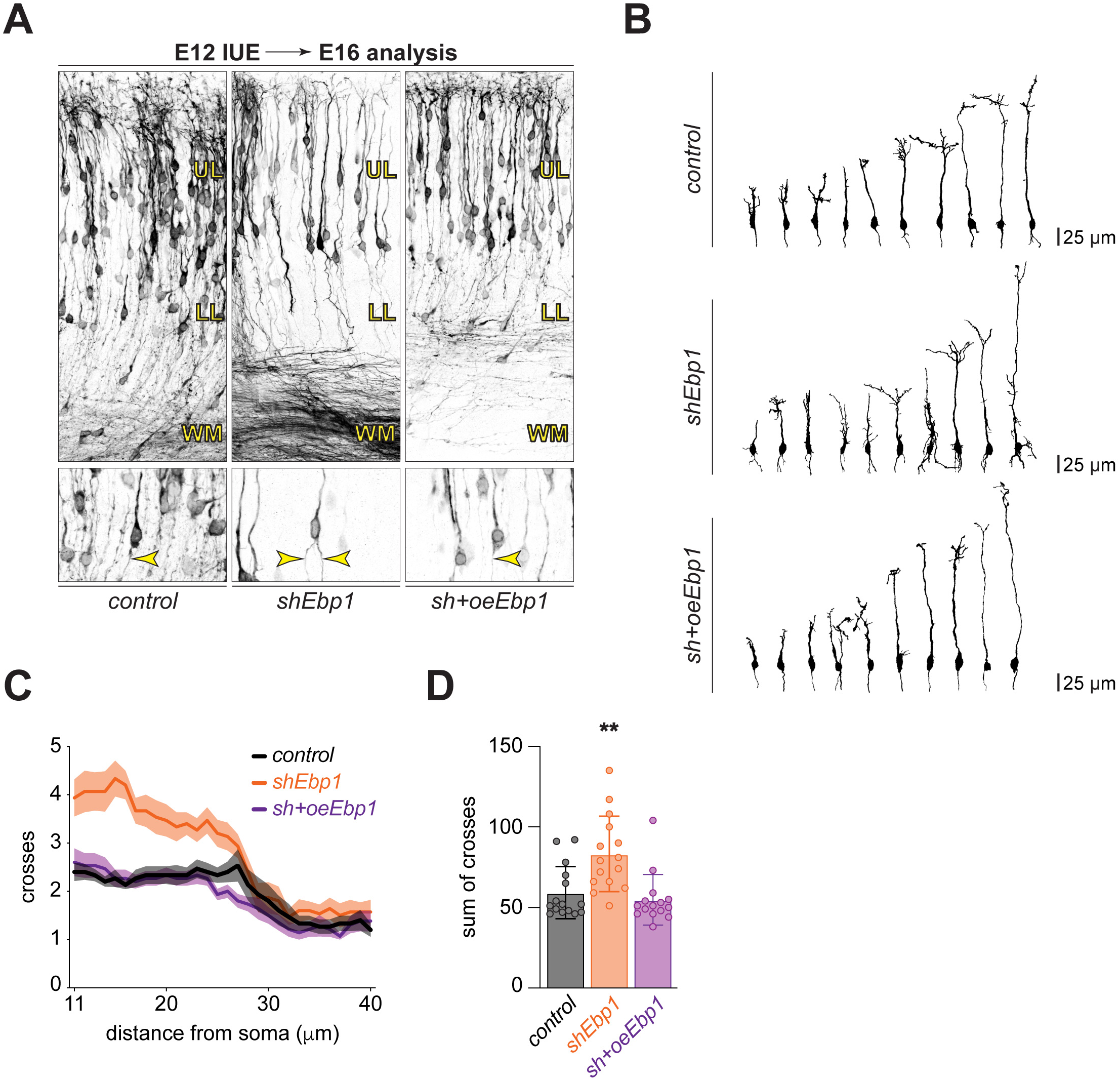
Ebp1 regulates neurite branching of early-born neocortical neurons during development *in vivo*. **(A)** E12 *in utero* electroporation (IUE) followed by analysis at E16, comparing *shEbp1* vs. scrambled shRNA control transfection, and rescue by co-electroporation of *shEbp1* with an Ebp1 overexpression plasmid (*oeEbp1*). Transfected cells are labeled by co-electroporation with CAG-GFP, and analyzed in coronal sections showing the WM to pial surface (top), with individual cells in each condition magnified (bottom), including labeled basally projecting axons (yellow arrows). **(B)** Morphology tracing of individual GFP labeled neurons in control, *shEbp1*, and rescue *sh+oeEbp1* conditions from (A). **(C)** Sholl analysis of (B), comparing branching per unit distance from the soma in control (black, *n* = 15), *shEbp1* (orange, *n* = 15), and rescue *sh+oeEbp1* (purple, *n* = 15) conditions. **(D)** Sum total branches of Sholl analysis in (C). Data in (C) and (D) shown as mean ± s.d. Significance in (D) assessed by one-way ANOVA with Bonferroni corrected *post hoc* test vs. control (\*\**p* < 0.01).

These findings reinforce the impact of Ebp1 on neurite outgrowth during neocortical NSC maturation in neurons, indicating that Ebp1 constrains the overproduction of neuronal processes, possibly through its maintenance of neurite associated proteostasis.

## DISCUSSION

Taken together, this study analyzes the architecture of protein synthesis in the developing neocortex at high resolution, positioning Ebp1 among 60S TE cofactors to fine-tune neuronal proteostasis in the molecular specification of morphology during neural stem cell differentiation. With a multidisciplinary approach, we demonstrate that Ebp1 is a chief component – rather than a niche regulator – of the protein synthesis machinery in neocortical development. Ebp1 expression is cell-type and temporally specific, with enrichment in the early-born neural stem cell pool, in direct proportion to the transient abundance of ribosomal complexes at this developmental stage. Therefore, Ebp1 is well positioned to shift the balance of proteostasis control on the ribosome surface.

Transcriptional control has been the principal focus in the analysis of gene expression during neocortical neurogenesis (Lein et al., 2017; Silbereis et al., 2016). Recent excellent work has advanced the resolution of this analysis to the single-cell level, with an emerging map of neocortical neural stem cell transcriptional programming coming into focus (Oberst et al., 2019; Telley et al., 2019). However, while these studies have assigned transcriptional signatures to cell subtypes, they also strongly suggest that generic gene expression programs are refined by successive layers of regulation (Cadwell et al., 2019), such as post-transcriptional mechanisms and environmental signals. Neocortical neurogenesis hinges on spatiotemporal gene expression (DeBoer et al., 2013; Kwan et al., 2012; Molyneaux et al., 2007), with the ribosome poised at the final essential step (Kraushar et al., 2016) for precisely timed and targeted protein synthesis (Hanus and Schuman, 2013; Holt and Schuman, 2013; Jung et al., 2014). By visualizing neocortical protein synthesis at near-atomic resolution, we find that the 60S TE is a locus of control in neurogenic gene expression.

The interaction of 60S TE cofactors exists in a dynamic equilibrium, competing for a common binding surface to sculpt protein synthesized by a dynamic macromolecular machine (Balchin et al., 2016; Deuerling et al., 2019). While the regime of *Rpl* and *Rps* mRNA expression appears to follow generally elevated levels in all neocortical NSCs compared to their daughter neurons (**Figure 2A**), there is a great diversity of TE cofactor expression patterns in the developing neocortex (**Figure S13**). Ebp1 is particularly enriched in early-born NSCs, similar to RAC subdomains, but in stark contrast to Metap2, Ltn1, or NAC. Modulating the balance of TE cofactors may be a key determinant of cell type-specific proteostasis, gatekeepers at the very moment a nascent protein emerges from the tunnel.

Our data indicate that Ebp1 participates in high occupancy binding with strong affinity to the 60S TE. This is supported by the abundance of Ebp1 available in the neocortical cytoplasm relative to other TE factors, and permissive binding requirements, including both translating and non-translating ribosomes. Whether Ebp1’s role in active and inactive complexes is linked or distinct remains unclear; for example, Ebp1 binding may protect a reserve of inactive, dormant ribosomes available to participate in translation. Ebp1’s potential interaction with nascent peptide chain and/or recruitment of other ribosome cofactors remains to be established. Since our *ex vivo* cryo-EM analysis of native Ebp1•ribosome complexes includes ribosomes engaging with the entire translated proteome, nascent chain density is highly fragmented in the tunnel, and lacking entirely at the TE vestibule, secondary to heterogeneity intrinsic in the data. Future studies in a more homogenous system will be required to interpret potential Ebp1-nascent chain interactions at high resolution.

The neuronal proteins most impacted by Ebp1 are in cell adhesion, synaptogenic, neuronal migration, and neurite outgrowth pathways, with neocortical Ebp1 knockdown resulting in increased neurite branching. While the mechanism of how Ebp1 knockdown overstimulates neurite branching is not clear, it is likely that maintaining a balance of proteins like Marcks that promote neurite outgrowth (Tanabe et al., 2012; Theis et al., 2013; Weimer et al., 2009; Xu et al., 2014), and Sparc that suppress synapse formation (Kucukdereli et al., 2011; López-Murcia et al., 2015), is required to elaborate appropriate projections and synapses. A balance of protein synthesis and degradation, such as through ubiquitin ligases (Ambrozkiewicz and Kawabe, 2015), is likely to be essential. Ribosomes locally translate mRNAs in neurites (Zappulo et al., 2017) and synaptic compartments (Hafner et al., 2019), including both presynaptic terminals and postsynaptic dendritic spines, providing an immediate and dynamic supply of proteins for synaptic activity and plasticity. Notably, many of the proteins impacted by Ebp1 knockdown are membrane or vesicle associated proteins (**Fig. S14D**), and thus Ebp1 may potentially play a role analogous to NAC in coordinating the subcellular targeting of neuronal protein synthesis (Gamerdinger et al., 2015). The enrichment of Ebp1 along with actively translating ribosomes at the synapse would allow for local proteostasis control, and the subcellular action of Ebp1•ribosome complexes is an interesting direction for future study.

Finally, it will be important to delineate the ribosomal and extra-ribosomal mechanisms of Ebp1 function in the nervous system. Ebp1 deletion restricts growth in mice (Zhang et al., 2008) and *Arabidopsis* (Horváth et al., 2006; Li et al., 2018). Ebp1 influences gene expression in stem cells of the neuroectoderm lineage (Somanath et al., 2018), and helps specify the neural border zone, neural crest, and cranial placode domains in *Xenopus* (Neilson et al., 2017). In *Drosophila*, overexpression of the Ebp1 homolog CG10576 results in ectopic neurogenic-like patches in muscle tissue (Bidet et al., 2003). The nervous system is uniquely sensitive to fluctuations in its proteome, and likewise particularly susceptible to abnormal proteostasis pathology in neurodevelopmental and neurodegenerative disease (Hipp et al., 2019; Jayaraj et al., 2019; Sossin and Costa-Mattioli, 2018). How Ebp1 and the dynamic architecture of ribosomal complexes at the 60S TE contribute to both nervous system development and dysfunction as gatekeepers of functional gene expression is an interesting direction for future study.

## ACKNOWLEDGMENTS

We apologize to the authors of key papers who we could not cite due to space limitations. We are particularly grateful to Rainer Nikolay, Anett Unbehaun, Tatyana Budkevich, Justus Loerke, and Dennis Kwiatkowski for fruitful scientific discussions and technical support. We thank Ludovic Telley (Denis Jabaudon lab) for technical support with the scRNAseq data. M.L.K. would like to thank James Millonig and Daniel Mehan of the Rutgers-RWJMS-Princeton Universities MD/PhD Program for encouraging this international postdoctoral research project during his training. M.L.K. was supported by an EMBO Long-Term Postdoctoral Fellowship (190-2016), Alexander von Humboldt Foundation Postdoctoral Fellowship, and a NIH NRSA F30 MD/PhD Fellowship (1F30MH106220). The study was further supported by funding from the Deutsche Forschungsgemeinschaft (DFG; SFB-740) to C.M.T.S. and T.M. High resolution cryo-electron microscopy data collection was performed with support from an iNEXT Cryo-electron Microscopy Instrumentation Grant (PID:2227; EMBL Heidelberg, special thanks to Wim Hagen and Felix Weis), and an Instruct Structural Biology Pilot R&D Grant (APPID: 2016-232; Diamond Light Source Oxfordshire, special thanks to Jason van Rooyen) awarded to M.L.K. Funding from a NeuroCure/Charité Cluster of Excellence Innovation Project Grant awarded to M.L.K. and C.M.T.S. further supported this work. This work was also supported by funding from the German Research Foundation (Grant #EXC 257 to I.V.).

## AUTHOR CONTRIBUTIONS

M.L.K. designed and conducted the study. C.M.T.S. supervised the study, with contributions from M.S., I.V., V.T., T.M., M.L., and D.B. M.C.A. and E.B. performed *in utero* electroporation experiments. P.T. and T.S. (Schaub) prepared pSILAC and BONCAT samples. K.I. and C.H.V.-V. processed, measured, and analyzed mass spectrometry samples. U.Z. processed and measured RNA sequencing samples. D.H. further analyzed mass spectrometry and RNA sequencing data. M.L.K. prepared, and J.B. froze cryo-electron microscopy samples. J.B. and T.M. performed initial cryo-electron microscopy data collection. F.K. and M.L.K. performed multiparticle cryo-electron microscopy sorting and refinement, T.S. (Sprink) modeled the data, and C.M.T.S. and M.L.K. interpreted the data. A.M.W. processed and imaged samples for immuno-electron microscopy. Primary cell culture and immunocytochemistry was performed by P.T. Tissue preparation and immunohistochemistry was performed by M.L.K. H.Y. cloned and purified recombinant Ebp1-His. M.-R.R. provided mice for a pilot study and expertise. M.L.K. wrote the manuscript and prepared the figures, with valuable input from all authors.

## STAR★METHODS

### KEY RESOURCES TABLE

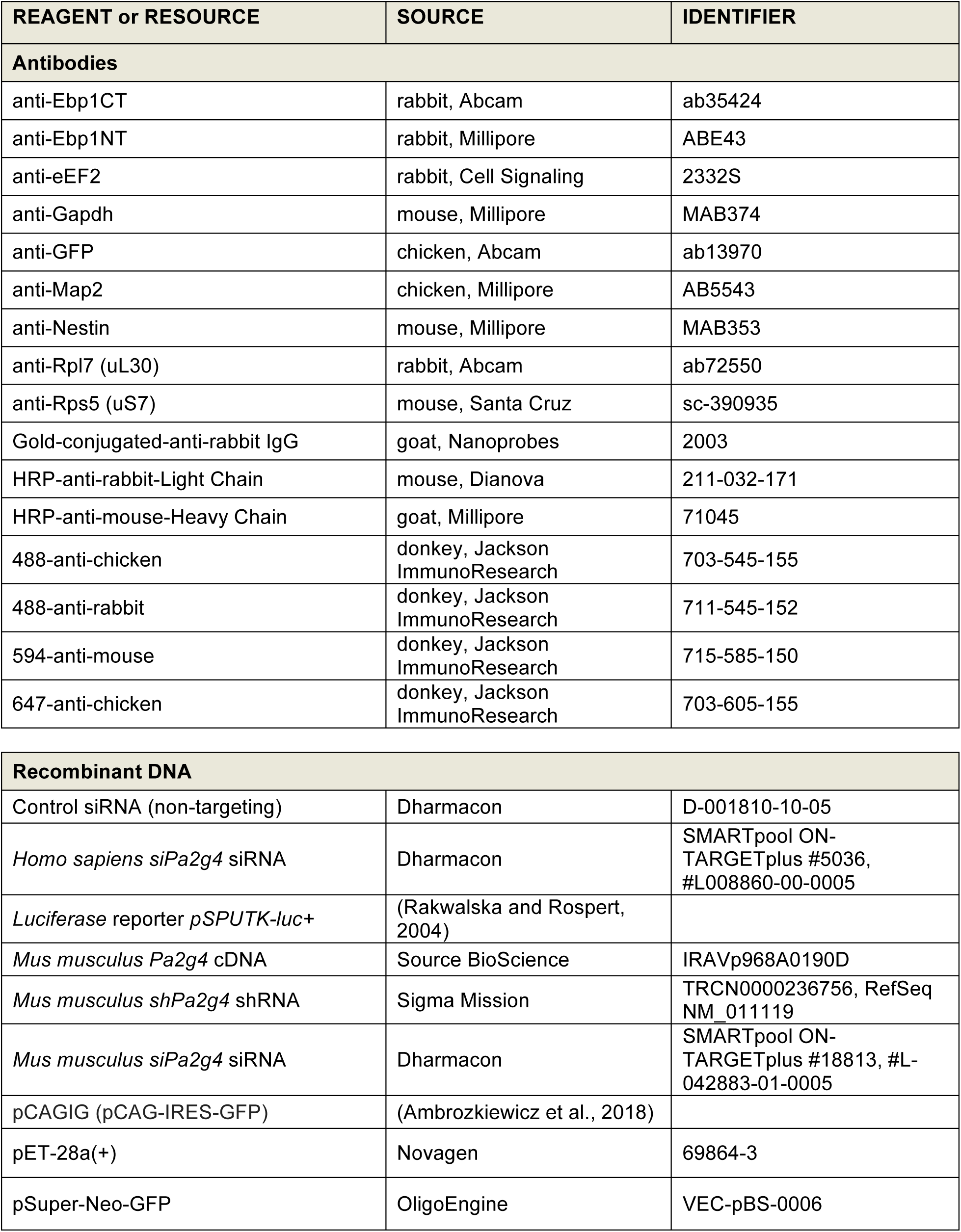
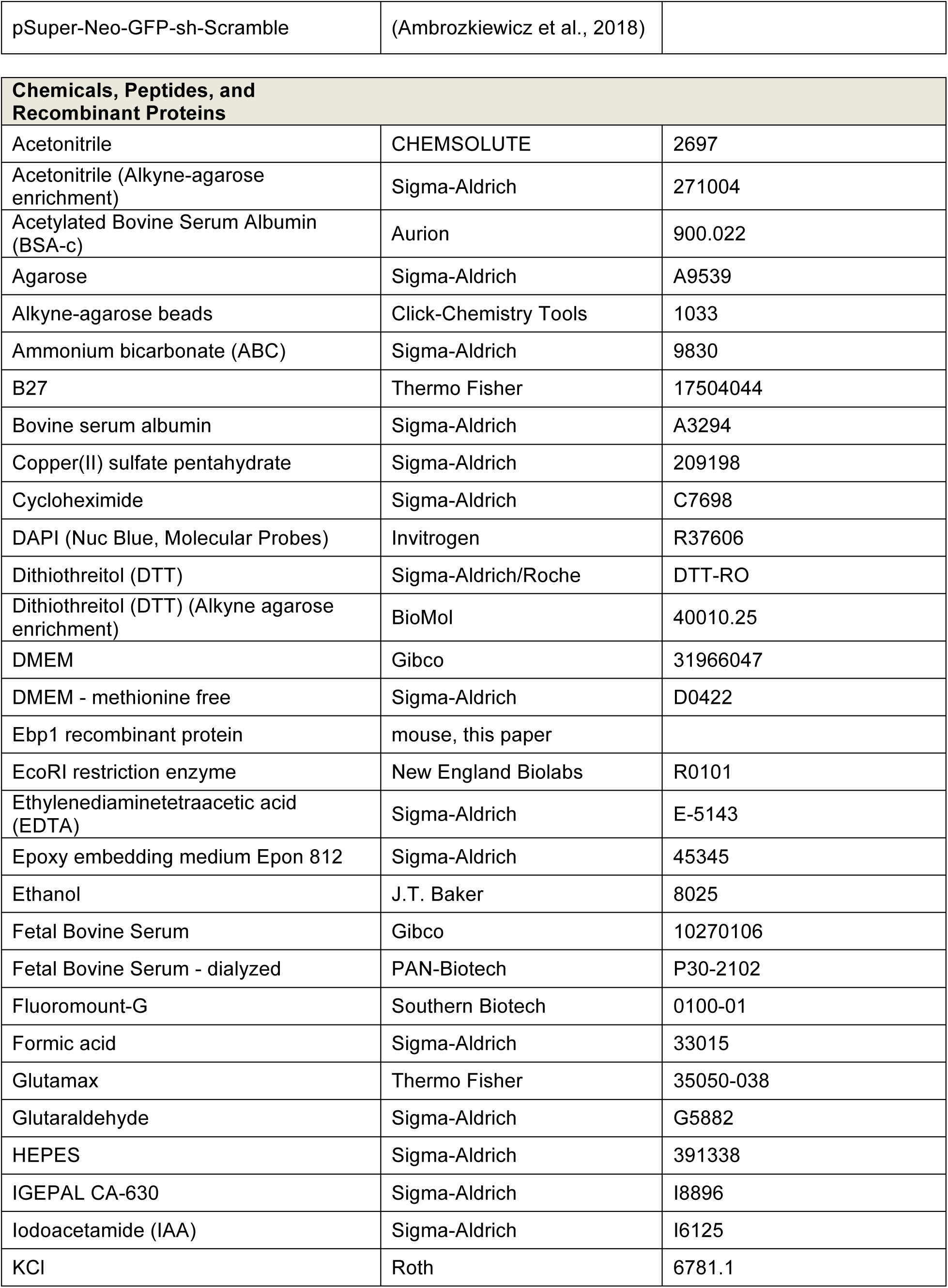
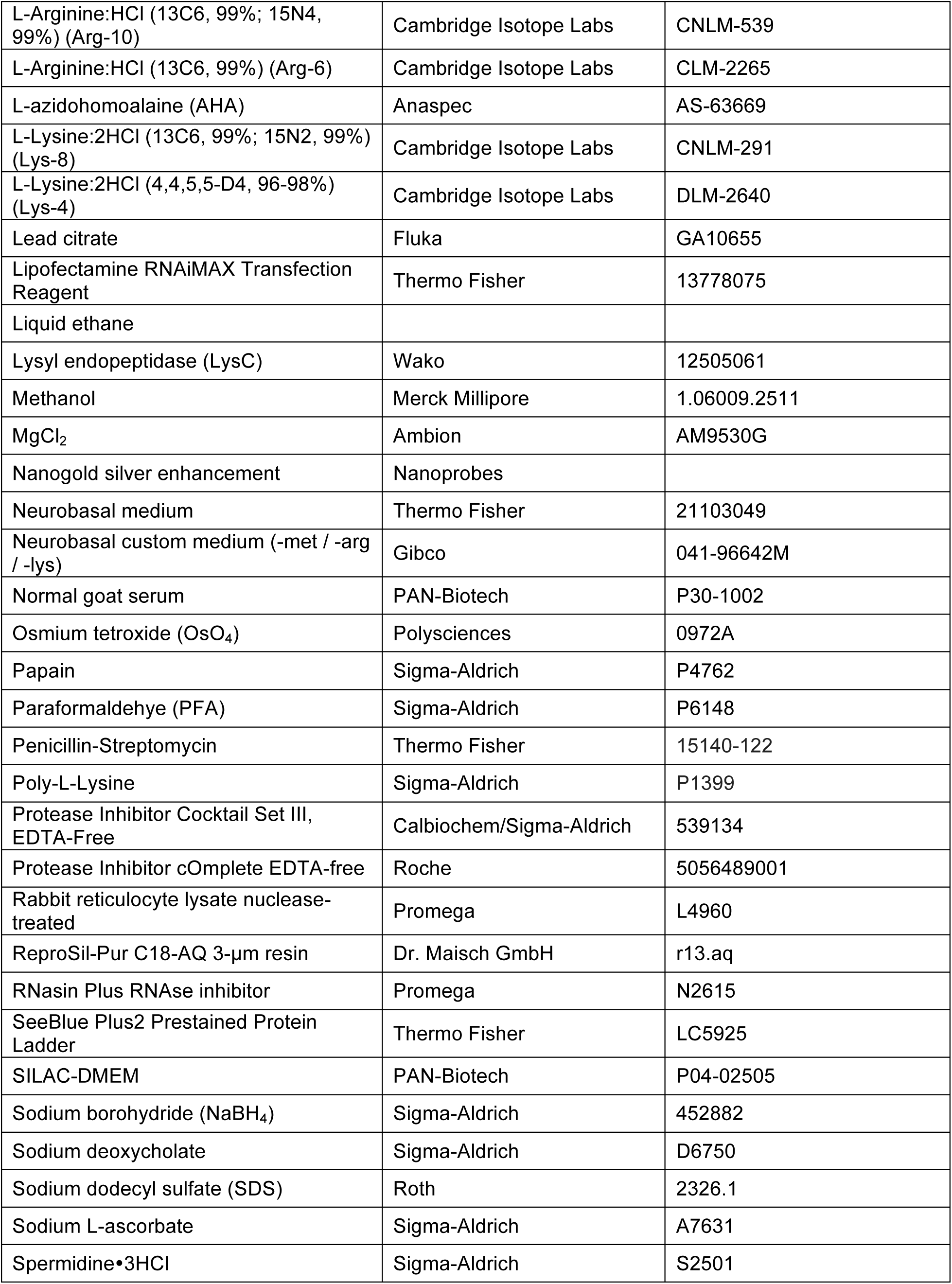
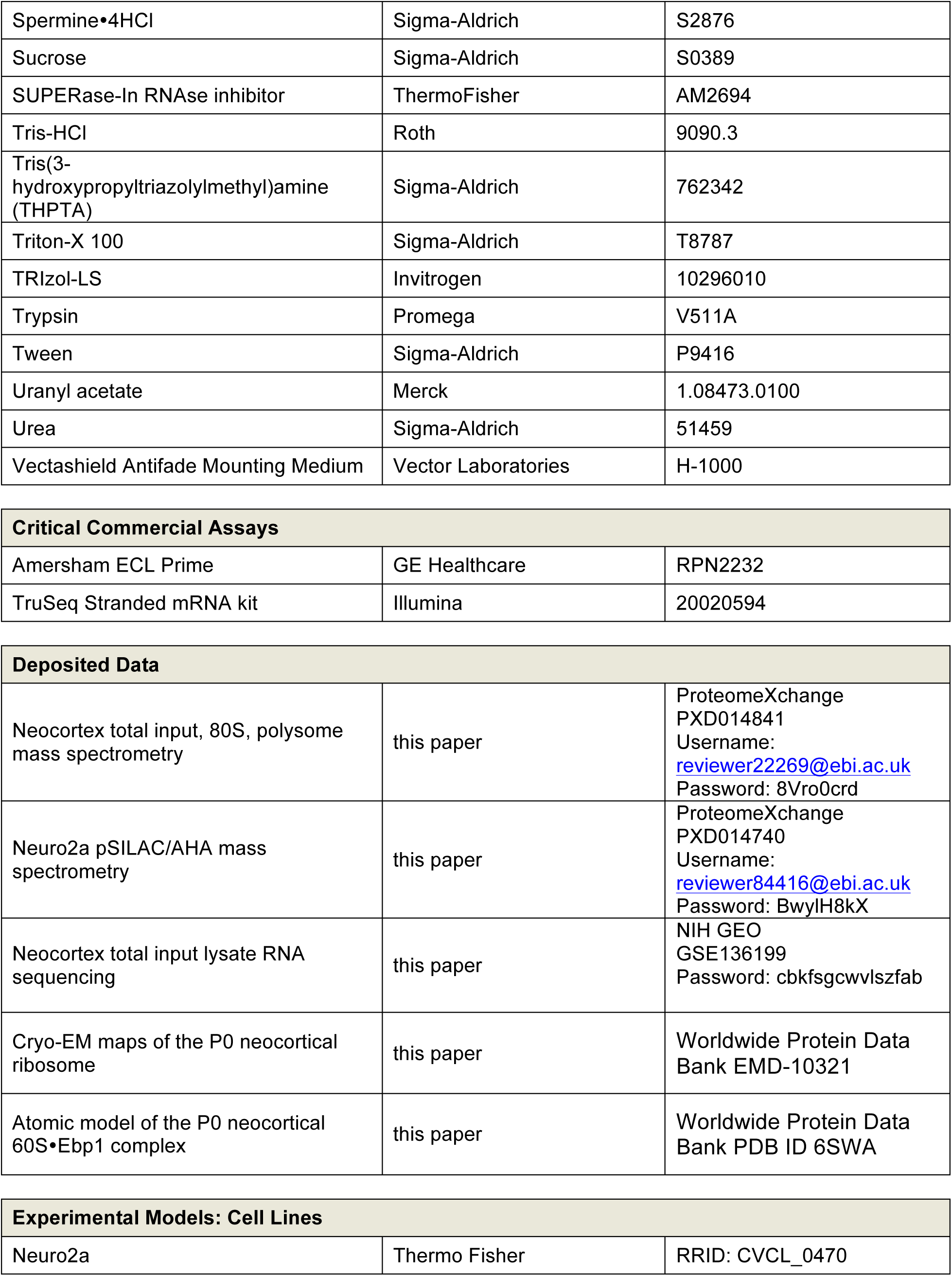
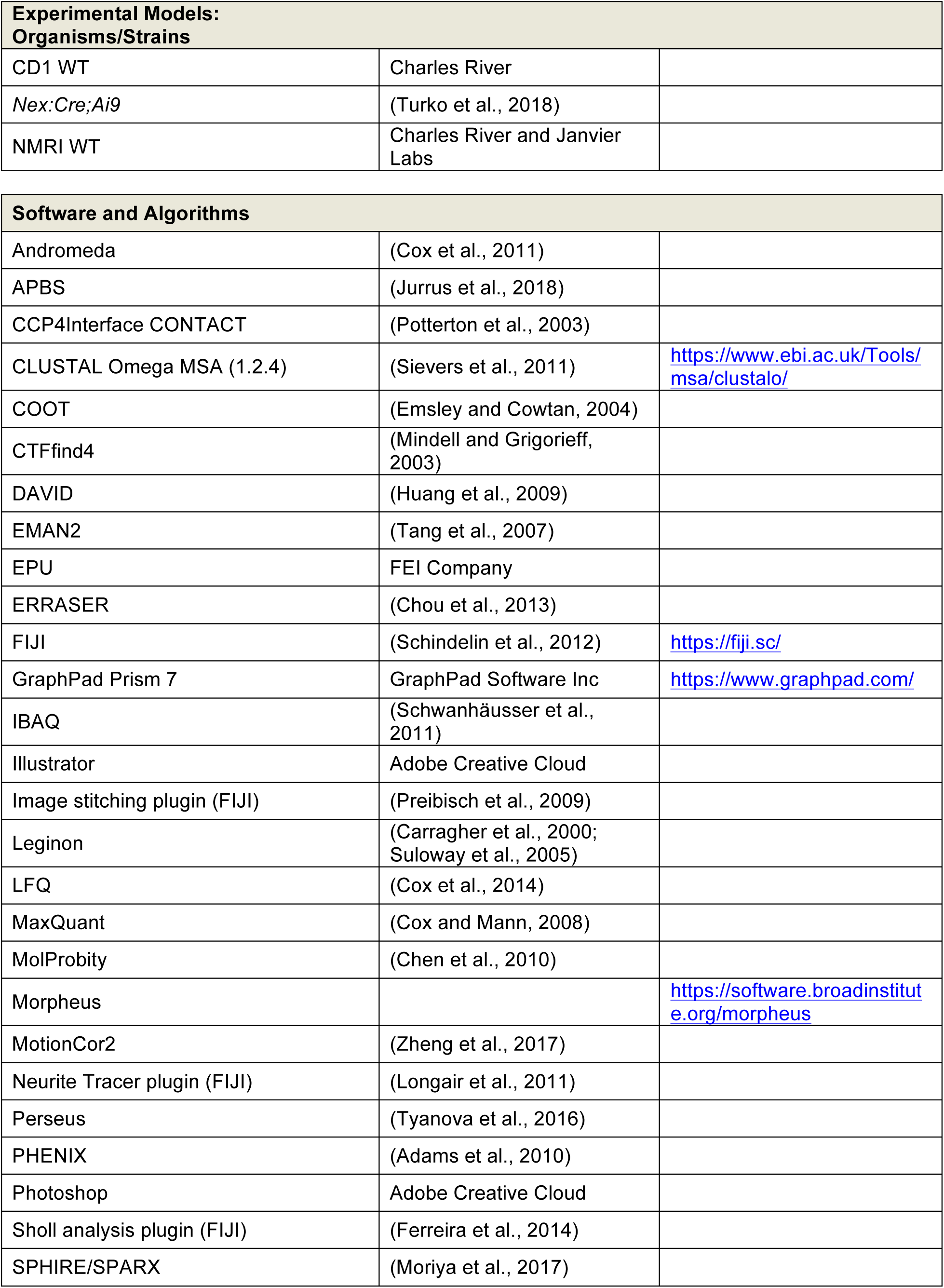
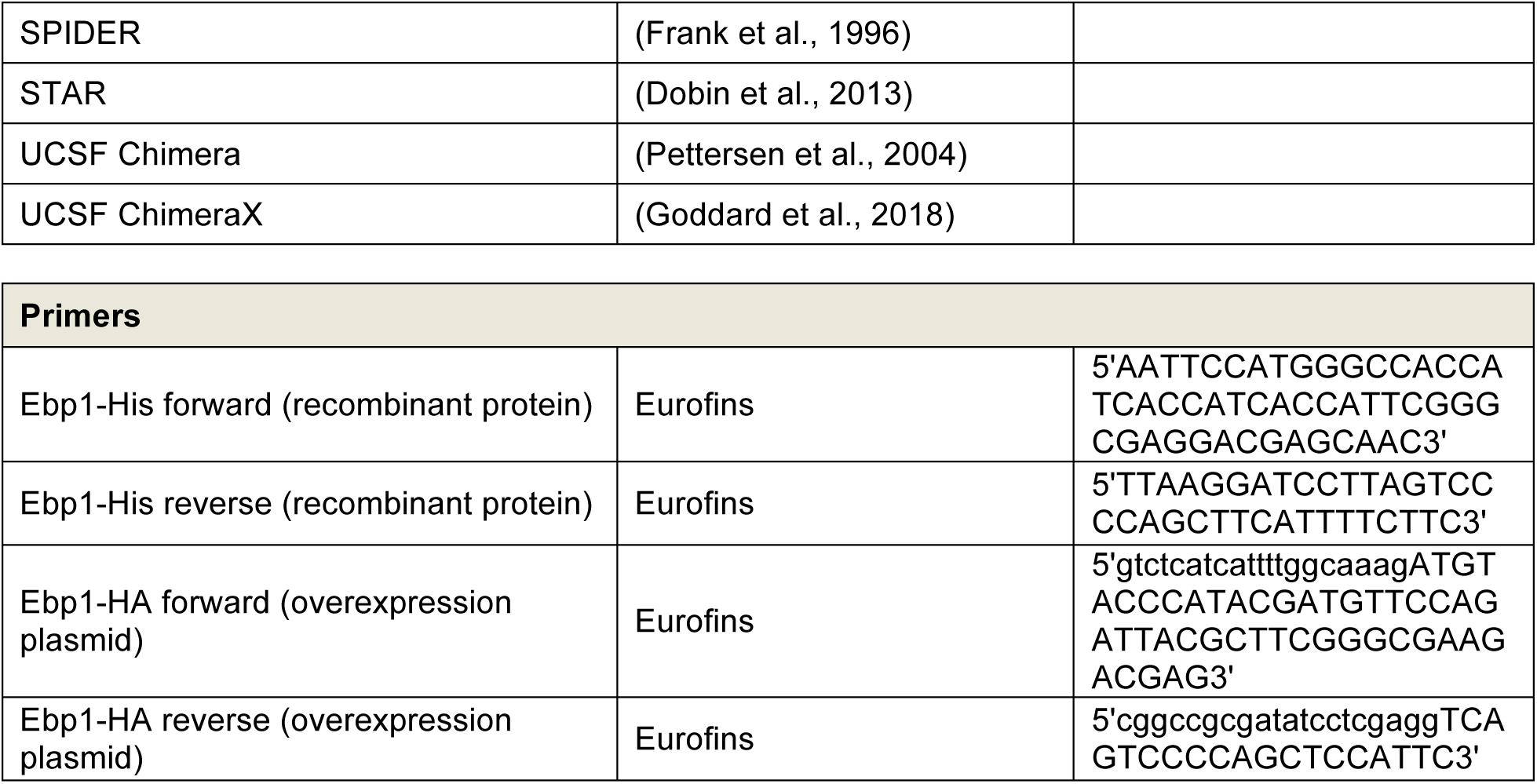

### CONTACT FOR REAGENT AND RESOURCE SHARING

Further information and requests for reagents may be directed to and will be fulfilled by the Lead Contact, (C.M.T.S.).

### EXPERIMENTAL MODEL AND SUBJECT DETAILS

#### Mice

All experiments and associated procedures involving animals in this study were conducted in compliance with the welfare guidelines of the Landesamt für Gesundheit und Soziales (LAGeSo) Berlin and Charité Universitätsmedizin Berlin under certified protocols (Spahn Lab: T0267/15; Vida Lab: T0215/11; Tarabykin Lab: G00206/16, G0054/19), and the Rutgers-Robert Wood Johnson Medical School Institutional Animal Care and Use Committee (IACUC) (Rasin Lab: I12-065-10). Mice were utilized in the embryonic (E12.5, E14, E15.5, E17) and early post-natal (P0) period, inclusive of both sexes in each litter without discrimination, towards the aim of studying common developmental mechanisms. Timed pregnant wild-type (WT) CD-1 mice were obtained from the Charles River Company and utilized for all experiments, with two exceptions: (1) for primary neocortical cell cultures and immunocytochemistry (**Figure 6F**), homozygous *Nex:Cre* females (C57BL/6) were crossed with hemizygous *Ai9* males (C57BL/6J) to produce *Nex:Cre;Ai9* mice as described previously (Turko et al., 2018), labeling post-mitotic glutamatergic neocortical neurons with tdTomato (protocol T0215/11); (2) for *in utero* electroporation (**Figures 7A-D**), NMRI WT (Charles River and Janvier Labs RRID:IMSR_TAC:nmri) mice were utilized (protocols G00206/16, G0054/19).

#### Cell lines

Mouse neuroblastoma Neuro2a cells were obtained from Thermo Fisher (RRID: CVCL_0470) for pulsed stable isotope labeling by amino acids in cell culture (pSILAC) (Schwanhäusser et al., 2009) and bioorthogonal noncanonical amino acid tagging (BONCAT) (Dieterich et al., 2006) coupled mass spectrometry experiments.

### METHOD DETAILS

#### Neocortex dissection and lysis

For all experiments, embryonic (E12.5, E14, E15.5, E17) and postnatal (P0) mouse neocortices were dissected in a 4**°**C room in ice-cold phosphate buffered saline (PBS; ThermoFisher #14040133), frozen as tissue pellets in 1.5mL tubes on dry ice, and stored at −80**°**C. Once sufficient stocks of tissue were generated, each experiment was performed in biological replicates, such that each replicate incorporated an equivalent number of neocortices pooled from distinct litters of mice to meet the input requirements. Frozen tissue pellets were gently lysed by cryogenic grinding on ice using a P1000 tip in 1.5 mL tubes, similar to prior studies (Kraushar et al., 2014, 2015), but with the following lysis buffer: 20 mM HEPES, 100 mM KCl, 10 mM MgCl_2_, pH 7.4, supplemented with 20 mM Dithiothreitol (DTT), 0.04 mM Spermine, 0.5 mM Spermidine, 1x Protease Inhibitor cOmplete EDTA-free (Roche #05056489001), 200 U/mL SUPERase-In RNAse inhibitor (ThermoFisher #AM2694), 0.3% v/v IGEPAL CA-630 detergent (Sigma #I8896). Tissue lysates were clarified of membranes to post-nuclear, post-mitochondrial supernatants by centrifugation at 16100*g* for 10 minutes at 4**°**C with a benchtop centrifuge, and directly applied to downstream analysis. Ribosomal content was estimated by A260 optical density units (ODU) with a NanoDrop 1000 Spectrophotometer. Two neocortical hemispheres (one brain) yields ∼ 2 ODU at P0, 1 ODU at E15.5, and 0.5 ODU at E12.5.

#### Sucrose density gradient ultracentrifugation fractionation

Sucrose density gradients were prepared in Beckman Coulter Ultra-Clear Tubes; #344057 for preparative 5 mL 10-50% gradients (for mass spectrometry, western blot), #344060 for quantitative/analytic 14 mL 5-45% gradients. Base buffer consisted of 20 mM HEPES, 100 mM KCl, 10 mM MgCl_2_, 20 mM Dithiothreitol (DTT), 0.04 mM Spermine, 0.5 mM Spermidine, 1x Protease Inhibitor cOmplete EDTA-free (Roche #05056489001), 20 U/mL SUPERase-In RNAse inhibitor (ThermoFisher #AM2694), pH 7.4, prepared with either 5 & 45% or 10 & 50% sucrose w/v. Overlaid 5 & 45% or 10 & 50% sucrose-buffer solutions were mixed to linearized gradients with a BioComp Gradient Master 107ip. Neocortical lysates were balanced to equivalent ODU and volume across samples for comparison in analytic gradients, with a minimum of 3 ODU required for each biological replicate in each experiment. Lysates were overlaid on gradients pre-cooled to 4**°**C. 5-45% gradients were centrifuged in a SW40 rotor (Beckman Coulter) for 5 hrs, 4**°**C, 25000 rpm; 10-50% gradients were centrifuged in a SW55 rotor (Beckman Coulter) for 1hr, 4**°**C, 37000 rpm. Gradients were fractionated using a BioComp Piston Gradient Fractionator and Pharmacia LKB SuperFrac, with real-time A260 measurement by an LKB 22238 Uvicord SII UV detector recorded using an ADC-16 PicoLogger and associated PicoLogger software. Collected samples were stored at −80**°**C for downstream analysis. Notably, with the gentle lysis technique described in the above Methods (0.3% v/v IGEPAL CA-630 detergent), only cytoplasmic mature ribosomal subunits and complexes were measured and fractionated. Analytic gradient area-under-the-curve analysis for 40S-60S, 80S, and polysome peaks was calculated with a Reimann sum, and significance testing by ANOVA with Dunnett’s *post hoc* test performed in GraphPad Prism software (https://www.graphpad.com/scientific-software/prism/), with *p*<0.05 considered significant.

#### Mass spectrometry analysis of total input, 80S, and polysomes

##### Sample preparation

Samples were prepared in biological triplicate, with each sample including mass spectrometry (MS) analysis of total input neocortex lysate, and purified corresponding 80S and polysomes by preparative 10-50% sucrose density gradient ultracentrifugation (**Figure S1**) as described in the Methods above. Notably, with the lysis method described above, only post-nuclear, post-mitochondrial, cytoplasmic mature ribosomal subunits and complexes were measured and fractionated. Each biological replicate incorporated 12 neocortices (6 animals) at P0, 18 neocortices (9 animals) at E17, 24 neocortices (12 animals) at E15.5, 30 neocortices (15 animals) at E14, and 36 neocortices (18 animals) at E12.5. Tissues were pooled such that each biological replicate included an equal number of neocortices derived from multiple distinct litters of embyros/pups.

Samples were processed essentially as described previously (Imami et al., 2018). Briefly, proteins were precipitated from input lysates, or directly from sucrose gradient fractions, with ethanol, then resuspended in 50 µL of 8 M urea and 0.1 M Tris-HCl, pH 8. Proteins were then reduced with 10 mM dithiothreitol (DTT) at room temperature for 30 min, and alkylated with 50 mM iodoacetamide (IAA) at room temperature for 30 min in the dark room. Protein digestion was first performed with lysyl endopeptidase (LysC) (Wako) at a protein-to-LysC ratio of 100:1 (w/w) at room temperature for 3 hrs. Then, the sample solution was diluted to final concentration of 2 M urea with 50 mM ammonium bicarbonate (ABC). Trypsin (Promega) digestion was performed at a protein-to-trypsin ratio of 100:1 (w/w) under constant agitation at room temperature for 16 hrs. Peptides were desalted with C18 Stage tips (Rappsilber et al., 2007) prior to LC-MS/MS analysis.

##### NanoLC-MS/MS analysis

Measurements were performed essentially as described previously with minor adjustments. Reversed-phase liquid chromatography was performed by employing an EASY nLC 1000 or 1200 (Thermo Fisher Scientific) using self-made fritless C18 microcolumns (Ishihama et al., 2002) (75 µm ID packed with ReproSil-Pur C18-AQ 3-µm resin, Dr. Maisch GmbH) connected on-line to the electrospray ion source (Proxeon) of a Q Exactive plus (Thermo Fisher Scientific). The mobile phases consisted of (A) 0.1% formic acid and 5% acetonitrile and (B) 0.1% formic acid and 80% acetonitrile. Peptides were eluted from the analytical column at a flow rate of 200 nL/min by altering the gradient: 5-6% B in 2 min, 6-8% B in 18 min, 8-20% B in 80 min, 20-33% in 80 min, 33-45% B in 20 min, 45-60% B in 2 min, 60-95% B in 1 min. The Q Exactive plus instrument was operated in the data dependent mode with a full scan in the Orbitrap followed by top 10 MS/MS scans using higher-energy collision dissociation (HCD). The full scans were performed with a resolution of 70,000, a target value of 3×10^6^ ions and a maximum injection time of 20ms. The MS/MS scans were performed with a 17,500 resolution, a 1×10^6^ target value, and a 60 ms maximum injection time. The isolation window was set to 2 and normalized collision energy was 26. Ions with an unassigned charge state and singly charged ions were rejected. Former target ions selected for MS/MS were dynamically excluded for 30 s.

##### Processing of mass spectrometry data

All raw data were analyzed and processed by MaxQuant (v1.5.1.2) (Cox and Mann, 2008). Default settings were kept except that ‘match between runs’ was turned on. Search parameters included two missed cleavage sites, cysteine carbamidomethyl fixed modification and variable modifications including methionine oxidation, protein N-terminal acetylation and deamidation of glutamine and asparagine. The peptide mass tolerance was 6ppm and the MS/MS tolerance was 20ppm. Minimal peptide length of 7 amino acids was required. Database search was performed with Andromeda (Cox and Mann, 2008; Cox et al., 2011) against the UniProt/SwissProt mouse database (downloaded 11/2014) with common serum contaminants and enzyme sequences. The false discovery rate (FDR) was set to 1% at peptide spectrum match (PSM) level and at protein level. Protein quantification across samples was performed using the label-free quantification (LFQ) algorithm (Cox et al., 2014). A minimum peptide count required for LFQ protein quantification was set to two. Only proteins quantified in at least two out of the three biological replicates were considered for further analyses. LFQ intensities were log_2_-transformed and imputation for missing values was performed in Perseus (Tyanova et al., 2016) software based on a simulated normal distribution to represent low abundance values below the noise level (generated at 1.8 standard deviations of the total intensity distribution, subtracted from the mean, and a width of 0.3 standard deviations). Hierarchical clustering of the input, 80S, and polysome data for ANOVA significant proteins (FDR = 0.05) in Morpheus (https://software.broadinstitute.org/morpheus) is shown in **Figure S2**, with clustering based on one minus Pearson correlation using an average linkage method. Proteins whose abundance differed significantly among developmental stages were identified by multiple sample ANOVA test at a permutation-based FDR cutoff of 0.05. Log_2_ LFQ intensities were further z-transformed for only significantly changed proteins.

To estimate protein abundance within input and ribosome fractions, the intensity-based absolute quantification (iBAQ) algorithm was used, which computes the sum of all the peptides’ intensities divided by the number of theoretically observable peptides. Stoichoimetry matrices (**Figures 1C and S3C-D**) compared the median iBAQ value across replicates for each gene, fraction, and time point, plotting the log_2_ transformed ratio for every pair of proteins as a heat map. Mass spectrometry proteomics data have been deposited to the ProteomeXchange Consortium (Vizcaíno et al., 2014) (http://proteomecentral.proteomexchange.org) via the PRIDE partner repository for reviewer access, and for immediate release on publication:

ProteomeXchange PXD014841

Username

Password: 8Vro0crd

#### Single-cell RNA sequencing analysis derived from (Telley et al., 2019)

The transcriptional birthdate and differentiation maps for individual genes (**Figures 2A** and **S13**) were acquired from the open source website associated with (Telley et al., 2019) (http://genebrowser.unige.ch/telagirdon/#query_the_atlas). Averaging the data across all *Rpl* and *Rps* mRNAs into combined single maps for these gene families (**Figure 2A**) was performed with the kind support of Ludovic Telley (Denis Jabaudon lab).

#### RNA sequencing analysis of total neocortical lysates

Total RNA was isolated from post-nuclear, post-mitochondrial, total neocortical lysates prepared as described above in biological duplicate, with each replicate including the following number of neocortical hemispheres (animals) at each developmental stage: E12.5, 80 (40); E14, 60 (30); E15.5, 42 (21); E17, 40 (20); P0, 34 (17). Tissues were pooled such that each biological replicate included an equal number of neocortices derived from multiple distinct litters of embyros/pups. RNA was isolated with TRIzol-LS (Invitrogen #10296010), and 1 µg of RNA per sample was used to prepare libraries with the TruSeq Stranded mRNA kit (Illumina #20020594) according to manufactureŕs instructions. Sequencing was performed on a HiSeq4000. Reads were aligned to the mouse M12 genome using the splice aware aligner STAR (Dobin et al., 2013), and GENCODE (Frankish et al., 2019) gene annotation GRCm38.p5. We used the STAR parameters ‘--alignSJoverhangMin 8 --alignSJDBoverhangMin 1 --outFilterMismatchNmax 999 --outFilterMismatchNoverLmax 0.04 --alignIntronMin 20’ and default otherwise. Gene-level counts were produced using the subread package, with duplicates and multi-mappers discarded. TPMs were calculated using the total exon length for each gene. Significantly changing levels over time of Ebp1, or the median value of Rpl, Rps, and translation-associated gene groups, was assessed by one-way ANOVA followed by Bonferroni corrected *post hoc* testing vs. E12.5, with *p*<0.05 considered significant. RNAseq data have been deposited in the NIH Gene Expression Omnibus (GEO) (Edgar et al., 2002) for reviewer access, and for immediate release on publication:

(https://www.ncbi.nlm.nih.gov/geo/query/acc.cgi?acc=GSE136199)

NIH GEO GSE136199

Password: cbkfsgcwvlszfab

#### Western blot

Analysis was performed with the NuPAGE (Invitrogen) Western blot system according to the manufacturer’s protocol, including 4-12% Bis-Tris NuPAGE gels (Invitrogen NP0321BOX, NP0322BOX, NP0323BOX), MES running buffer, and transfer onto nitrocellulose membranes (Amersham Protran 0.45 NC, GE Life Sciences #10600002) with NuPAGE transfer buffer (NP0006) prepared with 10% methanol. All membranes were blocked in phosphate buffered saline with Tween (PBST; 0.5% Tween) prepared with 5% milk (w/v) for 20 minutes at room temperature, followed by overnight incubation with primary antibody at 4**°**C in PBST-5% milk. Primary antibodies: anti-Ebp1^CT^ (rabbit, Abcam #ab35424), anti-Ebp1^NT^ (rabbit, Millipore #ABE43), anti-Gapdh (mouse, Millipore #MAB374), anti-Rpl7/uL30 (rabbit, Abcam #ab72550), anti-Rps5/uS7 (mouse, Santa Cruz # sc-390935). Membranes were then washed in PBST at room temperature, HRP secondary antibodies applied in PBST-5% milk for 1 hour at room temperature, and again washed in PBST before developing (Amersham ECL Prime Western Blotting Detection Reagent, GE Healthcare #RPN2232) and imaging (GE Amersham Imager 600). Secondary antibodies: HRP-anti-rabbit-Light Chain (mouse, Dianova #211-032-171), HRP-anti-mouse-Heavy Chain (goat, Millipore #71045). Importantly, note that HRP-anti-Light Chain secondary antibody was used because probing with HRP-anti-Heavy Chain secondary antibody introduced a non-specific band (∼50 kDa) just above Ebp1 signal (48 kDa), obscuring the interpretation of actual Ebp1 signal. Band molecular weights were compared to the SeeBlue Plus2 Prestained Standard Protein Ladder (Thermo Fisher #LC5925) as shown in each figure. Band signal intensity was measured using GE Amersham Imager 600 software, with significance testing by ANOVA with Dunnett’s *post hoc* test (≥3 comparisons), or two-tailed unpaired t-test (≤2 comparisons), vs. E12.5 with GraphPad Prism software. Western blot signal for endogenous Ebp1 in lysates was compared to full-length recombinant Ebp1 with a N-terminal Histidine tag (Ebp1-His) as a marker, which was cloned in a pET-28a(+) backbone (Novagen #69864-3) and purified as described (Kowalinski et al., 2007).

#### Binding specificity and affinity analysis of Ebp1**•**ribosmal subunits

40S and 60S subunits were purified from mouse neocortex and rabbit reticulocyte lysate (RRL) essentially as described previously (Pisarev et al., 2007). Briefly, 40 frozen P0 neocortices (40 ODU) were lysed as described above, and ribosomes pelleted through a 1 M sucrose cushion in base buffer (20 mM HEPES, 100 mM KCl, 10 mM MgCl_2,_ 42 U/mL SUPERase-In RNAse inhibitor, pH 7.4) in Beckman Coulter Ultra-Clear Tubes (#344057) with a SW55 rotor at 50000 rpm for 5.5 hrs, 4**°**C. 80S ribosome pellets were resuspended in base buffer, and subjected to a puromycin reaction as described (Pisarev et al., 2007) to release 40S and 60S subunits. Subunits were separated on a 10-30% sucrose high-salt gradient (20 mM HEPES, 0.5 M KCl, 10 mM MgCl_2_, 8 U/mL SUPERase-In RNAse inhibitor) prepared as described above, by ultracentrifugation in Beckman Coulter Ultra-Clear Tubes (#344060) with a SW40 rotor at 27000 rpm for 12 hrs, 4**°**C. Subunits were fractionated and collected as described above, and desalted using Amicon Ultra 0.5 mL 100 kDa MWCO spin columns (Millipore/Sigma UFC510024) and reconstituted 1:3 v/v with low salt buffer (20 mM HEPES, 10 mM KCl, 2 mM MgCl_2_). 40S and 60S subunit concentrations were quantified by NanoDrop Spectrophotometer.

Recombinant Ebp1 with an N-terminal Histidine tag (Ebp1-His) was cloned into a pET-28a(+) backbone (Novagen #69864-3) and purified as described (Kowalinski et al., 2007). For Ebp1-His binding to mouse neocortex 40S and 60S subunits (**Figure S6A**), 5 nM and 20 nM of Ebp1-His was reconstituted with 100 nM subunit in 20 mM HEPES, 100 mM KCl, 10 mM MgCl_2_, and incubated for 30 min at 37**°**C. Samples were pelleted through a 15% sucrose cushion containing 20 mM HEPES, 100 mM KCl, 10 mM MgCl_2_, 0.04 mM Spermine, 0.5 mM Spermidine in Beckman Coulter 230 µL Thickwall Polypropylene Tubes (#343621) with a TLA100 rotor at 35000 rpm for 20 hrs at 4**°**C, separating unbound Ebp1-His from pelleted subunits with bound Ebp1-His. Pellets of subunits•Ebp1-His were resuspended in 20 mM HEPES, 100 mM KCl, 10 mM MgCl_2_. Binding was assessed by Western blot loading supernatant and pellet resuspensions of 40S and 60S samples on the same gel, and probing with Ebp1^CT^ (rabbit, Abcam #ab35424), uL30/Rpl7 (rabbit, Abcam #ab72550), and uS7/Rps5 (mouse, Santa Cruz # sc-390935) antibodies on the same membrane.

Rabbit reticulocyte 40S and 60S subunits were purified and reconstituted to 80S ribosomes as described (Pisarev et al., 2007) from RRL (Promega #L4960). Binding of 200 nM Ebp1-His to 100 nM rabbit 40S, 60S, and 80S (**Figure 3A**) was performed as described above. For dose-response binding of Ebp1-His to 60S rabbit subunits (**Figure 3B**), 100 nM 60S was reconstituted with 1:1 serial dilutions (to 0.5x concentrations) of Ebp1-His from 500 nM to 15.625 nM with 20 mM HEPES, 100 mM KCl, 10 mM MgCl_2_, 0.04 mM Spermine, 0.5 mM Spermidine. Binding was assessed by pelleting and Western blot as described above at each dilution in parallel. This was repeated with a different Ebp1-His dose range between 325 nM to nM. Western blot quantification was performed by normalizing Ebp1^CT^ signal to uL30 (Rpl7) signal, subtracting any signal detected in the supernatant, and generating a single dose-response curve including both independent experiments; with the Ebp1-His concentration demonstrating maximum binding (Ebp1^CT^/uL30) in each experiment set to 100%. Curves were fit using the GraphPad Prism software, with the best fit achieved by non-linear one site-specific binding with Hill slope accommodation (**Figures 3C and S6B**).

Binding dynamics of Ebp1 to the rabbit 60S during mRNA translation were assessed by comparing the following mixtures: (1) 100 nM of rabbit 60S with saturating levels of (350 nM) Ebp1-His; (2) endogenous Ebp1 in RRL (∼100 nM ribosomes estimated as described in the Methods below); (3) 350 nM Ebp1-His added to RRL (100 nM ribosomes); (4) endogenous Ebp1 in RRL (100 nM ribosomes) undergoing *in vitro* translation of a *Luciferase* mRNA; (5) the same conditions as (4) but with 0.1 mg/mL cycloheximide (Sigma-Aldrich #C7698) added to stall Luciferase peptide elongation. All the above mixtures were prepared in parallel, and incubated at 30°C for 30 min, 650 rpm – allowing for protein synthesis to occur to completion in (4) and (5). Mixtures were pelleted through a sucrose cushion as described above to separate unbound vs. bound Ebp1, and pellets likewise analyzed by Western blot (**Figure 3D**).

#### Immuno-electron microscopy

Neocortex was dissected at E12.5, E15.5, and P0 at 4°C as described above, and immersion fixed at 4°C in phosphate buffered saline (PBS) containing 4% PFA and 0.1% Glutaraldehyde overnight, followed by 24 hours incubation in 4% PFA-PBS, and finally stored in 1% PFA-PBS. In order to identify the subcellular localization of EBP1 protein in neocortical precursor/stem and neuronal cells at different developmental stages, we performed pre-embedding nanogold-silver enhanced immunolabeling for Ebp1.

Fixed brains were rinsed several times in PBS and sectioned on a Vibratome (Leica VT1000S) at 50-100 µm. Floating sections were washed again in PBS, followed by incubation in 0.1% sodium borohydride (NaBH_4_; Sigma-Aldrich #452882) in PBS for 15 min to inactivate residual aldehyde groups. Sections were then washed with PBS several times until the solution was clear of bubbles. To improve reagent penetration, the sections were then treated with PBS containing 0.05% Triton X-100 for 30 min and then washed 3x with PBS. To avoid nonspecific binding, sections were incubated for 1 hr in blocking solution containing 5% normal goat serum (NGS; PAN Biotech #P30-1002), and 5% bovine serum albumin (BSA; Sigma-Aldrich #A3294) in PBS. All following immuno-incubations were done with gentle agitation, overnight at 4°C.

After blocking, sections were incubated with primary antibodies: rabbit anti-Ebp1^NT^ (rabbit, Millipore #ABE43) or rabbit anti-EBP1^CT^ (rabbit, Abcam #ab35424) diluted in PBS containing 0.5% acetylated BSA (BSA-c, Aurion #900.022). After washes with PBS/BSA-c, sections were incubated in the secondary nanogold conjugated antibody (Nanoprobes #2003) goat anti-rabbit IgG diluted 1:100 in PBS/BSA-c. To remove unbound secondary antibodies, sections were washed thoroughly with PBS/BSA-c and then with PBS. Subsequently, sections were post-fixed with 2% GA in PBS for 2 hrs to crosslink nanogold in the tissue in order to prevent the loss of labeling during subsequent processing. Next, sections were washed several times in PBS and in double distilled water (ddH_2_O) and prepared for silver enhancement according to the manufacturer’s instruction (Nanoprobes). For structural stabilization, section were incubated with buffered 1% osmium tetroxide (OsO_4;_ Polysciences #0972A) for 1 hr and then washed in PBS followed by ddH_2_O. Sections were dehydrated in increasing concentrations of ethanol and flat-embedded in Epoxy embedding medium (Epon 812; Sigma-Aldrich #45345) between two sheets of Aclar film (Plano #10501-10). After resin polymerization at 60°C, small pieces of cortex were dissected, mounted on plastic stubs, and sectioned *en face* into 60-65 nm sections on an Ultramicrotome (Reichert Ultracut S, Leica) and mounted on 200-mesh Formvar-coated nickel grids (Plano #G2710N). Ultrathin sections were finally stained with 2% aqueous uranyl acetate (Merck #1.08473.0100) for 2 min and with lead citrate (Fluka #GA10655) (Reynolds, 1963) for 30 s. Sections were imaged using a Zeiss TEM-912 equipped with a digital camera (Proscan 2K Slow-Scan CCD-Camera, Zeiss). Subcellular profiles of interest were highlighted in the images with pseudo-color in Adobe Photoshop (**Figures 3E and S7**) as detailed in the guide by Eric Jay Miller (http://www.nuance.northwestern.edu/docs/epic-pdf/Basic_Photoshop_for_Electron_Microscopy_06-2015.pdf).

#### Cryo-electron microscopy and data processing

##### Sample and grid preparation

Pooled 80S and polysomal ribosomes were purified *ex vivo* by preparative 10-50% sucrose density gradient ultracentrifugation from dissected frozen P0 mouse neocortex tissue as described above, but with the following adaptations optimizing for cryo-electron microscopy (cryo-EM). Frozen P0 mouse neocortex (32 animals, 64 neocortex hemispheres) were lysed by cryogenic pulverization with 20 mM HEPES, 100 mM KCl, 10 mM MgCl2, pH 7.4, supplemented with 20 mM Dithiothreitol (DTT), 0.04 mM Spermine, 0.5 mM Spermidine, 1x Protease Inhibitor cOmplete EDTA-free (Roche #05056489001), 480 U/mL RNasin Plus RNAse inhibitor (Promega #N2615), 0.3% v/v IGEPAL CA-630 detergent (Sigma #I8896), and 0.1 mg/mL cycloheximide (Sigma-Aldrich #C7698). Lysates were subjected to further passive lysis by incubation for 1 hr on ice to enhance lipid membrane dissociation, followed by lysate clarification as above. 10-50% sucrose gradients in Beckman Coulter Ultra-Clear Tubes (#344057) were prepared with a base buffer of 10 mM HEPES, 50 mM KCl, 5 mM MgCl2, to pH 7.4, supplemented with 20 mM Dithiothreitol (DTT), 0.04 mM Spermine, 0.5 mM Spermidine, 1x Protease Inhibitor cOmplete EDTA-free, 40 U/mL RNasin Plus RNAse inhibitor, and 0.1 mg/mL cycloheximide. Samples were centrifuged in a SW55 rotor for 50 min at 37000 rpm, 4°C. Fractions corresponding to the 80S and polysomal peaks were collected, pooled, and diluted 1:1 v/v with 20 mM HEPES, 100 mM KCl, 10 mM MgCl_2_, pH 7.4, supplemented with 20 mM Dithiothreitol (DTT), 0.04 mM Spermine, 0.5 mM Spermidine, 1x Protease Inhibitor cOmplete EDTA-free, and 0.1 mg/mL cycloheximide to dilute the sucrose concentration to ≤ 20%. Samples were then pelleted by ultracentrifugation in Beckman Coulter Ultra-Clear Tubes (#344057) with a SW55 rotor for 50 min at 37000 rpm, 4°C. Pellets were resuspended in the same dilution buffer, testing for concentration and quality control by negative stain EM with 2% uranyl acetate. Samples were diluted 1:6 with resuspension buffer, and 3.6 µL of sample were applied to glow-discharged holey carbon grids (Quantifoil R3/3 100 Holey Carbon Films; 2 nM carbon; Micro Tools GmbH), blotted with a Vitrobot device (FEI) for 2-4 s at 4°C, and plunged in liquid ethane. Samples were stored in liquid nitrogen until imaging.

##### Cryo-EM data collection

Initial datasets were collected for sample quality control and low-resolution ribosome reconstruction on a 120 keV Tecnai Spirit cryo-EM (FEI; MPI Molecular Genetics, Berlin) equipped with a CMOS camera (TVIPS), with automated Leginon software (Carragher et al., 2000; Suloway et al., 2005). Projection images were then analyzed by 3-D reconstruction and unsupervised classification for intrinsic ribosomal structure heterogeneity *in silico* with SPIDER (Frank et al., 1996) as described previously (Behrmann et al., 2015; Loerke et al., 2010). These data revealed the presence of extra-ribosomal density at the 60S exit tunnel. To validate these findings, an independent biological replicate sample was re-prepared, with new grids frozen, and likewise imaged using the same protocol, yielding identical density at the 60S exit tunnel.

High-resolution data (**Figures 4-5 and S8-S10**) were collected on a 300 keV Titan Krios (FEI; EMBL, Heidelberg) equipped with a Gatan Quantum K2 direct electron detector at a nominal magnification of 31000x, yielding a pixel size of 0.66 Å on the object scale. Movie stacks were collected in super-resolution mode with EPU automated software (FEI) with the following parameters: defocus range of 0.5-2.5 µm, 40 frames per movie, 20 s exposure time, electron dose of 1.589 e/Å^2^/s and a cumulative dose of 31.78 e/Å^2^ per movie.

##### Computational analysis

High-resolution data collection yielded 5379 movies. The movies were aligned and dose-weighted using MotionCor2 (Zheng et al., 2017) and initial estimation of the contrast transfer function (CTF) was performed with the CTFfind4 package (Mindell and Grigorieff, 2003). Resulting micrographs were manually inspected to exclude images with substantial contaminants (typically lipid/membranes) or grid artifacts. Power spectra were manually inspected to exclude images with astigmatic, weak, or poorly defined spectra. The dataset included 4501 micrographs after these quality control steps (84% of total). Ribosomal particle images were identified using the “swarm” function within e2boxer from the EMAN2 software package (Tang et al., 2007). After the manual removal of artifact particle images, the data set contained 208206 particle images (**Figure S8**).

For multiparticle sorting and 3D refinement (**Figure S9**), the SPHIRE package (Moriya et al., 2017) was used for all steps except for 3D classification, which was performed using a python/SPARX-implementation of the incremental k-means algorithm described previously (Loerke et al., 2010). Therein, two modes for classification exist: (1) refinement, either global or local; and (2) focused classification based on a binary mask, defining a region of interest (ROI) (Penczek et al., 2006). Such a focused mask was derived from 3D variability calculations, which visualizes regions of high heterogeneity with the 3D volume (Behrmann et al., 2015).

However, since heterogeneous regions outside the binary mask can influence the classification, a more sensitive approach was implemented. The “nue” mode, named after the hybrid beast in japanase folklore, creates a hybrid map for each class in a simple procedure: a weighted average of all classes is calculated and used as the “outside”. The ROI within the focused mask is extracted for each class and used as the “inside”. Therefore, the focused mask is transformed into a soft mask by adding a smooth falloff at the edges. For each class, the “outside” map is combined with the respective “inside” map, normalized and filtered, forming the “nue”-map for each class. These “nue” maps are then used as references for focused classification. The “nue” maps only differ within the region of interest, reducing the influence of any peripheral variations. A new set of “nue” maps are calculated at the beginning of each iteration. A similar approach was implemented in Frealign/cisTEM, in which the outside area can be filtered or weighted down in order to reduce its influence during the classification (Grant et al., 2018; Grigorieff, 2016; Zhang et al., 2019). A more detailed description of the procedure is in preparation (Krupp F, *et al*. In preparation).

For the initial refinement, particle images were extracted at a box size of 360 pixels with a pixel size of 1.32 Å/px. All particles were aligned using sxmeridien using a filtered 80S yeast ribosome cryo-EM map as a reference. The refinement yielded a consensus map with sub-nanometer global resolution depicting fragmented densities for the small subunit, tRNAs, eEF2, and Ebp1. In order to separate this dataset into homogeneous sub-states, a hierarchical classification scheme (**Figure S9**) was employed as described previously (Behrmann et al., 2015). Three tiers of sorting were performed, whereby large-scale heterogeneity (e.g. subunit rotation) was classified first, before sorting based on more subtle differences (+/-Ebp1).

In the first tier of sorting, particle images and parameters were decimated to 3.96 Å/px at a box size of 120px to minimize computational expense and limit the resolution for classification. This yielded a rotated 80S, classical 80S, and an artifact population, achieved by an incremental K-means procedure using global and local refinement.

In the second tier of sorting, rotated and classical populations were separated and treated independently. Particle images were decimated to 2.64 Å/px at a box size of 180px. Focused classification was performed, since the maps already depicted high-resolution features. A strong signal of 3D variability was detected in the tRNA-binding site and at the eEF2 binding site, and thus focus masks were constructed in order to separate classes with different compositions of tRNA and eEF2. The “rotated”-branch was separated into two classes: (1) +eEF2, and (2) +eEF2 +P/E-tRNA. The “classical”-branch was separated into three classes: (1) +A-tRNA +P-tRNA, (2) +E-tRNA, and (3) empty 80S. However, within these five classes, the Ebp1 density still appeared fragmented, suggesting further heterogeneity in this region. These findings were confirmed by 3D variability calculations.

In the final sorting tier, particle images were separated into these five classes, and decimated to 2.64 Å/px at a box size of 180 px. A focus mask enclosing the Ebp1 region was defined based on the 3D variability of each of the second-tier classes, and used for sorting into Ebp1-positive and Ebp1-negative classes. The results yielded a nearly equal distribution of Ebp1-positive and Ebp1-negative ribosomes in each sub-state, with an overall Ebp1•80S occupancy of 52% in our dataset (**Figure S9**).

Finally, four of these classes were refined at 1.326 Å/px decimation with box size of 360 px, yielding near-atomic global resolution for all ribosomal complexes, and allowing for the building of an atomic model of the mouse neocortical ribosome 60S•Ebp1. Euler distributions and global Fourier Shell Correlations (FSCs) were calculated (**Figures S10A-D**), in addition to the local resolutions of these maps (**Figures S10E-K’**) with SPHIRE. Local resolution for Ebp1 ranges from 4 Å at the rRNA binding site to 6 Å at the solvent-side periphery (**Figures S10I,I’,K,K’**). Ebp1-positive and Ebp1-negative maps yielded similar global and local resolutions from a similar particle number, permitting the use of the Ebp1-negative map as an internal control for the structural interpretation.

Cryo-EM maps for the neocortical 80S•Ebp1 complex, including both the rotated state with eEF2 and the classical state with A/A+P/P tRNAs, are deposited in the Worldwide Protein Data Bank (wwPDB; https://www.wwpdb.org/) with accession code EMD-10321, for immediate release on publication.

#### Model building

Since our focus was the interaction surface of Ebp1 on the mouse neocortical ribosome, we modeled the 60S subunit in complex with Ebp1 in the cryo-EM map. Modeling was performed in density for the rotated sub-state (+) Ebp1, since this map achieved the highest global resolution of 3.1 Å (**Figure S10C**). A 60S model derived from human polysomes (PDB 5AJ0) (Behrmann et al., 2015) was used as a starting model for the ribosomal proteins, and a rabbit 60S model (PDB 6GZ5) (Flis et al., 2018) was the starting model for rRNA. A pre-existing crystallographic model of mouse Ebp1 (PDB 2V6C) (Monie et al., 2007) was utilized to model Ebp1 density, downloaded from the Research Collaboratory for Structural Bioinformatics (RCSB) (Berman et al., 2000) website https://www.rcsb.org/. For all models, an initial rigid body docking was performed in UCSF Chimera v1.10.2 (Pettersen et al., 2004) (http://www.rbvi.ucsf.edu/chimera), with subsequent adjustment within the density performed in COOT (Emsley and Cowtan, 2004). Thereafter, the models were globally optimized by real-space refinement in PHENIX (Adams et al., 2010) and validated with MolProbity (Chen et al., 2010) (**Figure S8**). To prevent over-fitting during refinement, the applied weight was optimized by monitoring correlation of the map versus model in half-sets of the cryo-EM map (Brown et al., 2015; Greber et al., 2014; Sprink et al., 2016), using individually-determined weight factors. rRNA stuctures were further refined with ERRASER (Chou et al., 2013). Molecular graphics and analysis for figure preparation was performed with UCSF Chimera v1.10.2 and UCSF ChimeraX v0.9.0 (Goddard et al., 2018) (https://www.cgl.ucsf.edu/chimerax/). Analysis of atomic interactions between Ebp1 residues and ribosomal proteins/rRNA was aided by the CCP4Interface 7.0.073 (Potterton et al., 2003) CONTACT algorithm to compute atomic distances between the Ebp1 crystallographic model (PDB 2V6C) and modeled ribosomal proteins/rRNA as input. Atomic distances deemed significant and highlighted as electrostatic contacts (**Figures 4E-H**) were between 0.93-3.95Å. Electrostatic potential maps were generated for Ebp1 (PDB 2V6C), Metap2 (PDB 1KQ9), and Arx1 (PDB 5APN) in UCSF Chimera v1.10.2 using the APBS (Jurrus et al., 2018) interface and webserver http://nbcr-222.ucsd.edu/pdb2pqr_2.1.1/.

The neocortical 60S•Ebp1 atomic model is deposited in the wwPDB with accession code PDB ID 6SWA, for immediate release on publication. A PDB validation report is available upon request with the submission of this manuscript.

#### Pulsed SILAC and BONCAT coupled mass spectrometry

##### Knockdown confirmation

The mouse and human *siEbp1* oligos were obtained from the Dharmacon SMARTpool ON-TARGETplus collection (mouse *siPa2g4* #18813, order #L-042883-01-0005; human *siPa2g4* #5036; order #L008860-00-0005) and compared to non-targeting siRNA control (order #D-001810-10-05). Transfection was performed with Lipofectamine RNAiMAX Transfection Reagent (Thermo Fisher #13778075) according to the manufacturer’s protocol. To confirm robust and specific knockdown with the mouse *siEbp1* oligos, Neuro2a cells were treated in parallel with the following conditions in biological duplicate, followed by Western blot analysis of total lysates (**Figure 6A**): (1) mock transfection, (2) control *siRNA*, (3) mouse *siEbp1*, (4) human *siEbp1*, and (5) 1:1 mouse + human *siEbp1*.

##### Sample preparation

For pulsed stable isotope labeling by amino acids in cell culture (pSILAC) (Schwanhäusser et al., 2009) and bioorthogonal noncanonical amino acid tagging (BONCAT) (Dieterich et al., 2006) coupled mass spectrometry (QuaNCAT) (Eichelbaum et al., 2012; Howden et al., 2013) (**Figures 6C-D**), eight 10 cm plates of Neuro2a cells were grown in standard DMEM (Gibco #31966047) with 1% FBS (Gibco #10270106) to a confluence of 50% in humidified 37°C, 5% CO_2_. Then, mouse *siEbp1* vs. control siRNA tranfection was performed with Lipofectamine RNAiMAX Transfection Reagent (Thermo Fisher #13778075) according to the manufacturer’s protocol, in four plates each. The next morning, media was changed in each condition to either heavy SILAC (2 plates; Cambridge Isotope Labs #CNLM-539, CNLM-291) or medium SILAC (2 plates; Cambridge Isotope Labs #CLM-2265, DLM-2640) prepared with DMEM (Pan-Biotech #P04-02505), 1% dialyzed FBS (PAN-Biotech #P30-2102), GlutaMAX (Thermo Fisher #35050-038), and Penicillin-Streptomycin (Thermo Fisher #15140-122) along with repeated application of the siRNAs. Thus, throughout the course of Ebp1 vs. control knockdown, all newly made proteins were labeled with either heavy or light SILAC (pSILAC), i.e. “label swap” biological replicates. After 48 hr, SILAC media and siRNAs were refreshed. After another 24 hr, one heavy SILAC and one medium SILAC plate from each condition were pulsed with 1 mM L-azidohomoalaine (AHA; Anaspec #AS-63669) for four hours in the corresponding SILAC media prepared with methionine-free DMEM (Sigma-Aldrich #D0422) and 1% dialyzed FBS, labeling all newly made proteins during this acute interval with AHA in addition to the original SILAC label. Thus, acutely synthesized proteins at the point of maximal Ebp1 knockdown were labeled with both SILAC and AHA in parallel (pSILAC-AHA).

Media was then gently aspirated from each plate, followed by washing with ice-cold PBS, then scraping cells into 1 mL ice-cold PBS. Samples were lysed by the addition of 50 mM Tris pH 8, 150 mM NaCl, 1% IGEPAL CA-630 detergent (Sigma #I8896), and 0.5% Sodium Deoxycholate, followed by 5 min of boiling, then lysate clarification by centrifugation at 16000x*g* 4°C for 30 min. 10% of each sample was frozen for Western blot confirmation of Ebp1 knockdown. The remaining 90% of samples were then mixed 1:1 as per the following:

(1) Control+Heavy SILAC : *siEbp1*+Medium SILAC
(2) Control+Medium SILAC : *siEbp1*+Heavy SILAC
(3) Control+Heavy SILAC-AHA : *siEbp1*+Medium SILAC-AHA
(4) Control+Medium SILAC-AHA : *siEbp1*+Heavy SILAC-AHA

Mixtures (1) and (2) were combined with nine volumes of ice-cold ethanol, and frozen at −80°C for downstream MS analysis. Mixtures (3) and (4) were subjected to AHA-enrichment.

##### AHA-enrichment

In preparation for on-bead digestion, azide-containing proteins were enriched from Neuro2a cell lysates using alkyne-agarose beads (Click-Chemistry Tools #1033). Alkyne-agarose beads were rinsed 2 times in pure injection grade water (AMPUWA) before use. To facilitate azide-alkyne binding, a 4x-concentrated “click-solution” was prepared in pure water: 0.8 mM Tris(3-hydroxypropyltriazolylmethyl)amine (THPTA, Sigma-Aldrich #762342), 80 mM Sodium L-ascorbate (Sigma-Aldrich #A7631), and 0.8mM Copper(II) sulfate pentahydrate (Sigma-Aldrich #209198). 200 µl of 4x-click-solution and 200 µl of alkyne-agarose beads were first mixed before addition to 400 µl of Neuro2a lysate. To prevent protease degradation of peptides, Protease Inhibitor Cocktail Set III, EDTA-Free (Calbiochem/Sigma-Aldrich #539134) was added to the final solution (1:50 dilution). To allow time for the click reaction to proceed, the click-bead-lysate mix was briefly vortexed (∼8000x*g* for 5 s) before being placed on an orbital shaker maintained in the dark at room temperature. Following 3.5 hrs of incubation, alkyne-agarose beads were briefly centrifuged at 3000x*g* for 2 min, and resuspended in agarose wash buffer (100 mM Tris, 1% SDS, 250 mM NaCl, 5 mM EDTA, pH 8.0) containing dithiothreitol (DTT, 10 mM). To break disulfide bonds, alkyne-agarose beads were incubated with DTT-solution for 20 min at room temperature, then 10 min at 70°C, at 1000 rpm in a thermomixer (Eppendorf). Following DTT treatment, alkyne-agarose beads were resuspended in agarose wash buffer containing 40 mM Iodoacetamide (IAA). For the alkylation of free thiol groups, alkyne-agarose beads were incubated with IAA for 45 min on an orbital shaker maintained in the dark at room temperature. Following incubation, alkyne-agarose beads were washed using a bench top centrifuge (Roth) and 2 ml centrifuge columns (Pierce) with the following solutions, 10 times each: (1) agarose wash buffer, (2) 8 M Urea in 100 mM Tris, and (3) 70% acetonitrile solution (100 mM ammonium bicarbonate buffer; ABC). Following washing, beads were then resuspended in 35% acetonitrile (50 mM ABC buffer) before centrifugation at 3000x*g* for 2 min to form a bead-pellet. The resulting supernatant was removed and the tube containing the pellet was frozen on liquid nitrogen before storage at −20°C until on-bead digestion.

##### Mass spec analysis

Proteins from cell lysates were precipitated in 90% ethanol solution at - 20°C followed by 30 min centrifugation at 20000x*g* at 4°C. Protein pellets and AHA-clicked beads were resuspended in 2 M urea, 6 M Thiourea, 0.1 M Tris pH 8 solution. Proteins were reduced and alkylated with 10 mM DTT and 55 mM iodoacetamide at room temperature, respectively. For lysis, proteins were incubated with lysyl endopeptidase (Wako) at room temperature for 3 hr. Three volumes of 50 mM ammonium bicarbonate solution were added, and proteins were further digested with trypsin (Promega) under constant agitation at room temperature for 16 hr. Peptides were desalted with C18 Stage Tips prior to LC-MS/MS analysis. Peptide concentration was measured based on 280 nm UV light absorbance.

Reversed-phase liquid chromatography was performed employing an EASY nLC II (Thermo Fisher Scientific) using self-made C18 microcolumns (75 µm ID, packed with ReproSil-Pur C18-AQ 1.9 µm resin, Dr. Maisch, Germany) connected on-line to the electrospray ion source (Proxeon, Denmark) of a Q Exactive HF-X mass spectrometer (Thermo Fisher Scientific). Peptides were eluted at a flow rate of 250 nL/min over 1 or 2 hr with a 9% to 55.2% acetonitrile gradient in 0.1% formic acid. Settings for mass spectrometry analysis were as follows: one full scan (resolution, 60,000; m/z, 350-1,800) followed by top 20 MS/MS scans using higher-energy collisional dissociation (resolution, 15,000; AGC target, 1e^5^; max. injection time, 22 ms; isolation width, 1.3 m/z; normalized collision energy, 26). The Q Exactive HF-X instruments was operated in data dependent mode with a full scan in the Orbitrap followed by up to 20 consecutive MS/MS scans. Ions with an unassigned charge state, singly charged ions, and ions with charge state higher than six were rejected. Former target ions selected for MS/MS were dynamically excluded for 20 or 30 s.

All raw files were analyzed with MaxQuant software (v1.6.0.1) with default parameters, and with match between runs and requantify options on. Search parameters included two missed cleavage sites, cysteine carbamidomethyl fixed modification, and variable modifications including methionine oxidation and protein N-terminal acetylation. Peptide mass tolerance was 6ppm and the MS/MS tolerance was 20ppm. Database search was performed with Andromeda against UniProt/Swiss-Prot mouse database (downloaded on January 2019) with common serum and enzyme contaminant sequences. False discovery rate (FDR) was set to 1% at peptide spectrum match (PSM) and protein levels. Minimum peptide count required for protein quantification was set to two. Potential contaminants, reverse database hits and peptides only identified by modification were excluded from analysis. MaxQuant normalized SILAC ratios were used for quantitative data analysis.

We tested for miRNA-like off-target effects using seeds based on the siRNA sequences in the mouse *siEbp1* (*siPa2g4*) knockdown pool with the cWord software (Rasmussen et al., 2013). All proteins from our dataset were ranked according to their change ratio (mean H/M ratio in Forward and Reverse experiments; lowest to highest fold change). UniProt IDs were converted to Ensembl IDs and searched against mouse 3’UTR using cWords (http://servers.binf.ku.dk/cwords/) (Rasmussen et al., 2013). None of the possible 7-mers from these seeds showed specific enrichment in either AHA or pSILAC datasets. A cutoff of >2-fold change in *siEbp1* conditions compared to control in both replicates was considered significant. Gene ontology (GO) pathway analysis was performed with the Database for Annotation, Visualization and Integrated Discovery (DAVID) (Huang et al., 2009) for proteins with >2-fold change from control in *siEbp1* conditions (against all quantified proteins). Importantly, peptides corresponding to Ebp1 measured in *siEbp1* samples did not meet the minimum requirements for quantification, confirming robust knockdown, and therefore show no fold-change ratio. Mass spectrometry proteomics data have been deposited to the ProteomeXchange Consortium (http://proteomecentral.proteomexchange.org) via the PRIDE partner repository for reviewer access, and for immediate release on publication:

ProteomeXchange PXD014740

Username

Password: BwylH8kX

#### Primary neocortical culture and immunocytochemistry

Primary E12.5 neocortical cultures (**Figure 6F**) were prepared from *Nex:Cre;Ai9* animals, as previously described (Turko et al., 2018). Briefly, dissected neocortex tissue was dissociated with Papain for 25 min (1.5 mg/ml) before trituration in bovine serum albumin (10 mg/ml). Cells were then resuspended in Neurobasal (medium, supplemented with 1x B27, 1x Glutamax, and 100 U/ml Penicillin-Streptomycin). Dissociated cells were grown on 12 mm round, glass coverslips coated with Poly-L-Lysine (20 µg/ml) in 24-well plates. Cells were plated in 40 µl droplets at a concentration of 500 cells per µl (total: 20,000 cells per coverslip). Cultures were grown in humidified conditions at 37°C, 5% CO_2_. Cells were cultured for 5 days to allow for neural stem cell (NSC) differentiation into post-mitotic *Nex*-positive neurons.

At days *in vitro* 0, 2, 4 and 5 coverslips were fixed and analyzed by immunocytochemistry for Ebp1 expression in Nestin-positive NSCs and *Nex*-positive neurons as described previously (Turko et al., 2018). In brief, cells were fixed for 15 min in 4% paraformaldehyde (PFA), 4°C solution before subsequent washes in: 0.1 M phosphate buffered solution (PB) and phosphate buffered saline (PBS). All antibodies were diluted (1:1000) in PBS with 0.1% Triton-X 100, and incubated overnight at 4°C on an orbital shaker. Primary antibodies: anti-Nestin (mouse, Millipore #MAB353), anti-Ebp1^NT^ (rabbit, Millipore #ABE43). Secondary antibodies: Alexa Fluor 647-conjugated anti-mouse (goat, Jackson ImmunoResearch) and Alexa Fluor 488-conjugated anti-rabbit (goat, Jackson ImmunoResearch). DAPI was applied to visualize nuclei (NucBlue, Molecular Probes, Invitrogen #R37606). Coverslips were mounted on glass slides using Fluoromount-G (Southern Biotech, #0100-01). Images were captured on an upright confocal microscope (FV-1000, Olympus) using 30x silicon oil-immersion objective (1.05 NA, 0.8 mm WD). Images were analyzed using the FIJI distribution of ImageJ software (Schindelin et al., 2012) (https://fiji.sc/), maintaining constant LUT parameters across images.

#### Immunofluorescence labeling in neocortex tissue

Tissue processing and immunohistochemistry (**Figures 2C and S4**) was performed similar to the previously described method (Kraushar et al., 2014). In brief, embryonic (E12.5, E14, E15.5, E17) and postnatal (P0) mouse brains were dissected at 4°C in ice cold PBS (ThermoFisher #14040133), and initially immersion-fixed with 4% (w/v) paraformaldehyde (PFA) in PBS (PBS-PFA; pH 7.4) at room temperature for 30 min, followed by overnight PBS-PFA fixation at 4°C. Fixed brains were then embedded in 3.2% agarose-PBS, and coronally sectioned at 70 µm on a Leica vibratome (VT1000S). Sections of the anterior sensorimotor neocortex were collected, incubated in blocking solution (PBS, 10% normal donkey serum, 2% w/v BSA, 0.2% w/v glycine, 0.2% w/v lysine), then incubated overnight in probing solution with 0.4% Triton-X and primary antibody at 4°C. Primary antibodies: anti-Map2 (chicken, Millipore #AB5543), anti-Ebp1^CT^ (rabbit, Abcam, #ab35424), anti-Ebp1^NT^ (rabbit, Millipore #ABE43). Samples were washed in PBS, then all secondary antibodies, Alexa 488 anti-rabbit (goat, Jackson ImmunoResearch) and Alexa 647 anti-chicken (goat, Jackson ImmunoResearch), were applied at 1:250 dilution in probing solution for 2 hr at room temperature, washed, incubated with DAPI (NucBlue, Molecular Probes, Invitrogen #R37606) for 10 min, and mounted with Vectashield. Confocal imaging was performed with an upright confocal microscope (FV-1000, Olympus), 20x air objective, maintaining constant parameters and setting across all images. Images were likewise analyzed using FIJI software, including the pairwise stitching plugin (Preibisch et al., 2009), maintaining constant LUT parameters across images.

#### *In utero* electroporation and morphology analysis

The mouse *shEbp1* plasmid was obtained from the Sigma MISSION collection (*shPa2g4*; oligo name: TRCN0000236756; RefSeq NM_011119) in bacterial glycerol stock format, and amplified according to the manufacturer’s protocol, followed by plasmid purification with the Nucleobond Xtra Midi Kit (Macherey & Nagel #740410.100). The non-targeting scrambled shRNA control was generated as described in a prior study (Ambrozkiewicz et al., 2018). The *Ebp1* overexpression plasmid was generated by insert amplification from the Clone IRAVp968A0190D I.M.A.G.E. Fully Sequenced cDNA (Source BioScience) with primers forward 5’-gtctcatcattttggcaaagATGTACCCATACGATGTTCCAGATTACGCTTCGGGCGAAGACGAG-3’ and reverse 5’-cggccgcgatatcctcgaggTCAGTCCCCAGCTCCATTC-3’, followed by cloning into the pCAGIG (pCAG-IRES-GFP) backbone (Ambrozkiewicz et al., 2018) with the restriction enzyme EcoRI (NEB). Co-electroporation of the pCAG-IRES-GFP plasmid was used as a transfection reporter.

E12 *In utero* electroporation (IUE) of control, *shEbp1*, and *shEbp1+oeEbp1* conditions with the CAG-GFP reporter followed by analysis at E16 with confocal imaging, morphology tracing, and Sholl analysis (**Figures 7A-D**) was performed as described (Ambrozkiewicz et al., 2018). Briefly, GFP labeling of electroporated neurons in confocal images was analyzed by morphology tracing with the Neurite Tracer plugin (Longair et al., 2011) by a blinded investigator, followed by the Sholl analysis (Ferreira et al., 2014) plugin run in FIJI with 1 µm radius of concentric circles, plotting the average intersections over distance from the soma and average total summed intersections in each condition. Significance was assessed by one-way ANOVA followed by Bonferroni corrected *post hoc* testing in GraphPad Prism software, with *p* < 0.05 considered significant.

### QUANTIFICATION AND STATISTICAL ANALYSIS

#### Mass spectrometry

MaxQuant software (v1.6.0.1) (Cox and Mann, 2008) was run with default parameters, and with match between runs and requantify options on. Protein quantification across samples was performed using the label-free quantification (LFQ) algorithm (Cox et al., 2014). A minimum peptide count required for LFQ protein quantification was set to two. Only proteins quantified in at least two out of the three biological replicates in input, 80S, and polysome samples, and two out of two (label swap) biological replicates in pSILAC/AHA samples, were considered for further analyses. LFQ intensities were log_2_-transformed and imputation for missing values was performed in Perseus software (Tyanova et al., 2016) based on a simulated normal distribution to represent low abundance values below the noise level (generated at 1.8 standard deviations of the total intensity distribution, subtracted from the mean, and a width of 0.3 standard deviations). For stoichiometry matrices, the IBAQ algorithm (Schwanhäusser et al., 2011) was used to quantify within-sample abundance. Database search was performed with Andromeda (Cox et al., 2011) against UniProt/Swiss-Prot mouse database with common serum and enzyme contaminant sequences. False discovery rate (FDR) was set to 1% at peptide spectrum match (PSM) and protein levels. Minimum peptide count required for protein quantification was set to two. Potential contaminants, reverse database hits and peptides only identified by modification were excluded from analysis.

For input, 80S, and polysome MS performed in biological triplicate, Ebp1 and the median protein abundance within protein groups (Rpl, Rps, translation-associated) were tested for significantly changing levels across developmental stages by one-way ANOVA with Bonferroni corrected *post hoc* testing, with *p*<0.05 considered significant. For pSILAC/AHA MS performed in biological duplicate with SILAC label swap, MaxQuant normalized SILAC ratios were used for quantitative data analysis. All proteins from our dataset were ranked according to their change ratio (mean H/M ratio in Forward and Reverse experiments; lowest to highest fold change). A cutoff of >2-fold change from control in both replicates was considered significant.

#### RNAseq

Samples were prepared in biological duplicate at each developmental stage. Reads were aligned to the mouse M12 genome using the splice aware aligner STAR (Dobin et al., 2013), and GENCODE (Frankish et al., 2019) gene annotation GRCm38.p5. Gene-level counts were produced using the subread package. Significantly changing levels over time of Ebp1, or the median value of Rpl, Rps, and translation-associated gene groups, was assessed by one-way ANOVA followed by Bonferroni corrected *post hoc* testing vs. E12.5, with *p*<0.05 considered significant.

#### Sucrose density gradient ultracentrifugation fractionation curves

Analytic gradient area-under-the-curve (AUC) analysis for 40S-60S, 80S, and polysome peaks was calculated from real-time A260 values measured by PicoLogger recorder and software during sample fractions in biological duplicate or triplicate at each developmental stage. A Reimann sum was used to calculate the AUC corresponding to 40-60S, 80S, and polysome peaks of the gradient. Significance was tested by one-way ANOVA with Dunnett’s *post hoc* test vs. E12.5, performed in GraphPad Prism, with *p*<0.05 considered significant.

#### Quantitative Western blot

Western blot band signal intensity measured in duplicate membranes was quantified using GE Amersham Imager 600 software, with significance testing by two-tailed unpaired t-test (≤2 comparisons), vs. E12.5 with GraphPad Prism software, with *p*<0.05 considered significant.

#### Sholl analysis

Neuronal morphology tracings of GFP signal were generated with the Neurite Tracer plugin (Longair et al., 2011) in FIJI, then analyzed by the Sholl analysis plugin (Ferreira et al., 2014) with 1µm radius of concentric circles, plotting the average intersections over distance from the soma, and average total summed intersections in each condition. Significance of total summed intersections was assessed by one-way ANOVA followed by Bonferroni corrected *post hoc* testing in GraphPad Prism software, with *p*<0.05 considered significant.

### DATA AVAILABILITY

Mass spectrometry proteomics data (**Figures 1B-C, 6C-D, and S2-S3**) have been deposited to the ProteomeXchange Consortium (http://proteomecentral.proteomexchange.org) via the PRIDE partner repository for reviewer access, and for immediate release on publication:

ProteomeXchange PXD014740

Username

Password: BwylH8kX

RNAseq data (**Figure 2B**) have been deposited in the NIH Gene Expression Omnibus (GEO) for reviewer access, and for immediate release on publication: (https://www.ncbi.nlm.nih.gov/geo/query/acc.cgi?acc=GSE136199)

NIH GEO GSE136199

Password: cbkfsgcwvlszfab

Cryo-EM maps for the neocortical 80S•Ebp1 complex (**Figures 4**-**5 and S7-S10**), including both the rotated state with eEF2 and the classical state with A/A+P/P tRNAs, have been deposited in the Worldwide Protein Data Bank (wwPDB; https://www.wwpdb.org/) with accession code EMD-10321, for immediate release on publication.

The neocortical 60S•Ebp1 atomic model (**Figures 4**-**5, S8, and S11**) has been deposited in the wwPDB with accession code PDB ID 6SWA, for immediate release on publication. A PDB validation report is available upon request with the submission of this manuscript.

## SUPPLEMENTARY FIGURE LEGENDS

**Supplementary Figure 1.**
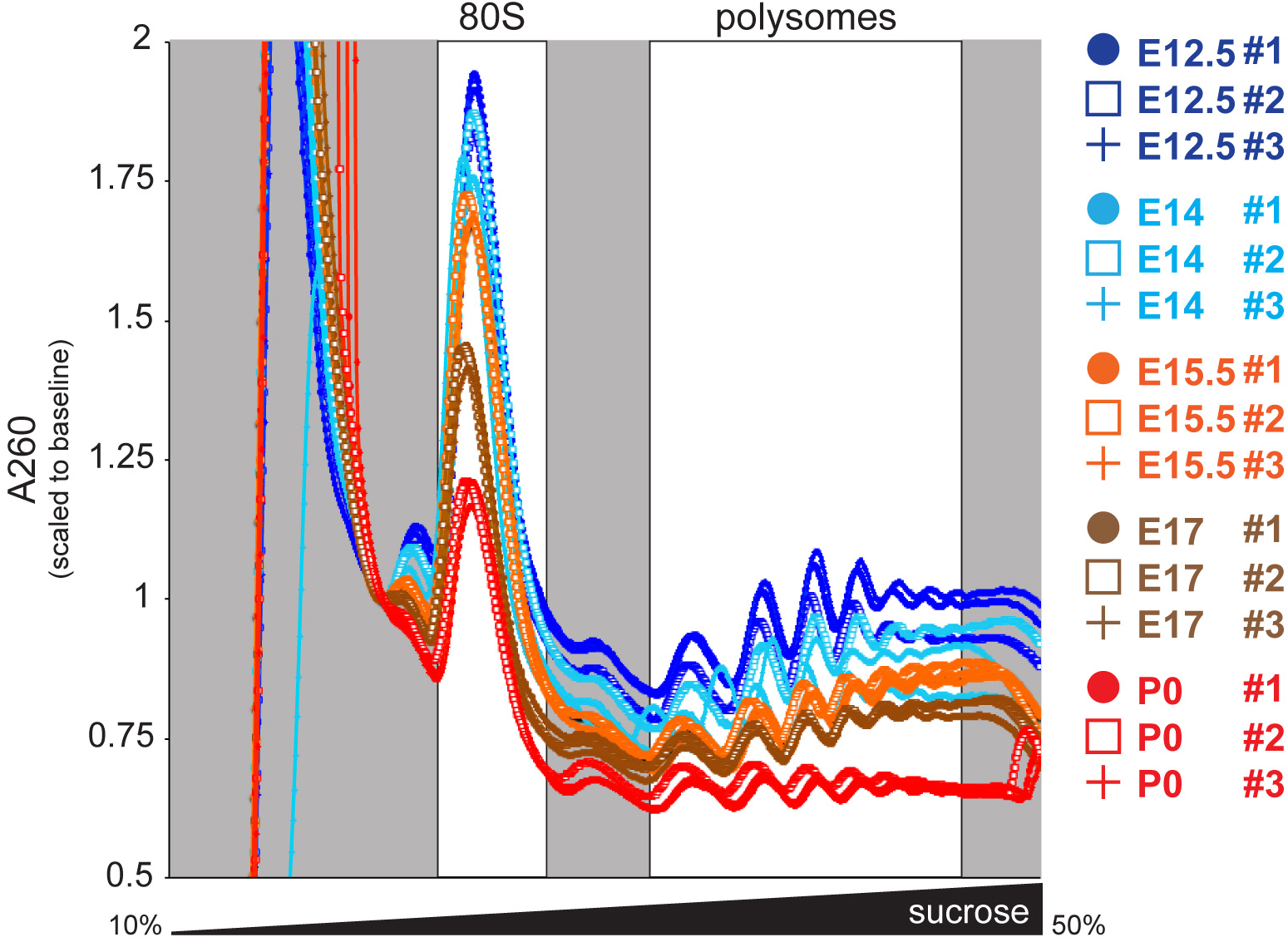
Preparative sucrose density gradient fractionations for MS samples, associated with Figure 1. Mouse neocortical lysates were subjected to preparative sucrose density gradient ultracentrifugation fractionation in biological triplicate to purify 80S and polysomal ribosome complexes at E12.5, E14, E15.5, E17, and P0. Input lysate, pooled 80S fractions, and pooled polysome fractions were analyzed by LC-MS/MS.

**Supplementary Figure 2.**
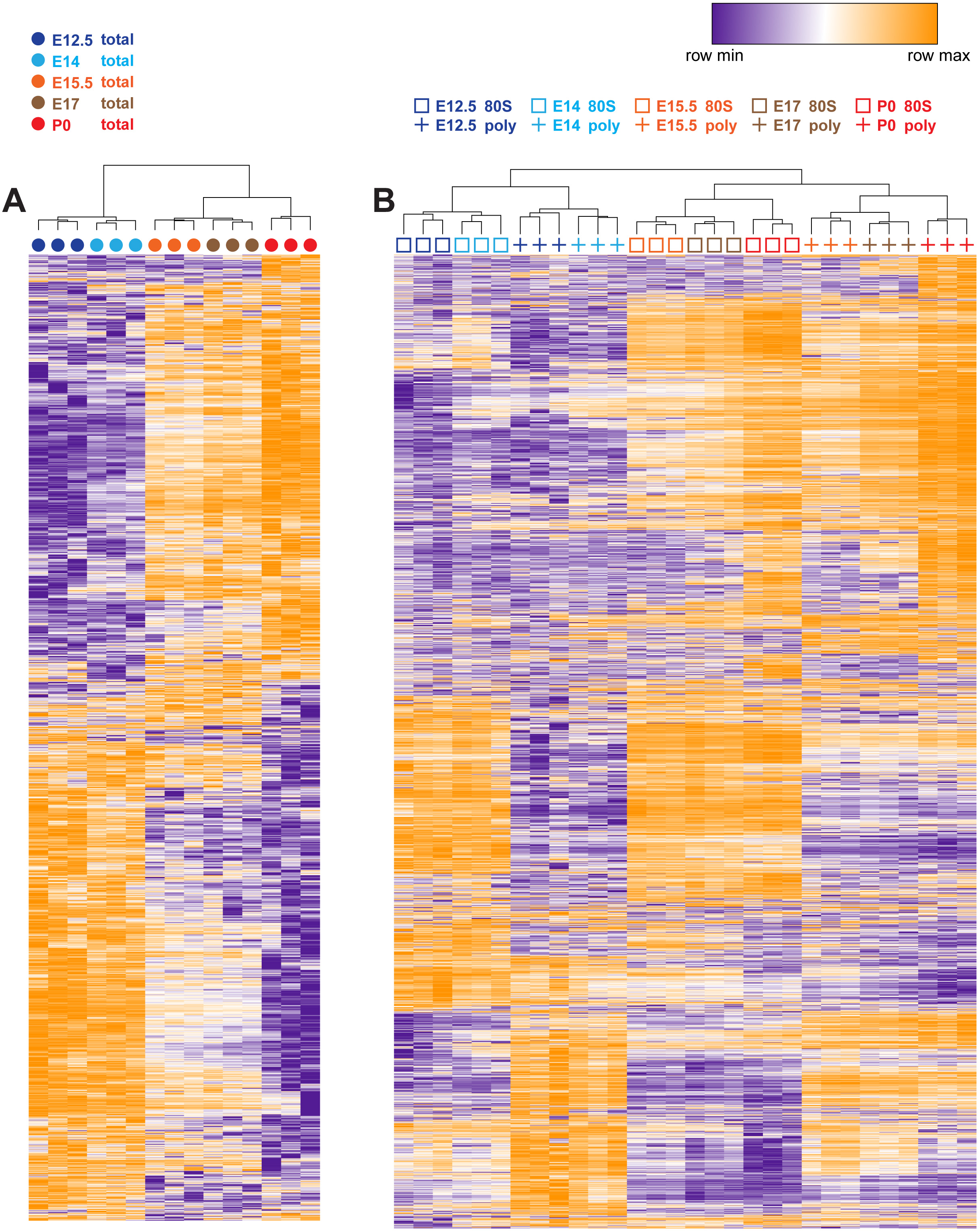
Hierarchical clustering of MS samples, associated with Figure 1. Hierarchical clustering of neocortex MS data for **(A)** total input lysates and **(B)** 80S and polysomes in biological triplicate at E12.5, E14, E15.5, E17, and P0. Clustering of ANOVA significant (FDR = 0.05) proteins based on one minus Pearson correlation with an average linkage method. Heat maps colored by higher (orange) and lower (purple) protein expression per row (protein ID) max and min, respectively. Results demonstrate clustering by replicates, 80S vs. polysomes, and developmental stages.

**Supplementary Figure 3.**
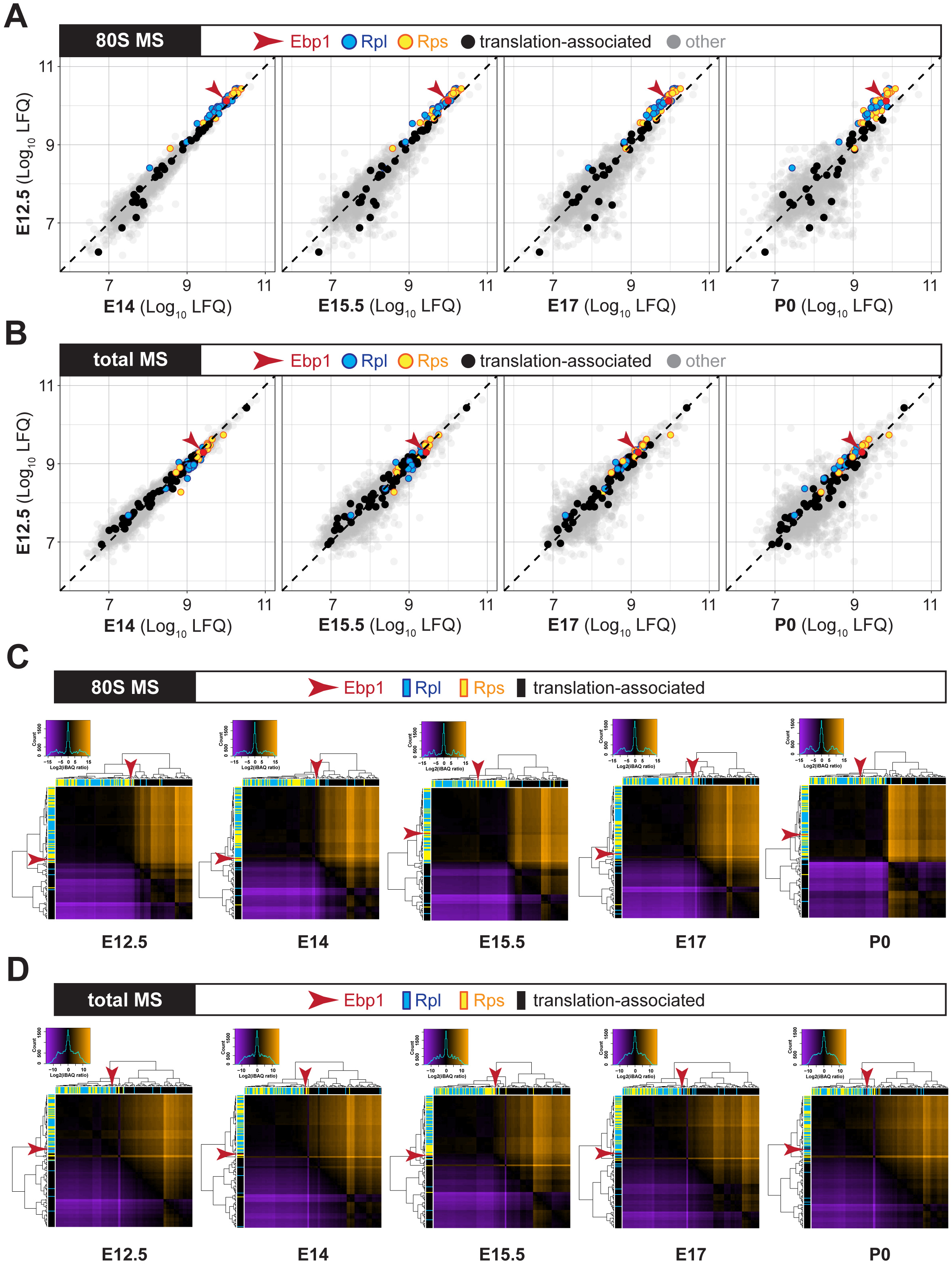
Scatter plots and stoichiometry cluster heat maps for input and 80S proteins, associated with polysome samples in Figures 1B-C. **(A-B)** Scatter plots of ribosomal complex protein enrichment measured by MS comparing early neurogenesis E12.5 vs. each subsequent stage. (A) 80S MS showing Ebp1 (red arrow) enriched among ribosomal proteins (RPs) of the large (Rpl, blue) and small (Rps, yellow) subunits, similar to the polysome MS (Figure 1B). Ebp1 enrichment is shown relative to the level of translation-associated proteins (black), and all other proteins (grey). (B) Input MS, showing Ebp1 is among the most abundant proteins in the neocortex. **(C-D)** Stoichiometry cluster heat maps comparing the enrichment of each RP (Rpl, blue; Rps, yellow), translation-associated proteins (black), and Ebp1 (red arrow) in comparison to every other protein per developmental stage in the (C) 80S MS and (D) total input MS, corresponding to the polysome MS in Figure 1C. The expression of adjacent proteins on the x-axis is shown as higher (orange), lower (purple), or similar (black) relative to each protein on the y-axis (legend and histogram at top left for each stage). Ebp1 is expressed at levels comparable to RPs in neocortical total lysates, and is nearly stoichiometric to RPs in the 80S, similar to polysomes. In contrast, other translation-associated proteins are sub-stoichiometric.

**Supplementary Figure 4.**
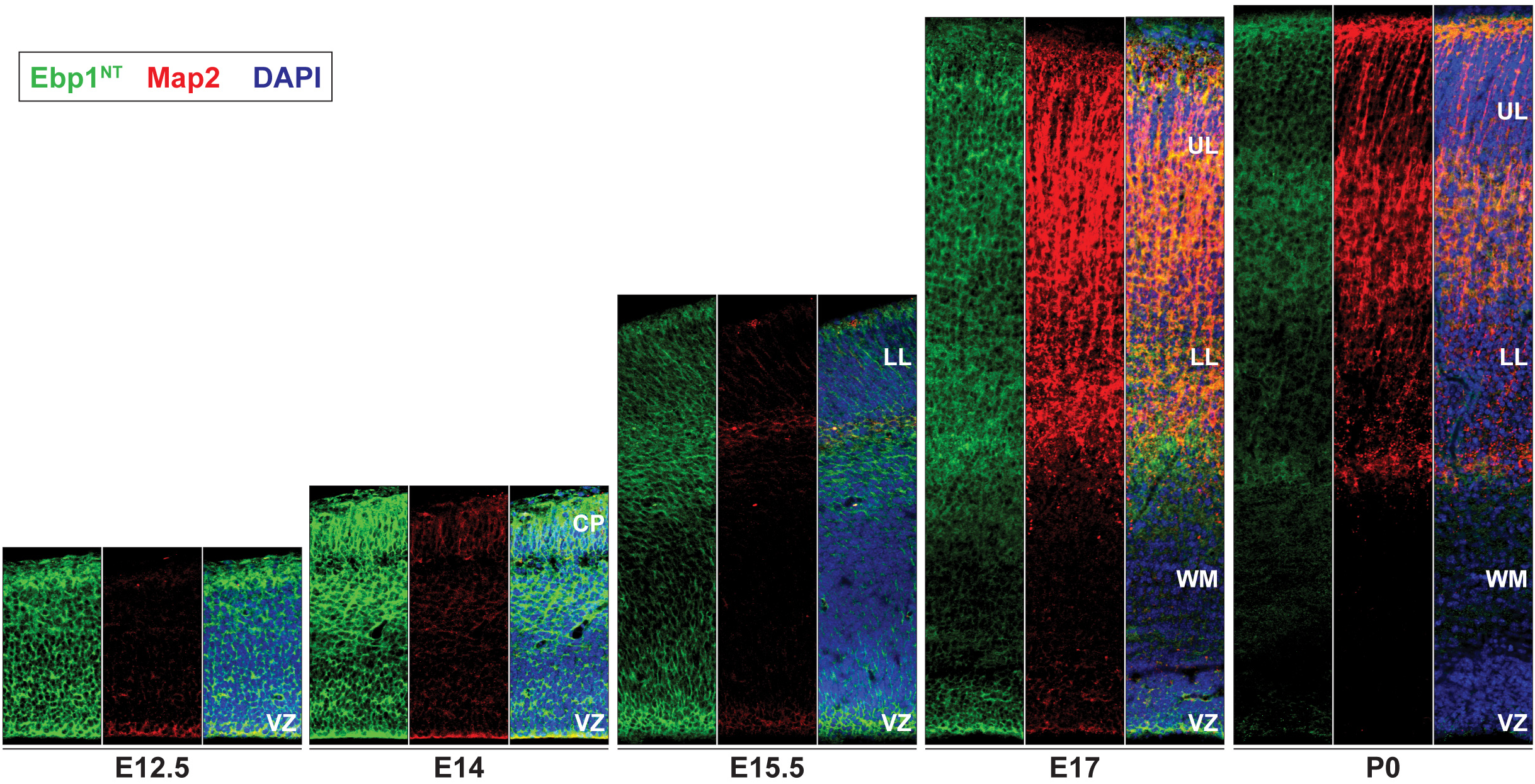
Ebp1^NT^ immunohistochemistry in the developing neocortex, associated with Figure 2C. Immunohistochemistry analysis of Ebp1 expression in coronal sections of the developing neocortex with the Ebp1^NT^ antibody (green), reinforcing findings with the Ebp1^CT^ antibody. Ebp1 levels are particularly high in early-born NSCs in ventricular zone (VZ), and nascent cortical plate (CP), at E12.5-E14. Ebp1 is expressed in the lower layers (LL) and upper layers (UL) of the expanding cortical plate at E15.5-P0, albeit at decreased levels. Co-immunostaining with Map2 (red) as a marker of maturing neurons in the CP, along with DAPI (blue) marking nuclei.

**Supplementary Figure 5.**
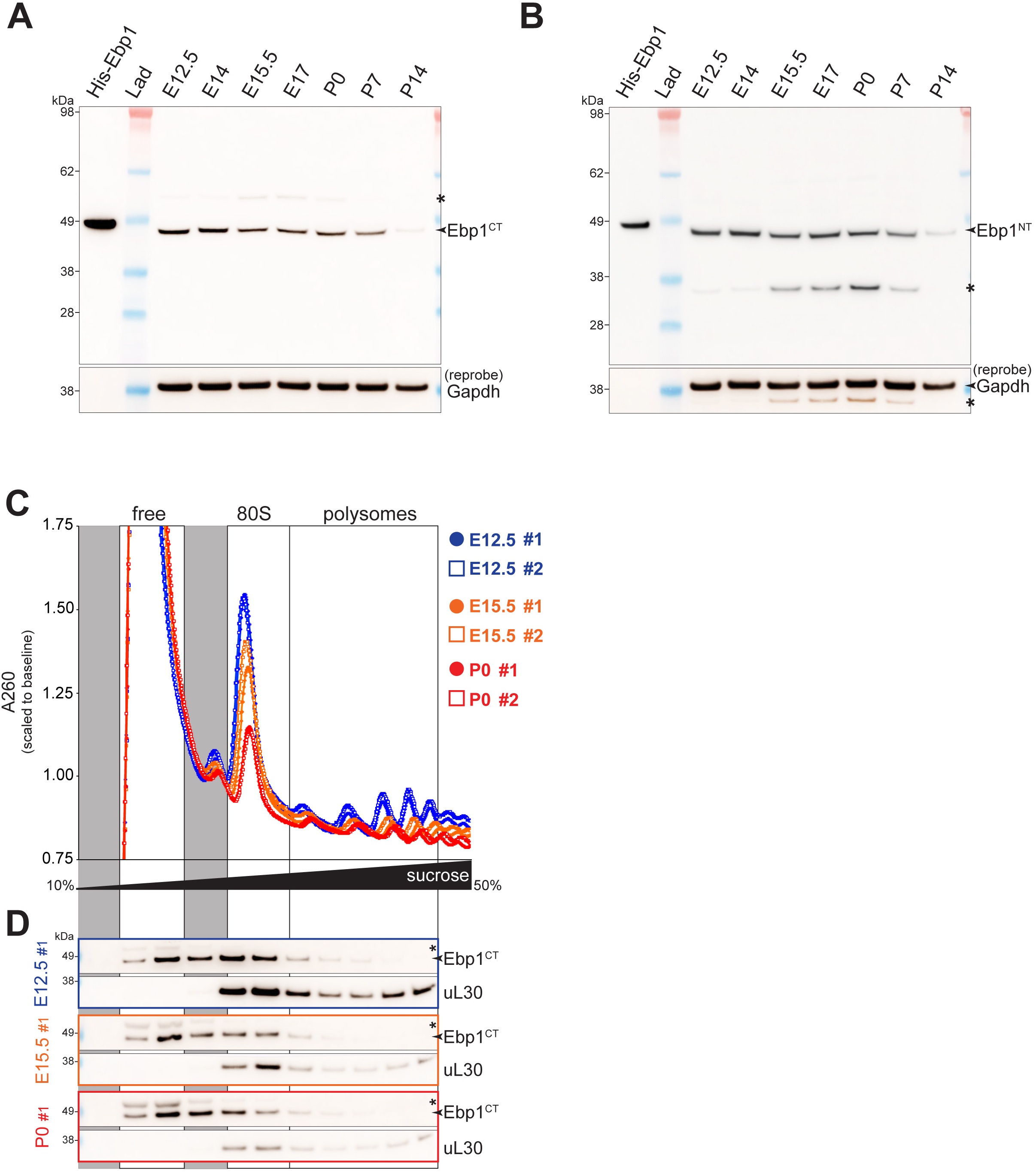
Full Western blots associated with Figure 2F, and sucrose density gradient fractionations and Western blots of each fraction, associated with Figure 2G. Western blot membranes of total neocortical lysates from E12.5, E14, E15.5, E17, and P0 probed with **(A)** C-terminal specific (Ebp1^CT^, membrane from Figure 2A; Abcam #ab35424) and **(B)** N-terminal specific (Ebp1^NT^, separate replicate; Millipore #ABE43) (Xia et al., 2001) anti-Ebp1, in comparison to full-length recombinant Ebp1 (Ebp1-His) as a band marker. Ebp1^CT^ antibody would be expected to identify both full-length Ebp1 (“p48”; ∼48kDa) and the N-terminal truncated isoform (“p42”; ∼42kDa), which lacks the first 54 amino acids (Liu et al., 2006). The Ebp1^NT^ antibody should exclusively identify full-length Ebp1. Results demonstrate dominant signal concordant with Ebp1 full-length at ∼48kDa (arrows), migrating slightly lower than recombinant Ebp1-His. Faint bands are seen at ∼55kDa and ∼35kDa (stars). Membranes were reprobed with Gapdh as a loading control. We concluded that the starred bands in (A) and (B) are both non-specific signal not corresponding to Ebp1, since these bands could not be replicated with both antibodies. Of note, all Ebp1 Western blots in this paper were performed with anti-light chain secondary antibodies, since application of anti-heavy chain secondaries was found to introduce strong artifact signal at ∼50kDa, creating an obstacle to the interpretation of the 48kDa Ebp1. **(C)** Total neocortical lysates at E12.5, E15.5, and P0 were A260 normalized, and fractionated by preparative sucrose density gradient ultracentrifugation into eleven fractions in biological duplicate, followed by **(D)** Western blot analysis of Ebp1 levels in each fraction corresponding to extra-ribosomal (free), 80S, and polysomes (replicate #1). Gapdh is a marker for pre-ribosomal free fractions, while uL30 is a marker for ribosome-associated fractions. While pre-ribosomal free Ebp1 levels are stable across neocortical development, ribosome-associated Ebp1 levels decrease in concert with decreasing levels of 80S and polysomes, seen in both the A260 curves (C) and likewise in uL30 protein levels (D). Individual fractions from replicate #2 were pooled to constitute free, 80S, and polysome samples in Figure 2G.

**Supplementary Figure 6.**
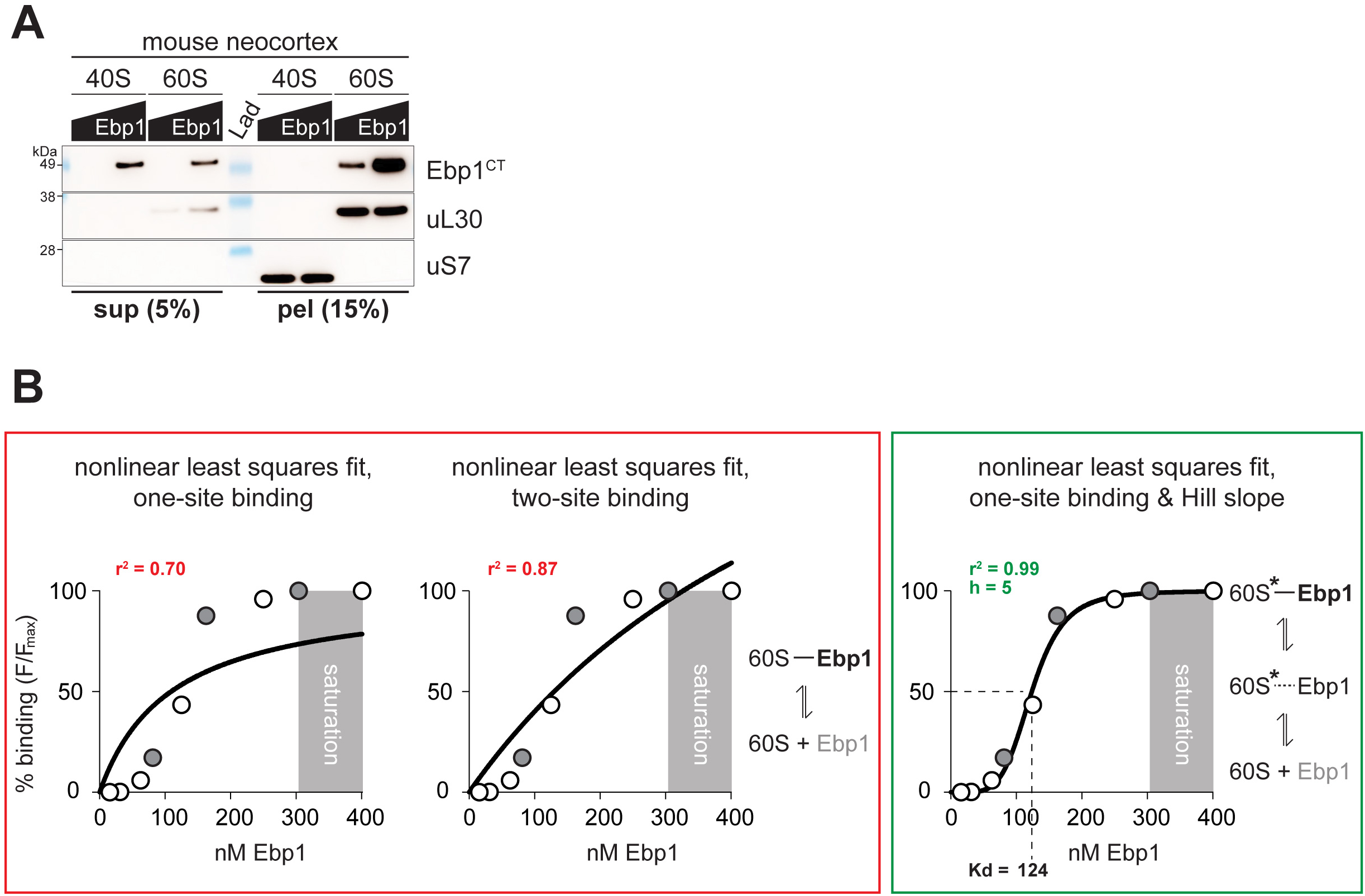
Biochemical analysis of Ebp1•ribosome binding, associated with Figure 3. **(A)** 40S and 60S subunits were purified from P0 mouse neocortex, and reconstituted with recombinant Ebp1-His, as in Figure 3A with rabbit-derived subunits. Escalating doses of Ebp1 (100nM, 200nM) were mixed with a constant 100nM of each subunit, and pelleted through a sucrose cushion to separate free unbound Ebp1 in the supernatant, vs. Ebp1•subunit complexes in the pellet. Western blot analysis demonstrates a dose-dependent binding of Ebp1 specifically to the 60S subunit. Marker for the 60S is uL30, and for 40S uS7. **(B)** Interpretation of binding data related to Figure 3C. Curves were fit to the data using a nonlinear least-squares fit, with one-site binding (left, r^2^=0.70), two-site binding (middle, r^2^=0.87), or one-site binding and Hill slope accommodation (right, r^2^=0.99). The best fit-to-data approximated by one-site binding and Hill slope accommodation (h=5) suggests that the 60S site serving as a receptor for Ebp1 binding acquires a conformational state (active state, 60S*) that is present with a higher probability when increasing concentrations of Ebp1 are present, yielding a higher probability of Ebp1 re-binding.

**Supplementary Figure 7.**
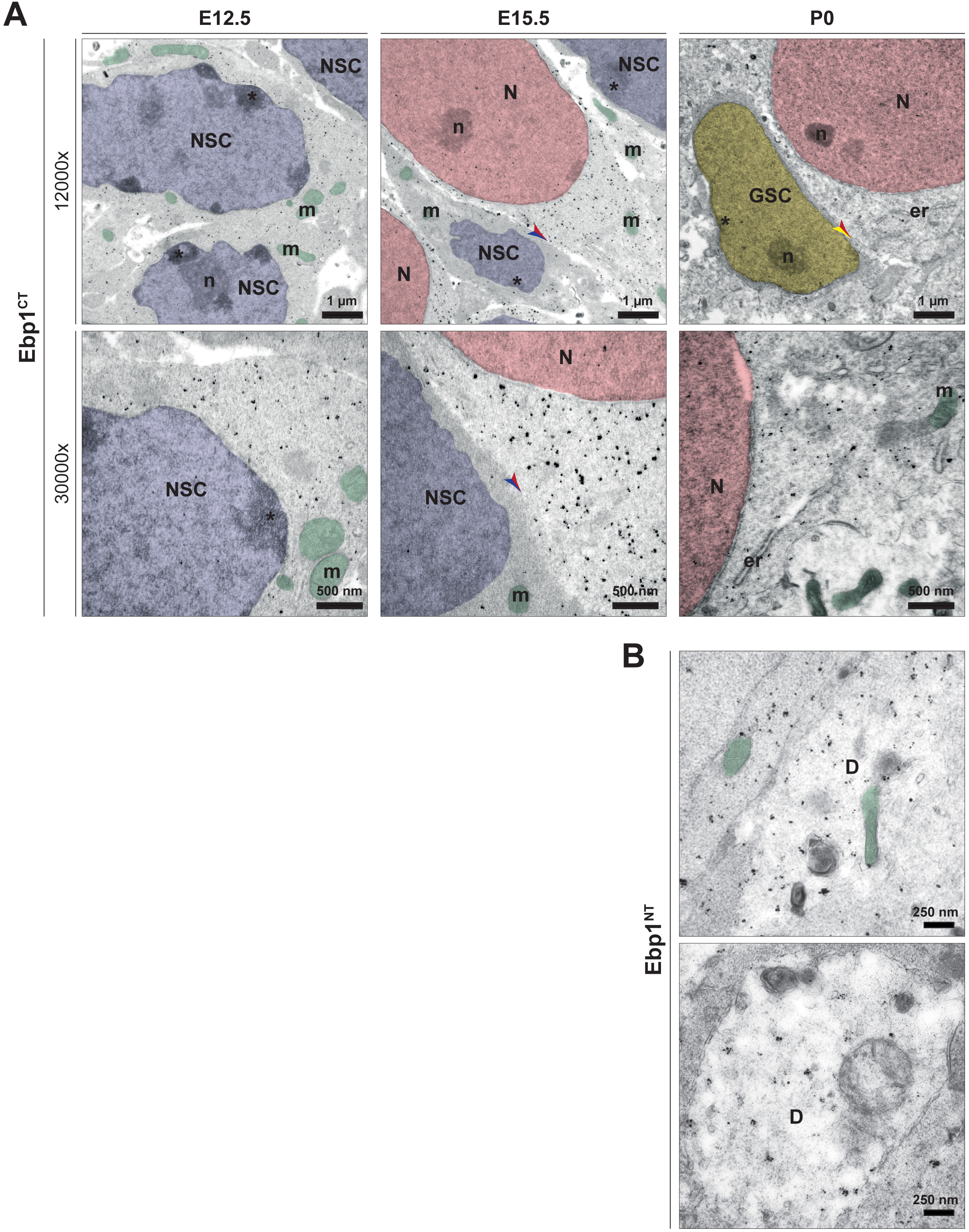
Immuno-electron microscopy with Ebp1^CT^ and Ebp1^NT^ antibodies, associated with Figure 3E. **(A)** Immunogold labeling for Ebp1^CT^ (black dots) in sections of the neocortex at E12.5, E15.5, and P0. Neural stem cells (NSC) and neurons (N) are identified by their distinctive nuclear morphologies (highlighted in blue, NSC and red, N) and positions in the developing cortical layers (ventricular zone, NSC; expanding cortical plate, N). Ebp1 labeling is predominantly found in clusters in the cytoplasm of these cells, but shows no association with the nucleolar site of RP synthesis (n), the endoplasmic reticulum (er), mitochondria (m, in green), or the plasma membranes at cell-cell junctions (arrows; blue on NSC side, red on N side). These results are in agreement with those obtained for Ebp1^NT^ in Figure 3E. **(B)** Electron micrographs of a dendritic branch (D) labeled for Ebp1^NT^ (black dots). Immunogold particles labeling Ebp1 are present in clusters in the dendritic cytoplasm, but not associated with mitochondria or the plasma membrane.

**Supplementary Figure 8.**
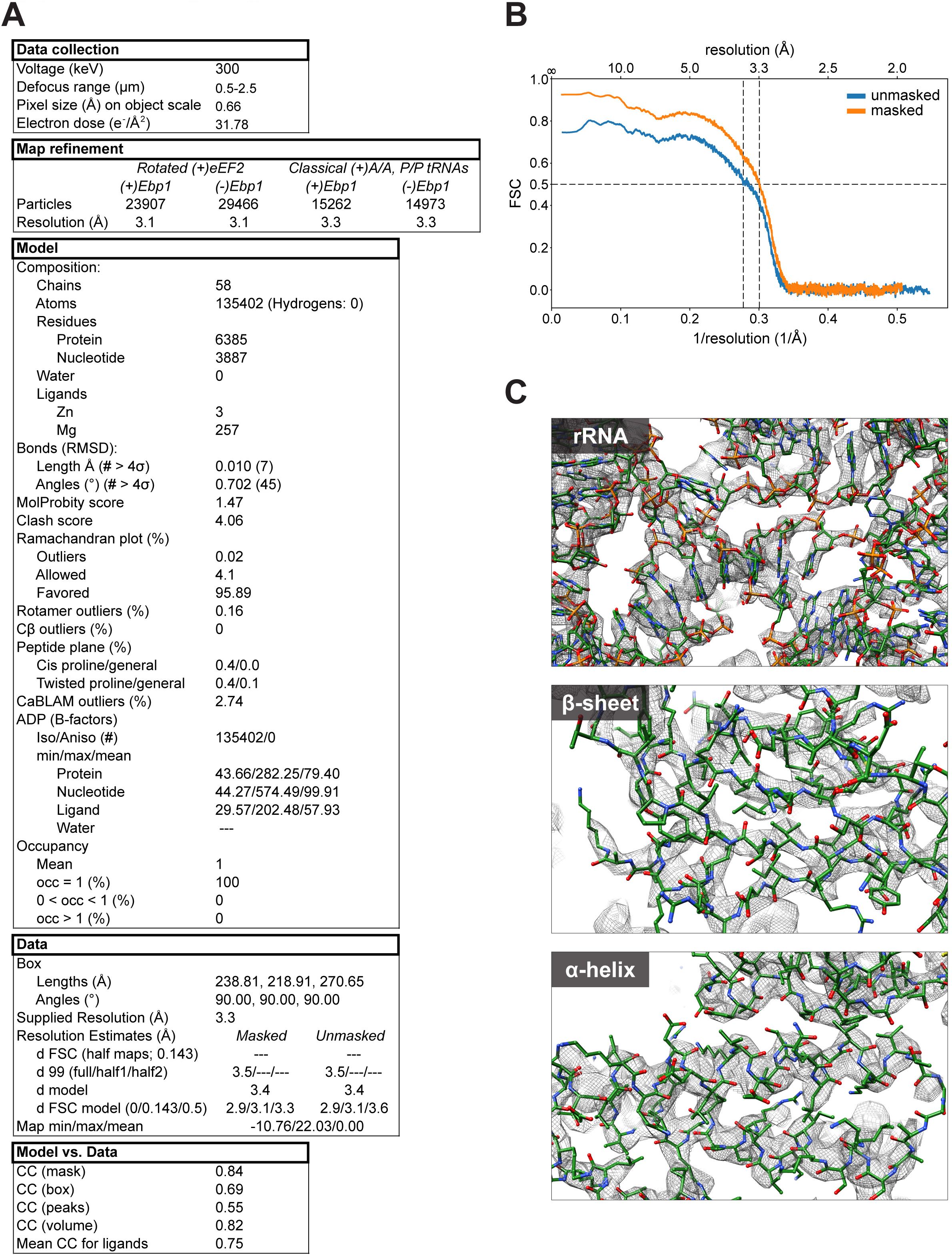
Cryo-EM data collection and model statistics, associated with Figures 4-5. **(A)** Statistics corresponding to cryo-EM data collection, map refinement, model characteristics, data resolution estimation, and cross correlation (CC) of model vs. data. **(B)** Fourier Shell Correlations (FSCs) for masked vs. unmasked maps. **(C)** Representative cryo-EM map (mesh) to model correspondence for rRNA, β-sheet, and α-helix structures.

**Supplementary Figure 9.**
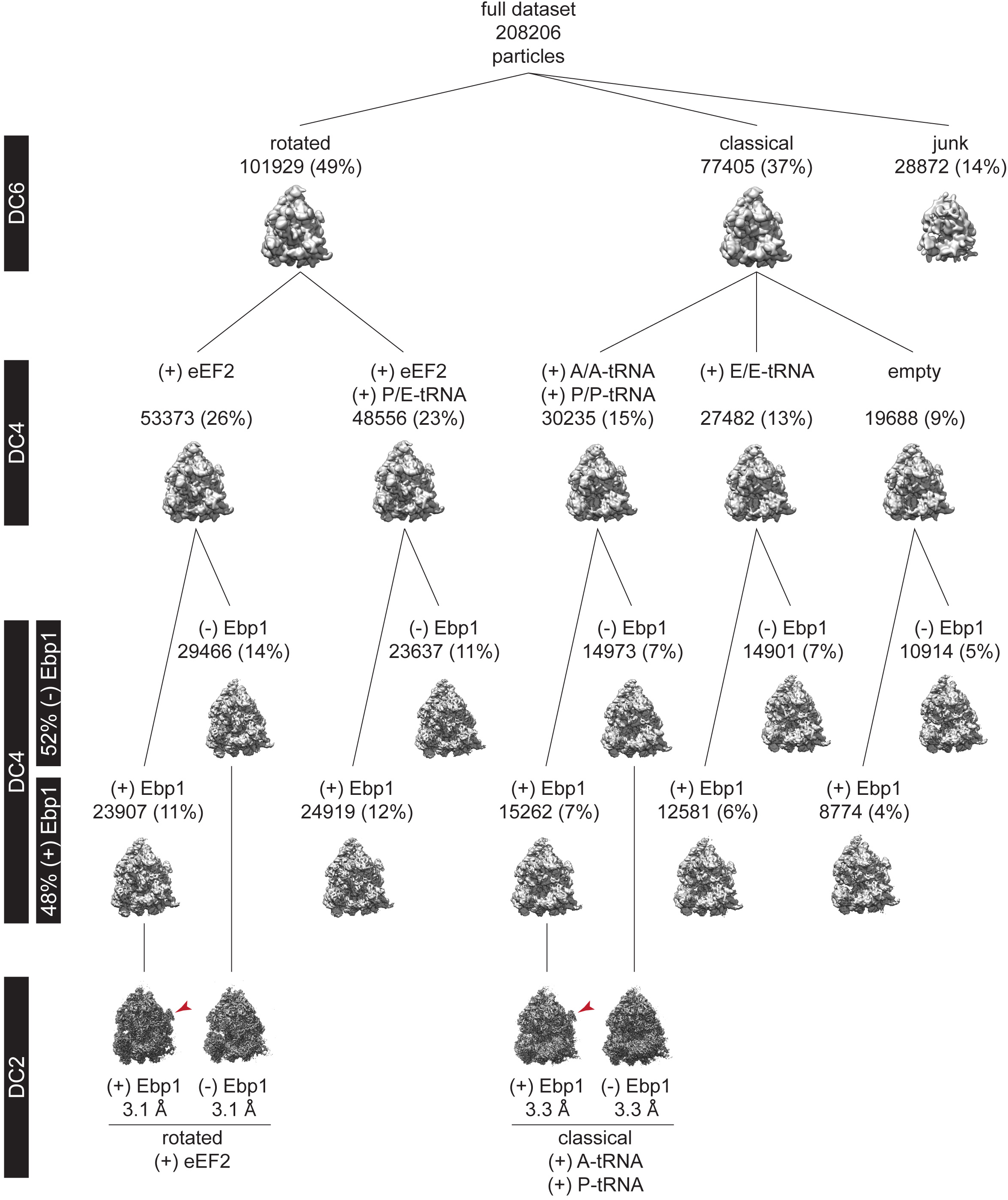
Multiparticle sorting and refinement of mouse neocortical ribosome states, associated with Figures 4-5. Cryo-EM imaging of pooled 80S and polysome complexes derived from the P0 mouse neocortex *ex vivo* yielded 208206 particles for *in silico* analysis. Initial multiparticle sorting of data at six times decimation (DC6) for large-scale heterogeneity generated ribosomes in the rotated and classical states, and a junk population. Further sorting within the rotated and classical states proceeded at DC4. In the rotated state, populations with eEF2 and eEF2+P/E tRNA emerged. In the classical state, populations with A/A+P/P tRNAs, E/E tRNA, and without tRNA emerged. In each of these five populations, extra-ribosomal density was observed at the peptide tunnel exit. Further sorting within each of these five populations proceeded at DC4 with a focus mask applied to this extra-ribosomal density as described in the Methods, to disentangle cofactor-positive and cofactor-negative ribosome sub-states. Each of these five populations yielded cofactor-positive and -negative populations. Final high-resolution refinement at DC2 proceeded for cofactor-positive (red arrow) and -negative populations of the rotated state with eEF2 (3.1Å global resolution), and the classical state with A/A+P/P tRNAs (3.3Å global resolution). These high-resolution data allowed for the identification of the cofactor as Ebp1, with cofactor-positive and -negative sub-states revealing 60S structural changes with Ebp1 binding.

**Supplementary Figure 10.**
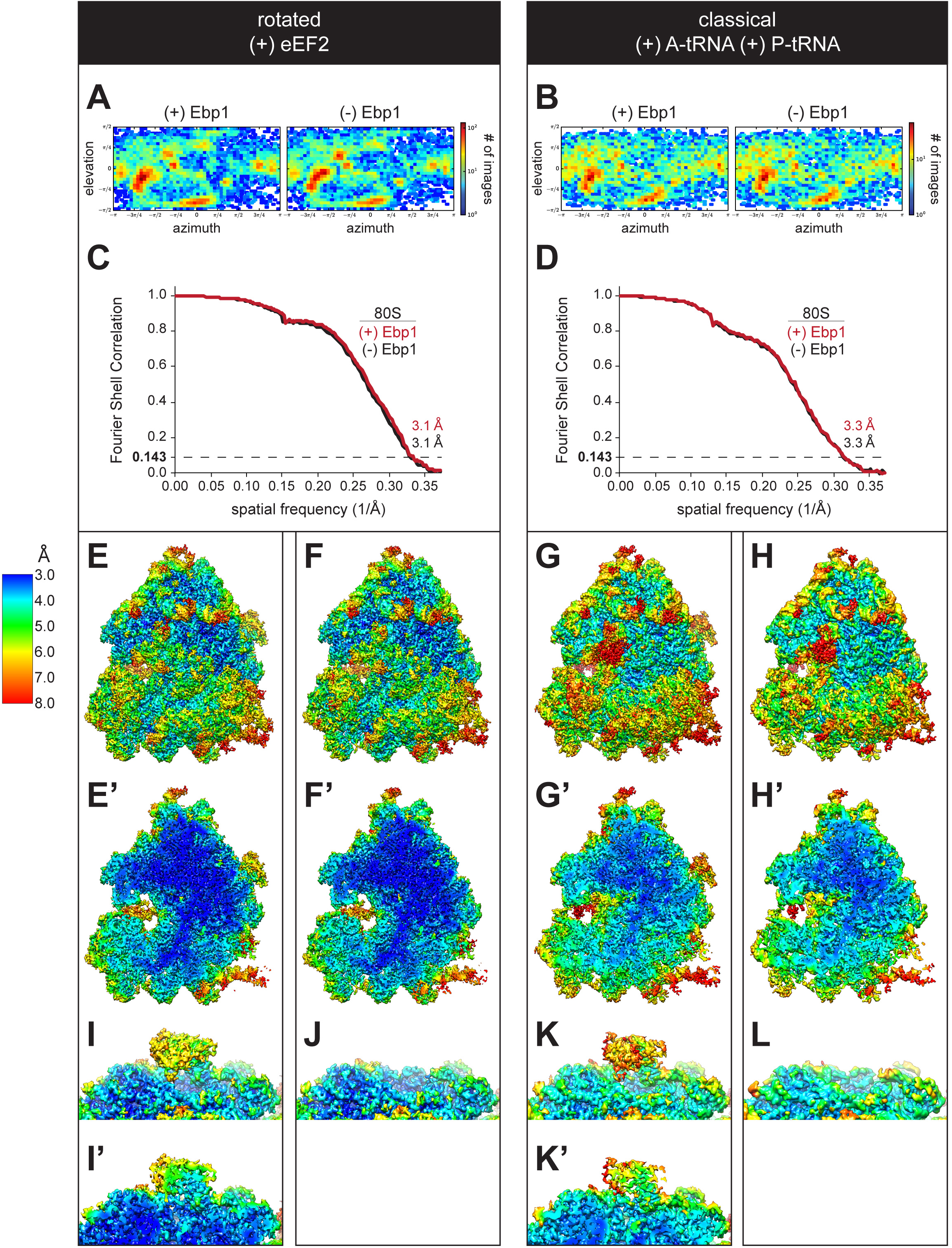
Global and local resolution measurements of cryo-EM maps, associated with Figures 4-5. Particle orientation distribution and global resolution Fourier Shell Correlation (FSC) for the **(A, C)** rotated state with eEF2, and **(B, D)** classical state with A/A+P/P tRNAs, both with and without Ebp1. Local resolution heat maps for the **(E, E’, F, F’, I, I’, J)** rotated state with eEF2, and **(G, G’, H, H’, K, K’, l)** classical state with A/A+P/P tRNAs, both with (E, E’, I, I’; G, G’, K, K’) and without (F, F’, J; H, H’, L) Ebp1. Maps are shown in both surface (E, F, G, H, I, J, K, L) and cross-section (E’, F’, G’, H’, I’, K’). The local resolution of Ebp1 is ∼4-6Å.

**Supplementary Figure 11.**
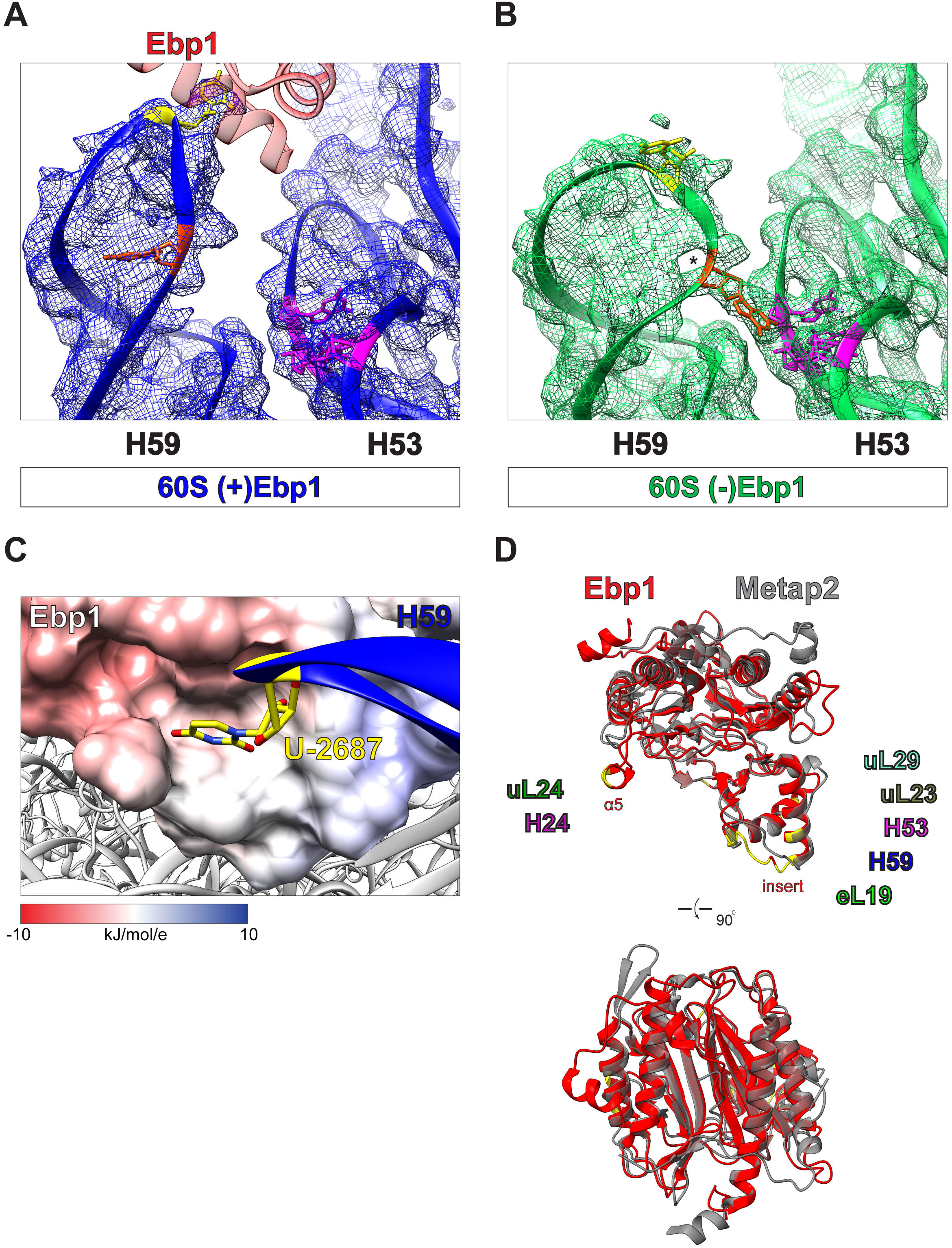
Cryo-EM density for H59 and H53 of Ebp1-positive and Ebp1-negative maps associated with the models shown in Figure 5A, electrostatics associated with Figure 5A, and Ebp1:Metap2 structure comparison associated with Figures 5C-D. Cryo-EM density (mesh) for the rotated state with eEF2 in **(A)** Ebp1-positive (blue) and **(B)** Ebp1-negative (green) sub-states. Density for base U-2687 (yellow) demonstrates flipping out when Ebp1 is bound, along with flipping in of base G-2690 (orange), and global reorganization of the H59 tip backbone away from H53. Base G-2690 in the Ebp1-negative state demonstrates density bridging to H53 bases G-2501, G-2502, and C-2513 (magenta) to stabilize H59’s canonical position, opening a gap (star) in H59 once occupied by the G-2690 intra-helical base stacking interactions. **(C)** Reorientation of H59 U-2687 into a pocket of Ebp1’s insert domain, facilitating Ebp1 stabilization on the 60S tunnel exit surface. Ebp1 shown as electrostatic potential map. **(D)** Global alignment of Ebp1 (red, PDB 2V6C) and Metap2 (grey, PDB 1KQ9), highlighting structural differences with Ebp1’s α5 domain, and overall structural similarity, such as the insert domain. Ebp1 residues making electrostatic interactions (yellow) with 60S TE rRNA helices and proteins are highlighted.

**Supplementary Figure 12.**
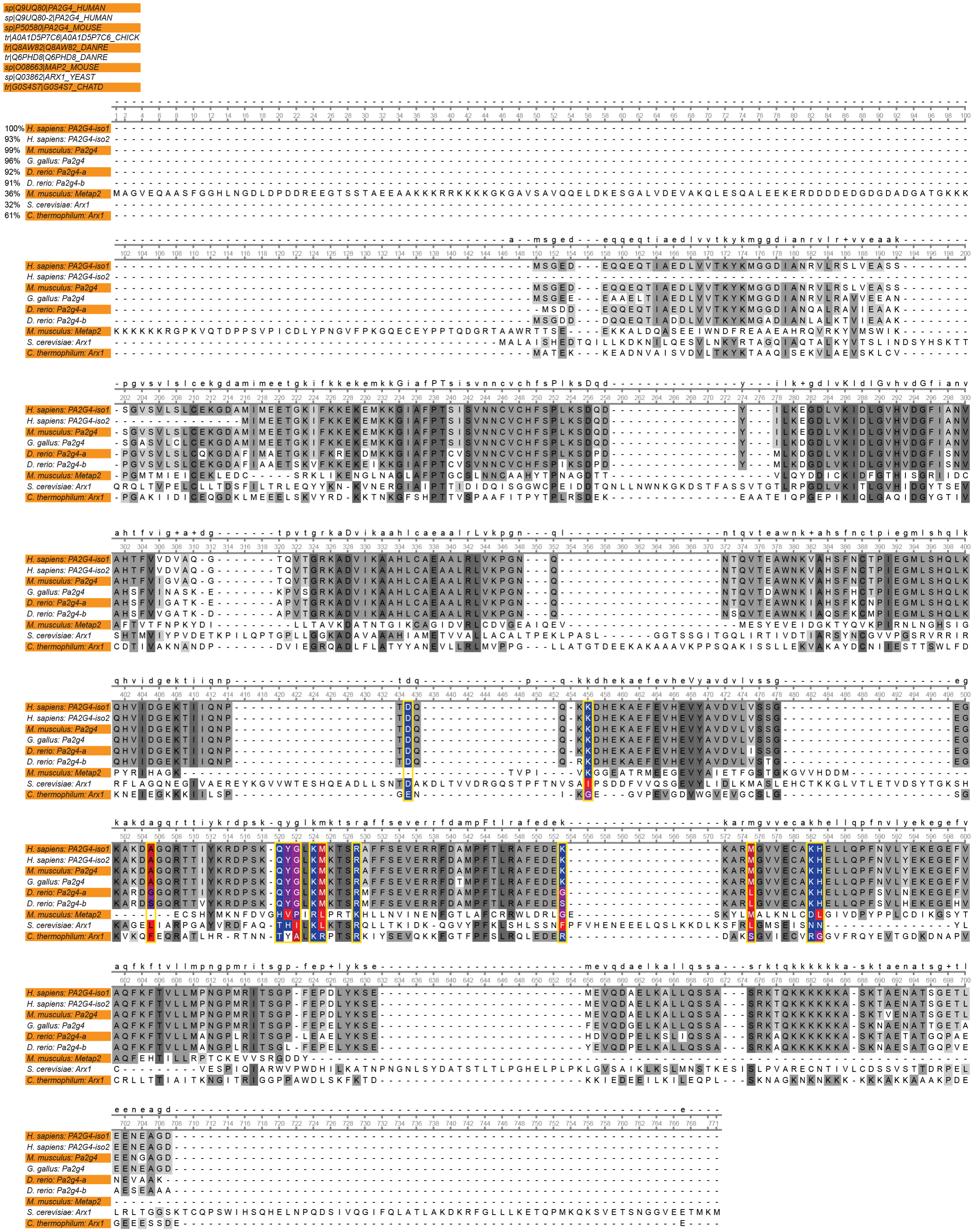
Sequence alignment of mouse Ebp1 with orthologs and structural homologs, associated with Figures 4-5. Ebp1 amino acid sequence alignment of human (*Homo sapiens* PA2G4 full-length, iso-1; N-terminal truncated, iso-2), mouse (*Mus musculus* Pa2g4 full length iso-1), chicken (*Gallus gallus* Pa2g4), and zebrafish (*Danio rerio* Pa2g4-a/b) orthologs with CLUSTAL Omega (1.2.4) (Sievers et al., 2011), default settings (https://www.ebi.ac.uk/Tools/msa/clustalo/) and visualized in Unipro UGENE (Okonechnikov et al., 2012). Alignment with structurally homologous proteins *Mus musculus* Metap2, *Saccharomyces cerevisiae* Arx1, and *Chaetomium thermophilum* Arx1 is also shown. Ebp1 residues involved in 60S binding are outlined in yellow aligned with homologous residues, colored by hydrophobicity. All other residues colored by percentage identity (grey). Percent similarity to the human full-length sequence is shown at left. Amino acid sequences were obtained from UniProt (The UniProt Consortium, 2019) (https://www.uniprot.org/) with entry number and name for each protein shown at top left.

**Supplementary Figure 13.**
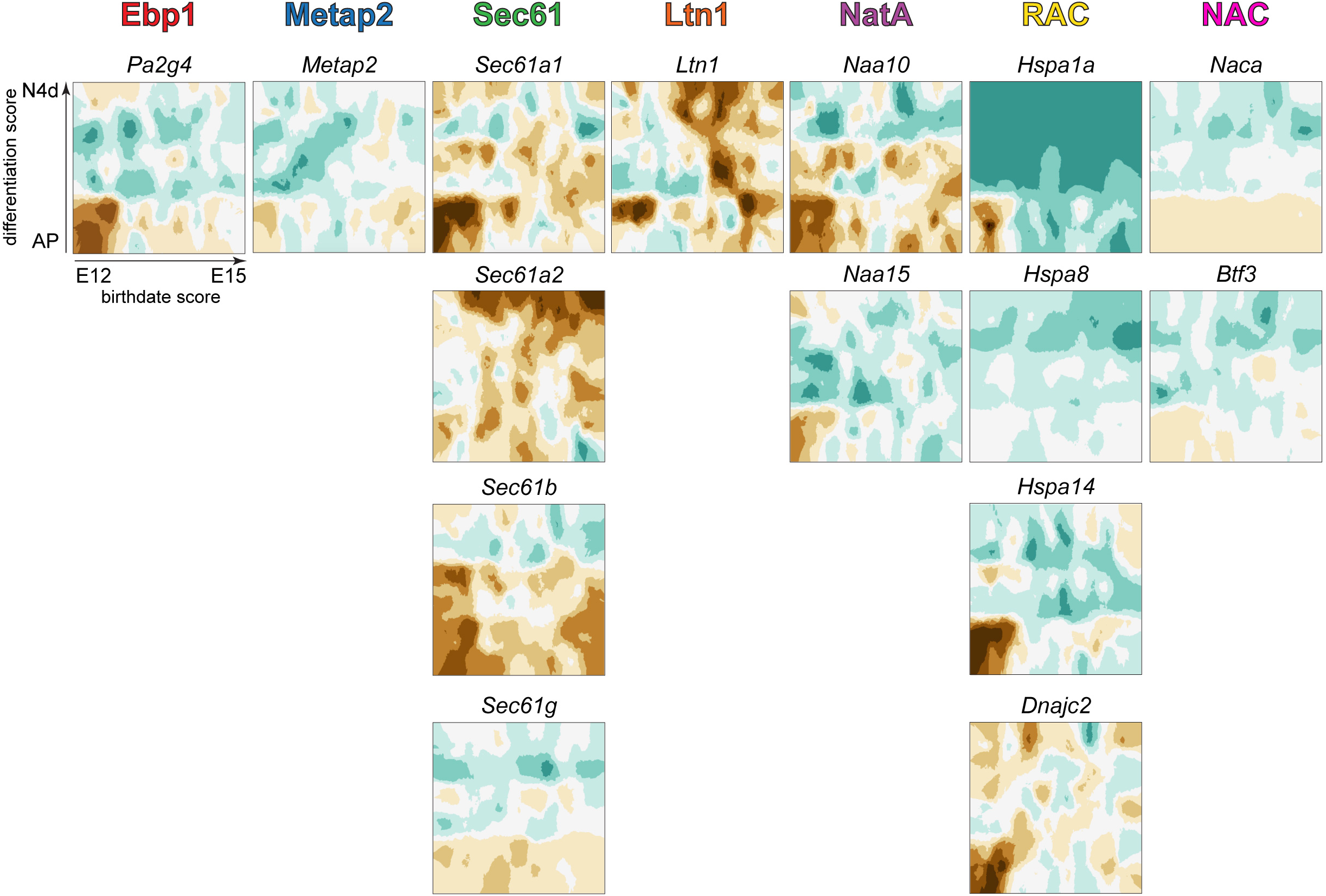
Transcriptional landscape of 60S tunnel exit binding cofactor expression in the developing neocortex derived from (Telley et al., 2019), associated with Figure 5E. Expression heat maps from scRNAseq data plotting NSC birthdate (x-axis) and differentiation (y-axis) scores, for the relative enrichment of mRNAs coding for 60S tunnel exit cofactors in the developing neocortex (ribosome-binding subunits shown). In each graph, the early-born apical progenitor (AP) NSC pool is plotted in the bottom left, with their corresponding differentiated lower layer neurons (N4d) in the top left. Late-born NSCs are plotted in the bottom right, with their corresponding differentiated upper layer neurons in the top right. Data from: http://genebrowser.unige.ch/telagirdon/#query_the_atlas

**Supplementary Figure 14.**
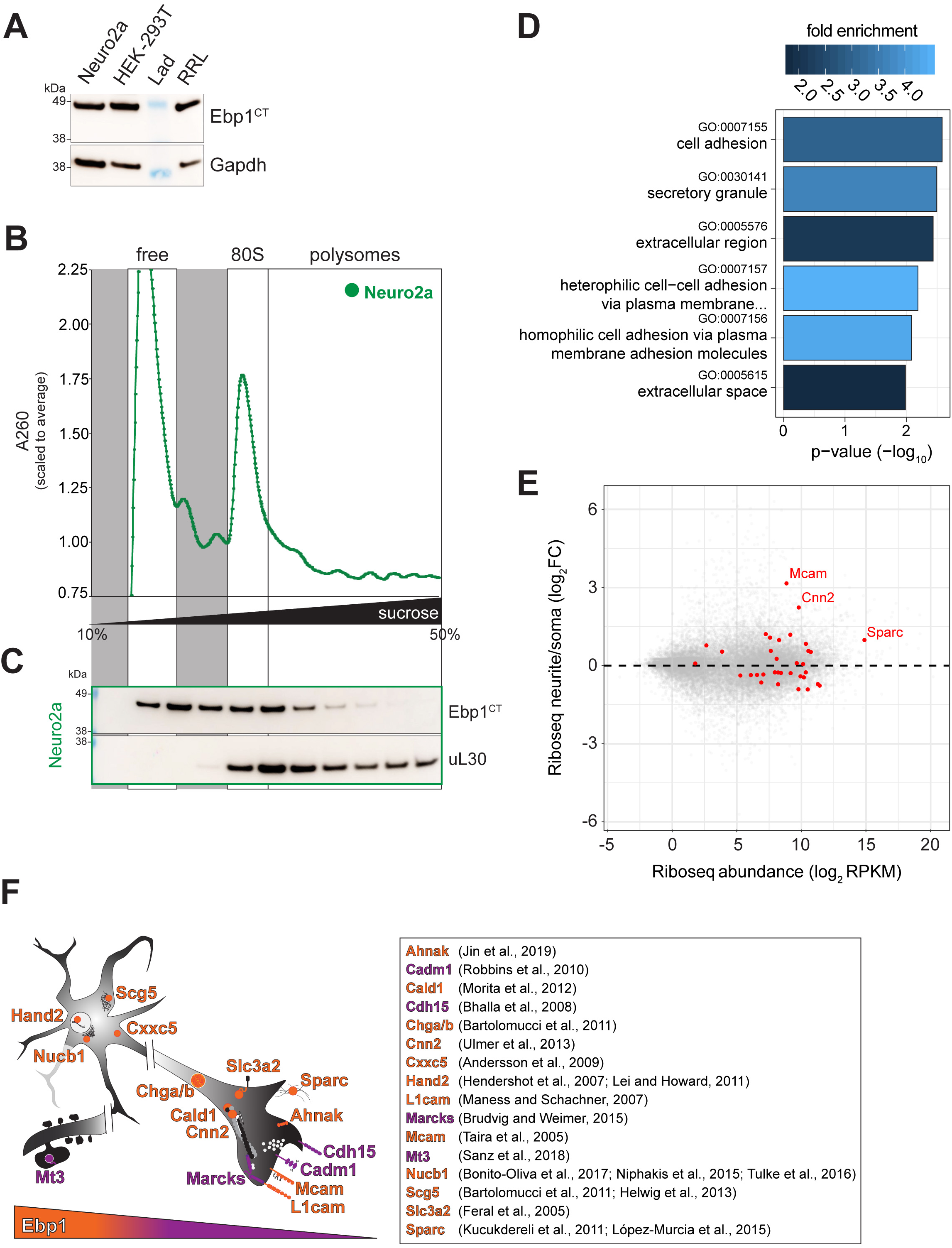
Ebp1 association with Neuro2a 80S and polysomes, and analysis associated with Figure 6. **(A)** Western blot of Neuro2a, HEK-293T, and rabbit reticulocyte (RRL) total lysates probed with Ebp1^CT^ antibody. **(B)** Preparative sucrose density gradient fractionation of Neuro2a cell lysates, followed by **(C)** Western blot analysis of Ebp1 enrichment of each fraction. Ebp1 associates with 80S and polysomes in Neuro2a cells, similar to mouse neocortical cells (**Figures S5C-D**). uL30 is a marker for ribosome-associated fractions. **(D)** Gene ontology (GO) pathways analysis with the Database for Annotation, Visualization and Integrated Discovery (DAVID) of proteins with >2-fold change from control in *siEbp1* conditions (against all quantified proteins) measured by pSILAC/AHA MS (**Figures 6C-D**). **(E)** Ribosome Profiling (Riboseq) derived from (Zappulo et al., 2017), highlighting the relative enrichment of Ebp1-regulated targets (red) in soma vs. neurites (y-axis) protein synthesis compared overall RPKM (x-axis) of cultured neurons. **(F)** Schematic summary of Ebp1’s impact on neuronal proteostasis, associated with **Figures 6C-D**, showing proteins that have been previously reported to influence neurogenesis. Ebp1 impacts cell adhesion molecules (de Wit and Ghosh, 2016), such as L1cam (Maness and Schachner, 2007), Mcam (Taira et al., 2005), Cadm1 (Robbins et al., 2010), and Cdh15 (Bhalla et al., 2008), in addition to the CAM modulator Slc3a2 (Feral et al., 2005). Ebp1 further regulates the synaptogenesis protein Sparc (Kucukdereli et al., 2011; López-Murcia et al., 2015), a membrane scaffold for calcium channels Ahnak (Jin et al., 2019), the neuronal migration and neurite outgrowth protein Marcks (Brudvig and Weimer, 2015), and the synaptogenic membrane-type metalloproteinase Mt3 (Sanz et al., 2018). Several of these proteins, in addition to Cald1 (Morita et al., 2012) and Cnn2 (Ulmer et al., 2013), interact with the cytoskeleton in growing neurites. Among the proteins modulating synaptic transmission that are sensitive to Ebp1 knockdown are chromogranins a/b (Chga/b) and the granin Scg5, that regulate the secretion of neuronal hormones, neurotransmitters, and growth factors (Bartolomucci et al., 2011). Ebp1 may further influence proteostasis through its regulation of protein aggregation modulators Nucb1 (Bonito-Oliva et al., 2017; Niphakis et al., 2015; Tulke et al., 2016) and Scg5 (Bartolomucci et al., 2011; Helwig et al., 2013). The impact of Ebp1 on the developmental neurogenic proteome is apparent in many of the proteins above, in addition to regulators of NSC signaling, such as Cxxc5 (Andersson et al., 2009) and Hand2 (Hendershot et al., 2007; Lei and Howard, 2011).

